# Spermatoproteasome-deficient mice are proficient in meiotic DNA repair but defective in meiotic exit

**DOI:** 10.1101/384354

**Authors:** Laura Gómez-H, Natalia Felipe-Medina, Yazmine B. Condezo, Rodrigo Garcia-Valiente, Isabel Ramos, Ignasi Roig, Manuel Sánchez-Martin, Dirk de Rooij, Elena Llano, Alberto M. Pendas

**Affiliations:** Instituto de Biología Molecular y Celular del Cáncer (CSIC-Universidad de Salamanca), 37007 Salamanca, Spain; Departamento de Fisiología y Farmacología, Universidad de Salamanca, 37007 Salamanca, Spain; Genome Integrity and Instability Group, Institut de Biotecnologia i Biomedicina, Universitat Autònoma de Barcelona, Cerdanyola del Vallès, Spain; Departamento de Medicina, Universidad de Salamanca, 37007 Salamanca, Spain; Reproductive Biology Group, Division of Developmental Biology, Department of Biology, Faculty of Science, Utrecht University, Utrecht 3584CM, The Netherlands

## Abstract

Meiotic recombination generates crossovers which are essential to ensure genome haploidization. The ubiquitin proteasome system regulates meiotic recombination through its association to the synaptonemal complex, a ‘zipper’-like structure that holds homologs and provides the structural framework for meiotic recombination. Here we show that the testis-specific α4s subunit (PSMA8) of the spermatoproteasome is located at the synaptonemal complex and is essential for the assembly of its activator PA200. Accordingly, synapsis-deficient mice show delocalization of PSMA8 from the synaptonemal complex. Genetic analysis of *Psma8*-deficient mice shows normal meiotic DNA repair, crossing over formation and an increase of spermatocytes at metaphase I and metaphase II which either enter into apoptosis or slip to give rise to an early spermatid arrest and infertility. Thus, spermatoproteasome-dependent histone degradation is dispensable for meiotic recombination. We show that PSMA8 deficiency alters the proteostasis of several key meiotic players such as acetylated histones, SYCP3, SYCP1, CDK1 and TRIP13 which in turn leads to an aberrant meiotic exit and early spermatid arrest prior to the histone displacement process that take place subsequently.

## Introduction

Intracellular protein contents are controlled through their rates of synthesis and degradation. In the cytosol and nucleus of eukaryotic cells, most of this degradation is carried out by the ubiquitin-proteasome system (UPS). The proteasome is a highly conserved protease complex that eliminates proteins typically labeled with ubiquitin by ATP-driven proteolysis (Collins & Goldberg, 2017). All the proteasome complexes have a cylindrical catalytic core particle (CP) and different numbers of regulatory particles (RP) that regulate the access to the CP by capping it at either end (Schmidt et al, 2005). Based on its sedimentation coefficients, the 26S proteasome is formed by one CP (20S) and one or two RP (19S). The CP is composed of seven α-type subunits (α1-α) and seven β-type subunits (β1-β7, three of which are catalytically active) arranged as a cylinder of four heptameric rings (α1-7, β1-7, β1-7, α1-7) (Collins & Goldberg, 2017; Murata et al, 2009). The two central β rings contain the proteolytic activity sites within a central chamber and the two alpha rings serve as a gate for substrate entry (Murata et al, 2009).

The 19S proteasome activator or RP is composed of 20 subunits and its association with the CP is ATP-dependent. However, there are four additional proteasome activators, the 11S regulator PA28α/β/γ (PSME1, PSME2 and PSME3) and the PA200 (PSME4) regulator that stimulate protein degradation by binding to the CP without ATP hydrolysis and independently of ubiquitin (Finley, 2009). Most proteasomes are comprised of one or two RP in combination with a CP but hybrid proteasomes enclosing a RP in one end of the CP and an activator on the other are also possible (Cascio et al, 2001).

This heterogeneity is further increased due to the existence of mammalian paralogs for the genes encoding the β1, β2 and β5 subunits which are expressed in the immunological system and constitutes the so called immunoproteasome (Griffin et al, 1998). In addition, there is a meiotic paralog of the α4 gene (*Psma7*) that encodes a protein with a very high identity to α4 named α4s which is encoded by *Psma8* (Uechi et al, 2014). The differences between α4s/PSMA8 and α4/PSMA7 are located in the outer surface of the CP, suggesting that it provides substrate specificity (Uechi et al, 2014). In addition, the ubiquitous subunit PA200 (highest expression in the male germline) is the main activator of the proteasome in the mammalian testis (spermatoproteasome) and play its main role in spermatogenesis. Genetic depletion of *Pa200* in the mouse abolishes acetylation-dependent degradation of somatic core histones during DNA double-strand breaks repair and delays core histone disappearance in elongated spermatids (Khor et al, 2006; Qian et al, 2013).

From a classical point of view, it is assumed that the rate of the protein ubiquitylation determines the rate of proteasome-dependent digestion (Collins & Goldberg, 2017). The resulting proteolytic products are short peptides most of which are digested by cytosolic peptidases to amino acids which are recycled by the cell for further protein synthesis (Kisselev et al, 1999). However, the proteolytic activity of the proteasome is also precisely regulated by the association of their CP and RP with E3 ubiquitin ligases and deubiquitinating enzymes (DUB) which edit their potential substrates (Inobe & Matouschek, 2014). The classical targets of the UPS are misfolded or damaged proteins with an abnormal conformations or short lived regulatory proteins, whose concentration is regulated by fine-tuning of their synthesis and degradation kinetics (Belle et al, 2006; Guo et al, 2016). Examples of the later proteins are cyclins in cell cycle control or the ZMM complex (also known as the synapsis initiation complex) in homologous recombination (Glotzer et al, 1991; Meyer & Rape, 2014; Rao et al, 2017). Functionally, the UPS is involved in a diverse array of biological processes, including protein quality control, apoptosis, immune response, cell-cycle regulation, DNA repair, and meiotic recombination.

Meiosis is a fundamental process in sexually reproducing species that generates new combinations between maternal and paternal genomes. During meiosis, two successive rounds of chromosome segregation follow a single round of replication, which leads to the production of haploid gametes from diploid progenitors (Zickler & Kleckner, 2015). This reduction in chromosome number is achieved by differences in chromosome dynamics during meiosis as compared with mitosis such as suppression of sister centromere separation (Kim et al, 2015) and also the physical connections between homologs by chiasmata. To generate these chiasmata the homologs; i) self-induce DSBs, ii) use as repair template the homolog non-sister chromatid, iii) find each other in the nucleus (pairing), iv) intimately associate (synapse) their proteinaceous core structures or axial elements (AEs) that scaffold the chromosomal DNA content, and, finally, v) connect physically through the assembly of the synaptonemal complex (SC) which provides the structural framework for all these processes to take place (Baudat et al, 2013). In yeast and mouse, the UPS regulates meiotic recombination in meiocytes via its physical association to the AEs (Ahuja et al, 2017; Rao et al, 2017). Through the RNF212 (E3 sumo ligase)-Hei10 (E3 ubiquitin ligase) pathway, the UPS regulates the proteostatic turnover of the ZMM (Shinohara et al, 2008), which is required for efficient synapsis and crossover (Rao et al, 2017).

Given our interest in meiotic recombination and fertility (Caburet et al, 2014; Gomez et al, 2016; Llano et al, 2014) and bearing in mind the existence of a specific proteasome (i.e. spermatoproteasome) in the male germline and the pivotal role that the USP plays in meiosis, we now have explored the functional analysis of the spermatoproteasome *in vivo* through the PSMA8 subunit of the mouse. We show that PSMA8 is located at the SC and is essential for the assembly of its activator PA200. Our results show that synapsis-deficient mice show a delocalization of PSMA8 from the SC and that male mice lacking PSMA8 (and thus also PA200) show normal meiotic DNA repair despite of abnormal acetylation-dependent degradation of histones. In addition, mutant male mice show an increase of spermatocytes at metaphase I and metaphase II which either enter into apoptosis or escape to give rise to fully arrested early round spermatids leading to male infertility. Finally, we show that PSMA8 deficiency provokes an alteration of the proteostasis of several key meiotic players such as SYCP3, SYCP1, CDK1 and TRIP 13 leading to an aberrant meiotic exit and deficient spermiogenesis with an early arrest in the development of round spermatid which is apparently unrelated with the classical histone displacement taking place later at mid round spermatid.

## Results

### Alpha4S/*Psma8* is expressed in spermatocytes and decorates the LEs

Analysis of *Psma8* mRNA expression in mouse tissues by RT-qPCR (Fig 1A) revealed that it is almost exclusively transcribed in testis (in agreement with the GTEx database (Consortium, 2015) and previous studies (Uechi et al, 2014)). We examined the expression of PSMA8 by western blot analysis of testis extracts at various postnatal ages during the first wave of spermatogenesis, which progresses more synchronously than in adult mice. The expression of PSMA8 (using a specific antibody against PSMA8 C-term, see Fig 1A (Uechi et al, 2014)), started on P12 and increased from P14 to P20 peaking at P16 (late pachytene-early diplotene). We also used an anti-PSMA8-R2 antibody raised against the whole recombinant PSMA8 protein (Fig 1B and EV1) and showed the expression of both PSMA7 (already detected at P8 when meiosis has not started) and PSMA8 (Fig 1B). In addition, we also analysed testis cell lines (including spermatogonium GC1-spg, Leydig cell TM3, and Sertoli cell TM4) showing ubiquitous expression of PSMA7 and only positive PSMA8 expression in whole testis which indicate that PSMA8 expression is restricted to cells undergoing meiosis.

**Figure 1.**
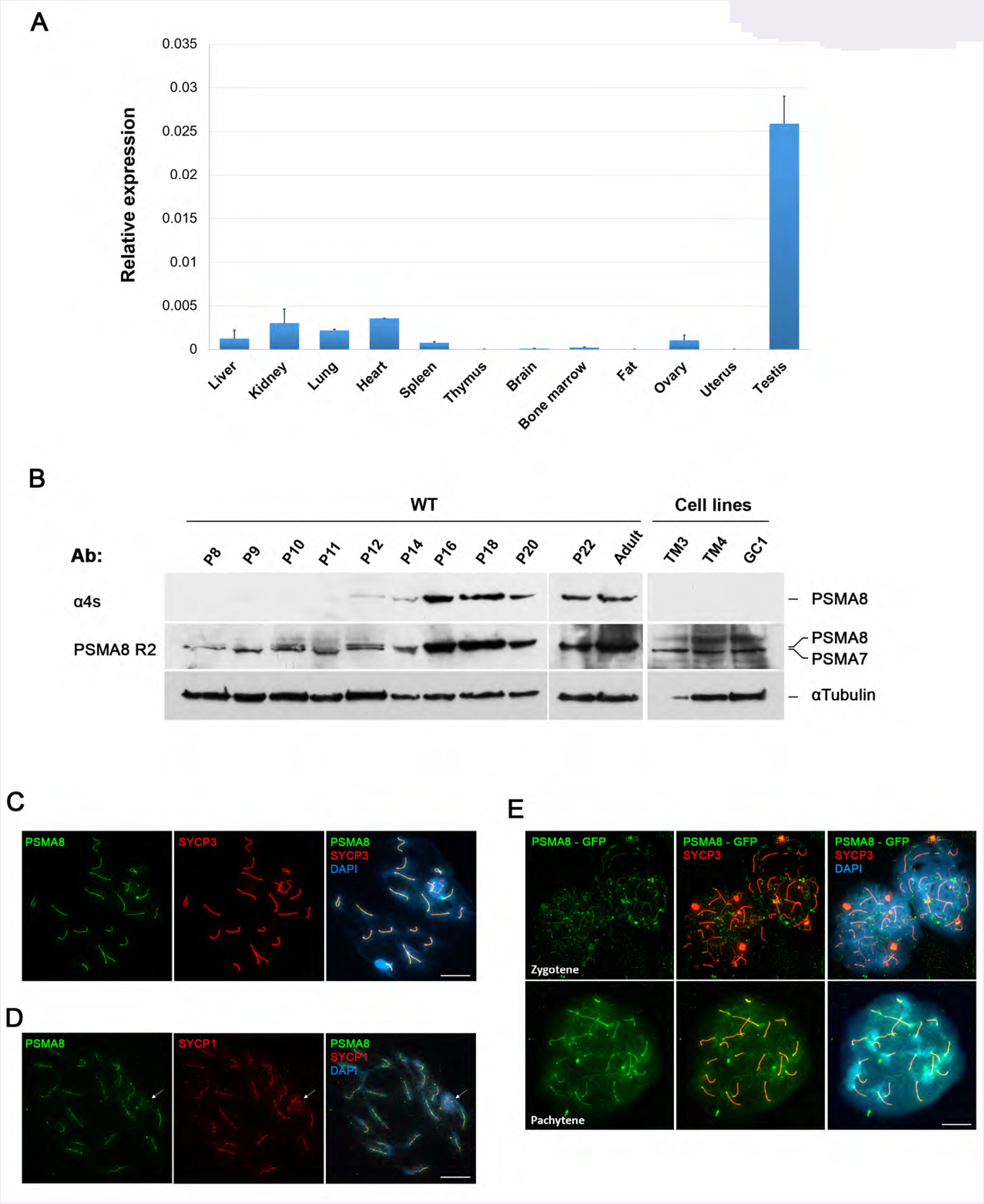
Transcriptional analysis and distribution of PSMA8 in mouse. (A) Relative transcription of *Psma8* mRNA by quantitative reverse transcription PCR (RT-qPCR) in mouse tissues. *β-actin* transcription was used to normalize the expression (mean ± s.d., three replicates). (B) Western blot analysis of protein extracts from wild type testis (from P8 to adult) and cell lines (TM3, TM4 and GC1) with a specific antibody against the C-term (α4S) and whole recombinant PSMA8 protein (PSMA8-R2). α-Tubulin was used as loading control. The corresponding bands to PSMA8 and PSMA7 are indicated in the right of the panel. Note that from P16 to adult the intensity of both PSMA8 and PSMA7 bands impedes its independent observation. (C) Double labelling of endogenous PSMA8 (green) and SYCP3 (red) in mouse spermatocytes. DNA was stained with DAPI (blue). During pachytene, PSMA8 is located at the synapsed autosomal LEs and at the PAR of the sex XY bivalent. (D) Double immunolabelling of spermatocytes spread preparations with PSMA8 (green) and SYCP1 (red), showing that PSMA8 localizes to the synapsed LEs but do not perfectly co-localize with SYCP1. PAR of the sex XY bivalent is indicated with an arrow. (E) Immuno-localization of PSMA8 in mouse testis after *in vivo* electroporation of a plasmid encoding a protein fusion of PSMA8 with GFP (GFP-PSMA8). PSMA8 was detected with anti-GFP antibody (green) and endogenous SYCP3 was detected using mouse anti-SYCP3 (red). DNA was stained with DAPI (blue). Bar in panels, 10 μm.

To explore the subcellular localization of PSMA8, we employed the R2 antibody (PSMA7/8). Double immunolabelling of PSMA8 with the AE protein SYCP3 and with SYCP1, the transverse protein essential for synapsis (Fig 1C-D and EV2), showed that PSMA7/8 (R2 antibody) was detected from zygotene (start synapsis) to pachytene along the fully synapsed LEs where SYCP1 is localized. On the XY bivalent only the pseudoautosomal region (PAR) is synapsed and this region labeled positive for PSMA7/8 (Fig 1D). As desynapsis progresses through diplotene, PSMA7/8 was only observed at the still synapsed regions (Fig EV2). The specific C-term antibody did not produce any labelling by IF, preventing its use for this application. In order to validate the localization of PSMA8 independently of PSMA7, we *in vivo* electroporated an expression plasmid encoding GFP-PSMA8 in the mouse testis (Gomez et al, 2016). After 48 hours, the GFP-PSMA8 colocalized with SYCP3 along the synapsed LEs at pachytene (Fig 1E), corroborating its localization to the chromosomal axes of mouse spermatocytes. This result agrees with the cytological localization of the proteasome to the LEs of the chromosomes (Rao et al, 2017). Altogether, our results indicate a specific meiotic function of the CP of the spermatoproteasome during mouse male meiosis.

### PSMA8 loading is dependent on synapsis

To investigate the possible dependence of PSMA8 localization on the degree of chromosome synapsis, we analysed spermatocytes of mice deficient for *Six6os1* (Gomez et al, 2016) and *Rec8* (Bannister et al, 2004). These meiotic mutants display different synaptic defects, from mild to more severe. Interestingly, in Rec8-deficient mice, where there is no true synapsis between homologues but instead in some pachynema-like cells there is “auto-synapsis” between sister chromatids (Xu et al, 2005), PSMA8 is also present at these atypical synapsed-like regions (Fig 2). However, in mice lacking the novel central element protein SIX6OS1, in which AEs completely fail to synapse in a pachytene-like stage with most homologs aligned but physically separated (Gomez et al, 2016), PSMA8 was not detected onto the LEs. Instead, a wide and disperse labelling appears decorating the chromatin of the axes (Fig 2). Together, our results indicate that PSMA8 localization to the CE is consequently dependent on the assembly of the SC either between the homologous chromosomes or the sister chromatids.

**Figure 2.**
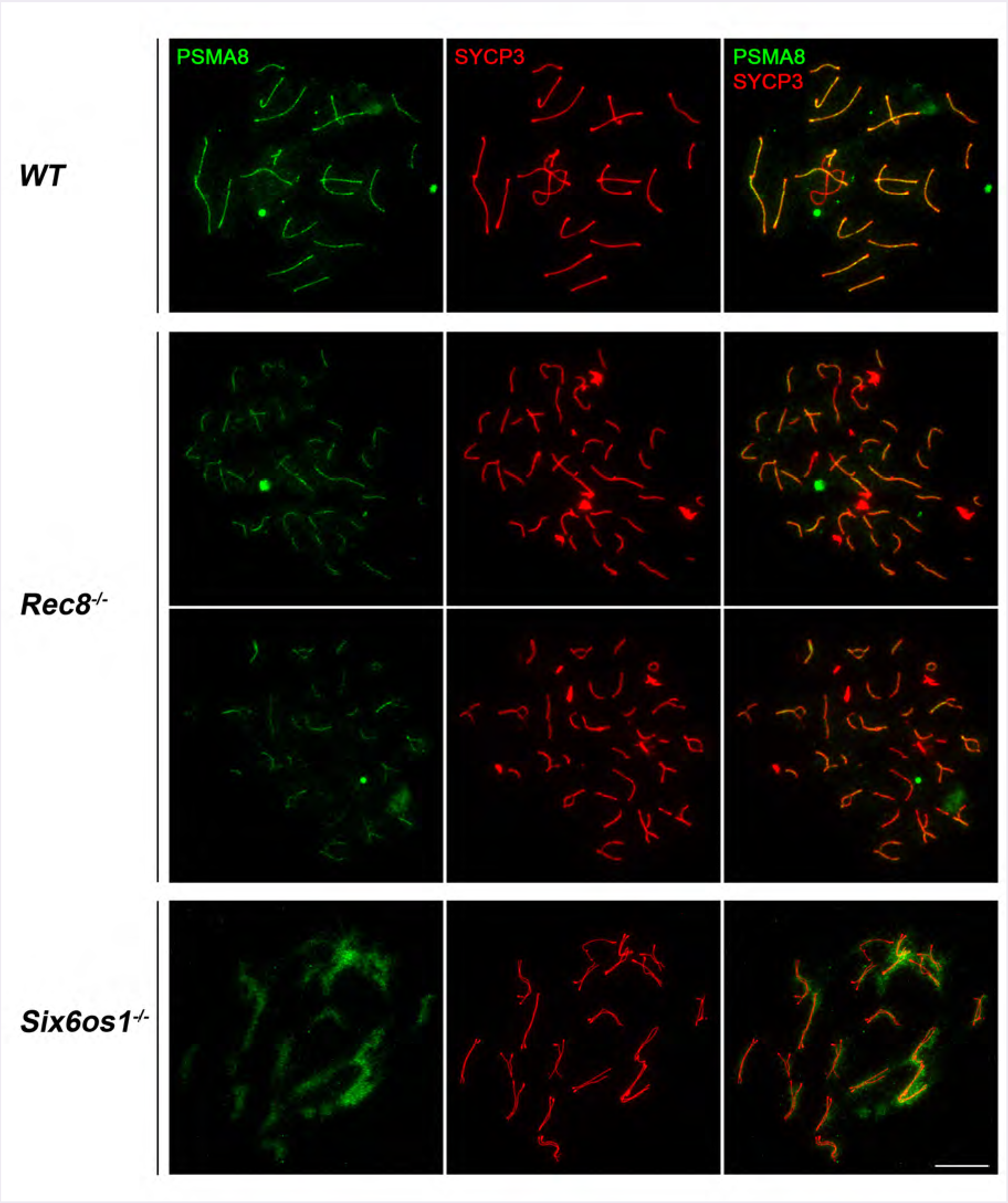
PSMA8 localization is dependent on synapsis. Double labelling of PSMA8 (green) and SYCP3 (red) in *Rec8*^−/−^ and *Six6os1*^−/−^. PSMA8 is detected in the pseudosynapsed AEs of the meiotic *Rec8* cohesin mutant but is absent from unsynapsed AEs in *Six6os1*^−/−^ spermatocytes. Bar in panels, 10 μm.

The interdependency of the transverse filament and central element proteins for the proper assembly of the SC is evidenced by the lack of loading of any of the central element subunits in the synapsis-less *Six6os1* or *Syce1* mutants (Bolcun-Filas et al, 2009; Gomez et al, 2016). Thus, we expect delocalization of the spermatoproteasome from the LEs in the mouse mutants of the central element proteins such as SYCP1, SYCE1, SYCE2, SYCE3 and TEX12 with unexplored functional consequences for their corresponding meiotic arrest.

### Purification of PSMA8 interacting proteins

The composition of the CPs and their RPs, has previously been shown by mass-spectrometry analysis of crude preparation of proteasomes from the whole testis (Bousquet-Dubouch et al, 2009). In order to gain further insight into the composition of the spermatoproteasome and specially their proteasome-interacting proteins, we purified PSMA8 interacting proteins by a single step affinity chromatography procedure using Sepharose-bound to the antibody R1, which detects endogenous PSMA8 (in addition to PSMA7) under the fixative conditions of the chromosome spreads. As a proof of concept the most prevalent protein detected was PSMA8 followed by PSMA7 (two orders of magnitude lower; Table EV1), suggesting a moderate specificity of the R1 antibody (for PSMA8 vs PSMA7) under more native conditions than in western blot and IF. Most of the canonical subunits of the CP and RP were present within the more than 596 proteins of the PSMA8 proteome (Table EV1, using a conservative cut off, see methods). In agreement with previous results, among the two activators of the spermatoproteasome detected (PA200 and pa28γ), PA200 was the most abundant (Qian et al, 2013). In contrast with previous observations, we were not able to detect neither pa28α and pa28β nor the inducible catalytic subunits of the immunoproteasome (P1i, P2i and P5i) (Qian et al, 2013). Of interest, we detected several novel potential candidate proteasome interacting proteins (PIPs) with high confidence (Table EV2). Among ubiquitin ligases and USPs, we detected known PIPs such as chaperones HSP90-β, HSP90AB, HSP70a2, and PAC1 (PSMC1), E3 ligases like SKP1, RBX, CAND1, and USP7, USP14 and USP47. The absence of known PIPs of the 20S proteasome such as UCH37 (UCHL5) and RPN13 (ADRM1) and RPN14 (PAAF1) is noteworthy. Among new PIPs we detected, were chaperones like CCT6b and CCT2, ubiquitin ligases like TRIP12, NEDD4, TRIM36 and RAD18 (USP7), and new USPs like USP9X, USP34, USP5 and USP47. Altogether, this list of novel proteins with the potential ability to edit the ubiquitin code of the interactome of the spermatoproteasome reflects the complexity of the spermatoproteasome system.

Interestingly, we identified proteins *a priori* unrelated to the UPS such as SYCP1, TRIP13 (pachytene checkpoint), CDK1 (cyclin dependent kinase 1), SMC6 or SMC4 (subunit of the condensin complexes). Furthermore, there were proteins with important meiotic functions or involved in spermatogenesis such as DAZL (deleted in azoospermia), SPAG1 (Sperm-associated antigen 1), SPATA5 / SPATA20 (Spermatogenesis-associated protein 5 / 20), the tudor domain proteins TDRD1, TDRD6 and TDRD9, MAEL (reppresor of transposable elements) and RNF17 (see below). These PIPs could represent proteins captured during the Ub-dependent targeted degradation as has been previously described (Verma et al, 2000) and/or proteins interacting via the Ubiquitin-independent proteasomal degradation (UIPD) as has been shown for other subunits of the CP and especially the related subunit α4/PSMA7 (Sanchez-Lanzas & Castano, 2014).

Finally, we studied the proteins enriched in the IP through its functional (gene ontology, GO) and pathway analysis (KEGG). The results of the top GO and KEGG overrepresentation test were related to the proteasome complex, proteasomal catabolic process and to ribonucleoproteins. Pathway analysis were linked to spermatogenesis, cell cycle, and meiosis (Appendix).

### Mice lacking PSMA8 are infertile

To study the role of PSMA8 we generated a targeted mutation in exon 1-intron1 of the murine *Psma8* gene by CRISPR/Cas9 genome editing (Fig 3A). The predicted best null mutation was selected by PCR sequencing of the targeted region of the *Psma8* gene (Fig 3B). The selected founder was crossed with wild-type (WT) C57BL/6J. Heterozygotes were crossed to yield homozygous mutant mice (knock-out, KO) which were identified by PCR (Fig 3B). Homozygous mutant mice showed no PSMA8 protein expression by western blot when analysed using two independent polyclonal antibodies raised against the whole recombinant PSMA8 protein (Figs 3C and EV1). When we analysed PSMA8 expression on mutant spermatocytes by IF (R2 antibody, Fig 3D) we obtained a weaker signal (51% less; 4.22±1,9 WT vs 2.05±1.7 KO) in the synapsed LEs of the spermatocytes in comparison with the wild type control, likely representing the PSMA7 detected by the R2 Ab (observed also in the WB; Fig 3C). These results indicate that the generated mutation is a null allele of the *Psma8* gene (herein *Psma8*^−/−^).

**Figure 3.**
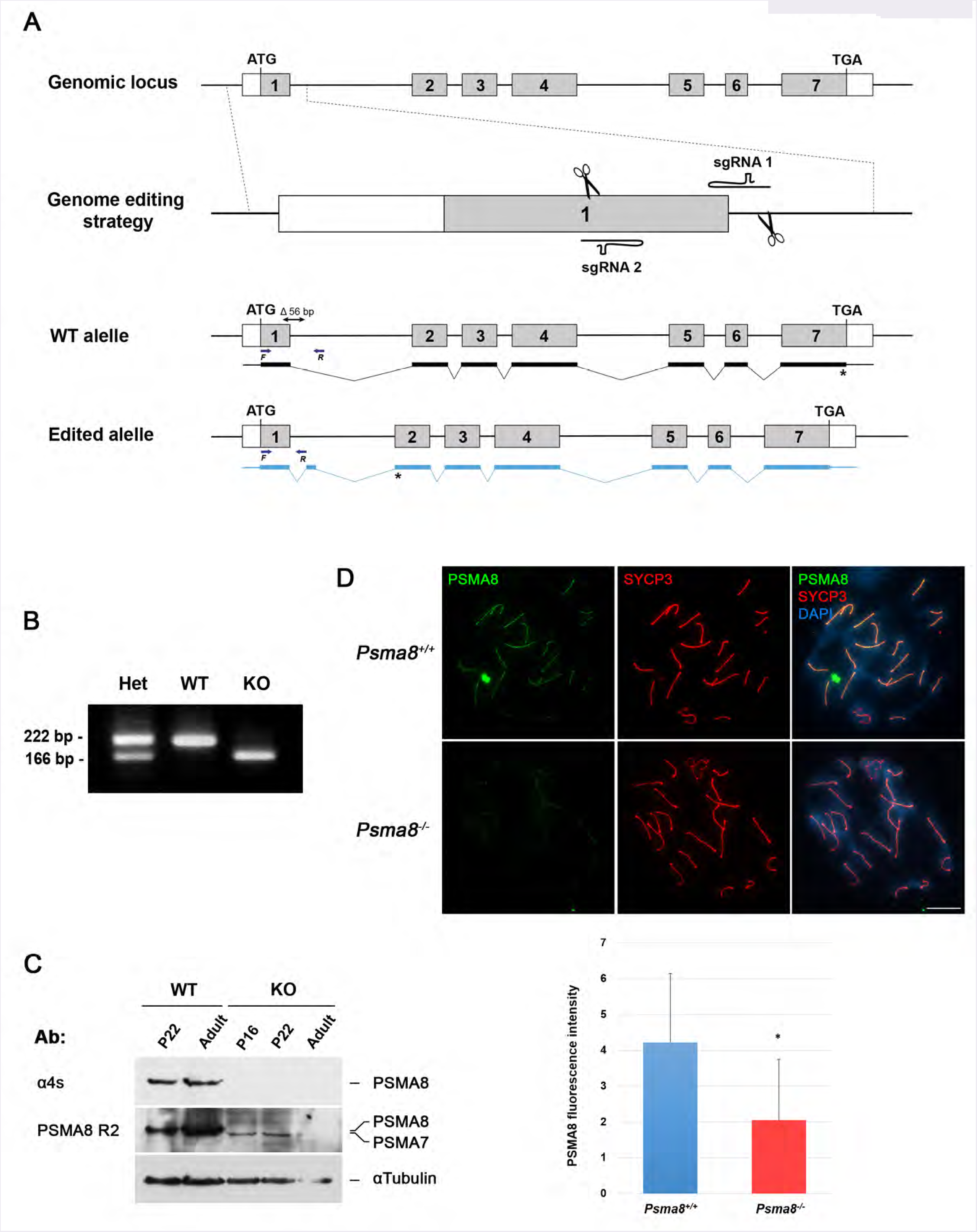
Generation and genetic characterization of *Psma8* deficient mice. (A) Diagrammatic representation of the mouse *Psma8* locus (WT) and the genome editing strategy showing the sgRNAs located on exon 1 and intron 1 (see methods), the corresponding coding exons (light grey) and non-coding exons (open boxes). Thin (noncoding) and thick (coding sequences) lines under exons represent the expected transcript derived from wild-type (black) and *Psma8* edited allele (blue). ATG, initiation codon; TGA, stop codon. The nucleotide sequence of the 56 base pair deletion derived from PCR amplification of DNA from the *Psma8^edlted/edlted^* is indicated (A). Primers (F and R) are represented by arrows. (B) PCR analysis of genomic DNA from three littermate progeny of *Psma8*^#/−^ heterozygote crosses. The PCR amplification with primers F and R revealed 222 and 166 bp fragments for wild-type and disrupted alleles respectively. Wild-type (WT, +/+), heterozygous (Het, +/-), and homozygous knock-out (KO, -/-) animals. (C) Western blot analysis of protein extracts from wild type testis (P22 and adult), KO testis (P16, P22 and adult) with a specific antibody against the C-term (α4S) and whole recombinant PSMA8 protein (PSMA8-R2). α-Tubulin was used as loading control. The corresponding bands to PSMA8 and PSMA7 are indicated in the right of the panel. Note that at the P22 and in adult stages the intensity of both bands abolishes its independent observation. (D) Double immunofluorescence of spermatocytes at pachytene stage obtained from *Psma8*^+/+^ and *Psma8*^−/−^ mice using SYCP3 (red) and a rabbit polyclonal antibody against PSMA8 (green). Green labelling in Psma8^−/−^ spermatocytes (49% of the wild type) represents cross-reactivity of the antiserum with PSMA7. Plot under the image panel represents the quantification of intensity from *Psma8*^+/+^ and *Psma8*^−/−^ spermatocytes. Welch’s t-test analysis: * p<0.01; ** p<0.001; *** p<0.0001. Bar in panels, 10 μm.

Mice lacking PSMA8 did not display any obvious abnormalities but male mice were sterile. Indeed, genetic ablation of *Psma8* leads to a reduction of the testis weight (63.09% lighter; N=6) and to the absence of spermatozoa in the epididymides (Fig 4A-B). Histological analysis of adult *Psma8*^−/−^ testes revealed the presence of apparently normal numbers of spermatogonia, spermatocytes, Sertoli cells and Leydig cells (Fig 4B). Mouse seminiferous tubules can be classified from stage I to XII by determining the groups of associated germ cell types in histological sections. Following these criteria, the spermatogenic process in the mutant testes proceeds normal up to the diplonema spermatocytes in epithelial stage XI. Spermatocytes in meiotic divisions were seen to occur at epithelial stage XII but our counts by squashed analysis of seminiferous tubules revealed the presence of increased numbers of these cells compared to in WT testes (Fig 4C), suggesting a delay in the completion of the meiotic divisions. Together with meiotic divisions, massive numbers of apoptotic cells can be seen that from their size are most likely round spermatids. Indeed, seminiferous tubules in PSMA8 deficient testes contain few if any normal round spermatids and the ones present were often apoptotic, no elongating spermatids were observed (Fig 4B). FACs analysis of whole cells from seminiferous tubules was carried out to verify this conclusion. The results obtained sustained the presence of the haploid compartment in *Psma8*^−/^ testes (Fig 4D). Taken together, we conclude that the PSMA8 deficiency causes a delay during the meiotic divisions with a relative accumulation of spermatocytes at metaphase I/II and that the resulting round spermatids enter apoptosis soon after their formation, leading to nonobstructive azoospermia (NOA) and consequently to infertility. We verified the arrest by lack of immunolabelling with TNP1 and TNP2 (Fig EV3A-B), two transition proteins involved in the replacement of histones by protamines at later stages of spermiogenesis (Barral et al, 2017). We also analysed the presence of H2AL2 (absent from mutant spermatids, see Fig EV3C), a transition histone essential for the first replacement of histones by TNP1 and TNP2 before protamine incorporation (Barral et al, 2017).

**Figure 4.**
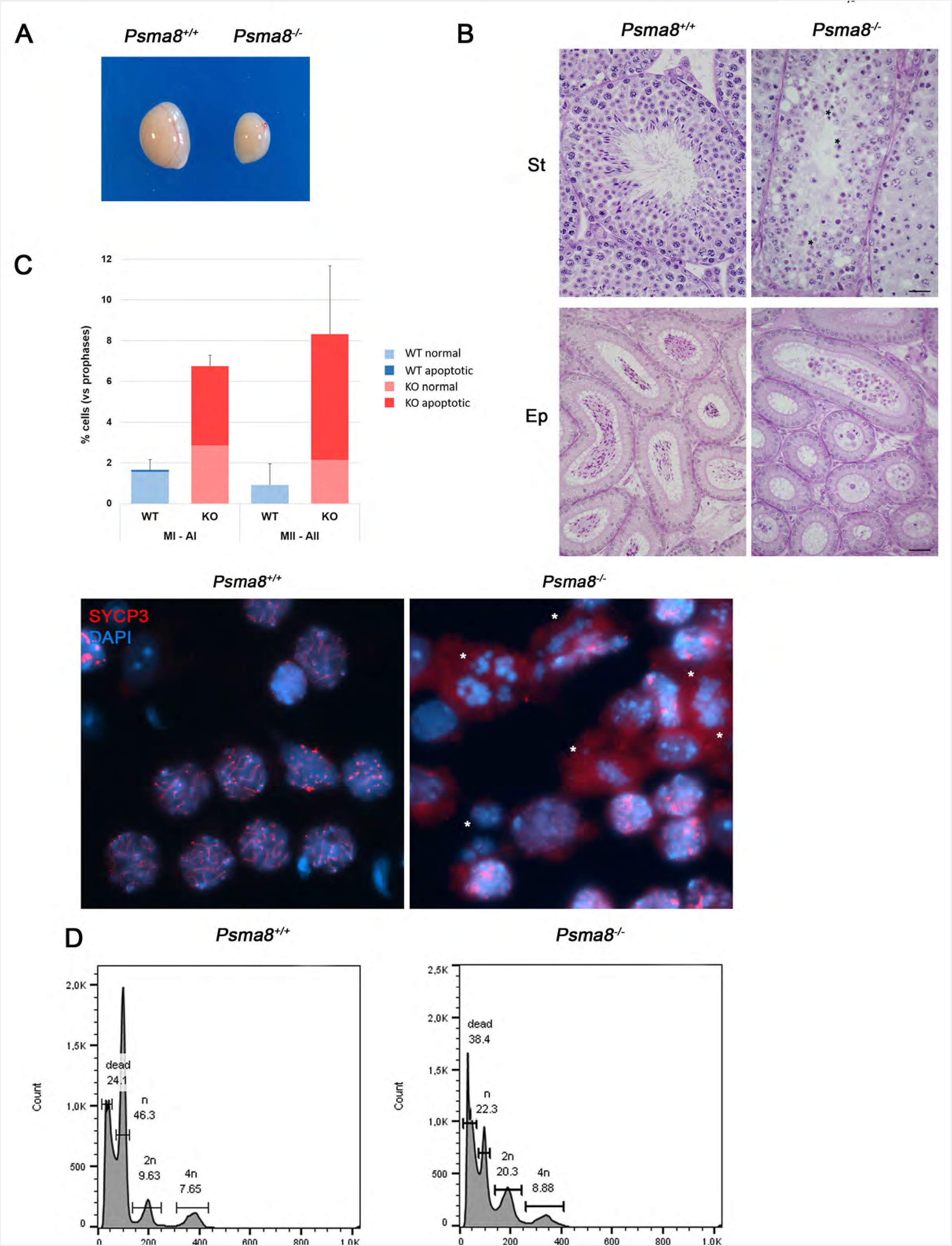
PSMA8 deficiency leads to azoospermia. (A) Genetic ablation of *Psma8* leads to a reduction of the testis size (n=6, WT and KO; Welch’s t-test analysis: p<0.0001), and (B) a complete arrest of spermatogenesis in early round spermatid as shown in PAS and haematoxylin stained testis sections. Massive apoptosis of spermatocytes is indicated (asterisks). The spermatogenic arrest leads to empty epididymides and azoospermia. Bar in upper panels, 100 μm and in lower panels, 5 μm. (**St**) Seminiferous tubules. (**Ep**) Epididymides. (C) Low magnification view of a representative squash preparation of seminiferous tubules showing the accumulation of metaphases I and metaphase II in knockout *Psma8* in comparison with a representative wild-type view (WT, left). The identity of metaphases I (asterisks)/metaphases II was confirmed by the immunolabelling of SYCP3. Chromosomes were counterstained with DAPI (blue). Plot onto the image panel represent the quantification of metaphase I and II (normal and apoptotic) from *Psma8*^+/+^ and *Psma8*^−/−^ tubules. Welch’s t-test analysis: * p<0.01; ** p<0.001; *** p<0.0001. (D) FACs analysis of cells from whole seminiferous tubules from wild type and *Psma8^−/^* showing in both genotypes (N=2) the presence of the tetraploid, diploid and haploid compartment as a result of the early spermatid arrest.

The core subunit PSMA8 co-immunoprecipitates PA200 (Table EV1), an activator of the spermatoproteasome that mediates the acetylation-dependent (ubiquitin independent) degradation of histones in response to DNA damage (Qian et al, 2013). Given the stoichiometric relationship between the CP and RP, we analysed the expression of PA200 by western blot. We observed a complete loss of expression in *Psma8*--deficient mice (undetectable signal, Fig 5A). Similar to PSMA8, PA200 decorates also the LEs of the wild type spermatocytes (Fig 5B). Accordingly, we did not detect PA200 in the LEs of mutant spermatocytes. These results indicate that PSMA8 is necessary or promotes the assembly of PA200 to the CP. Thus the *Psma8* mutation is functionally a double mutant of both PSMA8 and pa200 subunits thus constituting a spermatoproteasome-null mouse mutant.

**Figure 5.**
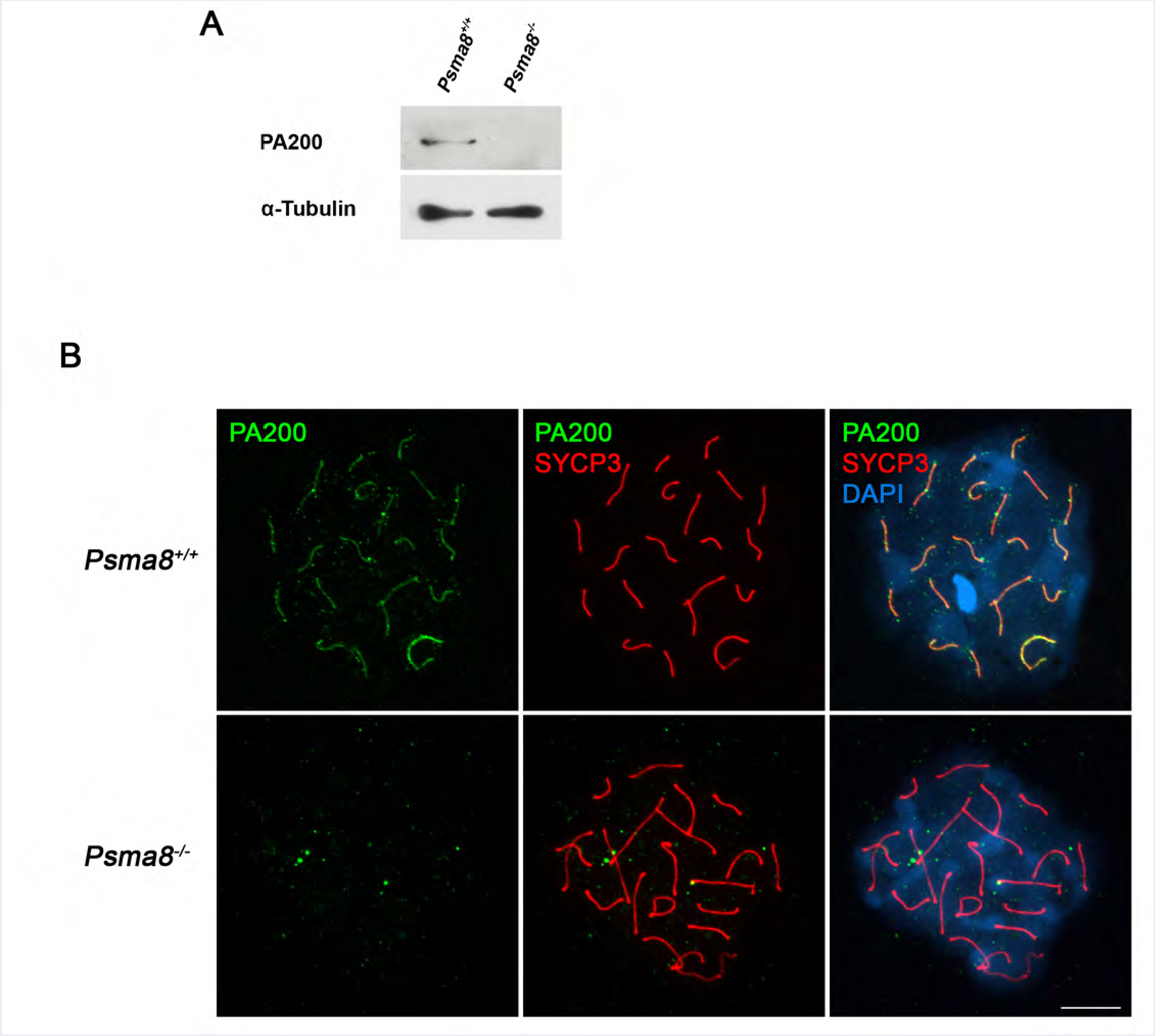
PSMA8 is needed for the expression/assembly of the activator subunit pa200. (A) Western blot analysis of protein extracts from WT and KO testis. A band of the expected molecular weight of pa200 was obtained in the WT but was completely absent in the KO. a-tubulin was used as loading control. (B) Double immunolabelling of pa200 and SYCP3 in chromosome spreads. pa200 is detected at the chromosome axes of the autosomal and XY bivalents during pachytene in wild type spermatocytes in contrast to the absence of labelling in *Psma8^−/^* spermatocytes. Nuclear DNA was counterstained with DAPI. Bar in panels, 10 pm.

### *Psma8* deficient spermatocytes show normal DSB repair, synapsis/desynapsis but have abnormal metaphases I and II

Metaphase I accumulation can occur either because of a difficulty to enter anaphase or because of abnormal events during meiotic prophase I in *Psma8*^−/^ spermatocytes leads to a checkpoint mediated delay. Given that pa200-deficient mice show defects in somatic DNA repair and acetylation-dependent degradation of histones, we tested meiosis progression, DSB processing, and synapsis in *Psma8*^−^ spermatocytes.

To evaluate the role of PSMA8 in meiotic progression and DNA repair, we initially analysed the assembly/disassembly of the SC by monitoring the distribution of the TF protein SYCP1 as co-labelling of SYCP3 and SYCP1 highlights regions of synapsis in wild type testes. We did not observe differences in the synapsis and chromosome behaviour from zygotene to pachytene and during the desynapsis process from diplotene to diakinesis (Fig EV4). The fraction of spermatocytes in metaphase I was increased (1.56±0.6% WT vs 6.7±0.5 KO; increase 77% Table EV3) as well as the number of cells in stage XII and both were apoptotic by Caspase or TUNEL staining (Fig 6A-B). However, the accumulation of metaphase I was not an arrest since we observed cells at the interkinesis stage (by DAPI staining) and also at metaphase II (0.92±1.0 WT vs 8.32±3.3 KO; increase 89%), though a fraction of them were also apoptotic (Table EV3) and showed aberrant labelling of SYCP3 in their centromeres (no SYCP3 labelling is observed in metaphase II chromosomes in wild type cells, Fig 6C). Finally, aberrant round spermatids were observed showing a very fragmented heterochromatin after DAPI staining (Fig 6D). We could no detect any further stage of spermiogenesis in the chromosome spreads (lack of elongating spermatids by immunolabelling for transition proteins, Fig EV3A-C) nor in the histological sections (Fig 4B) indicating that the PSMA8 deficiency leads to an arrest at early round spermatid.

**Figure 6.**
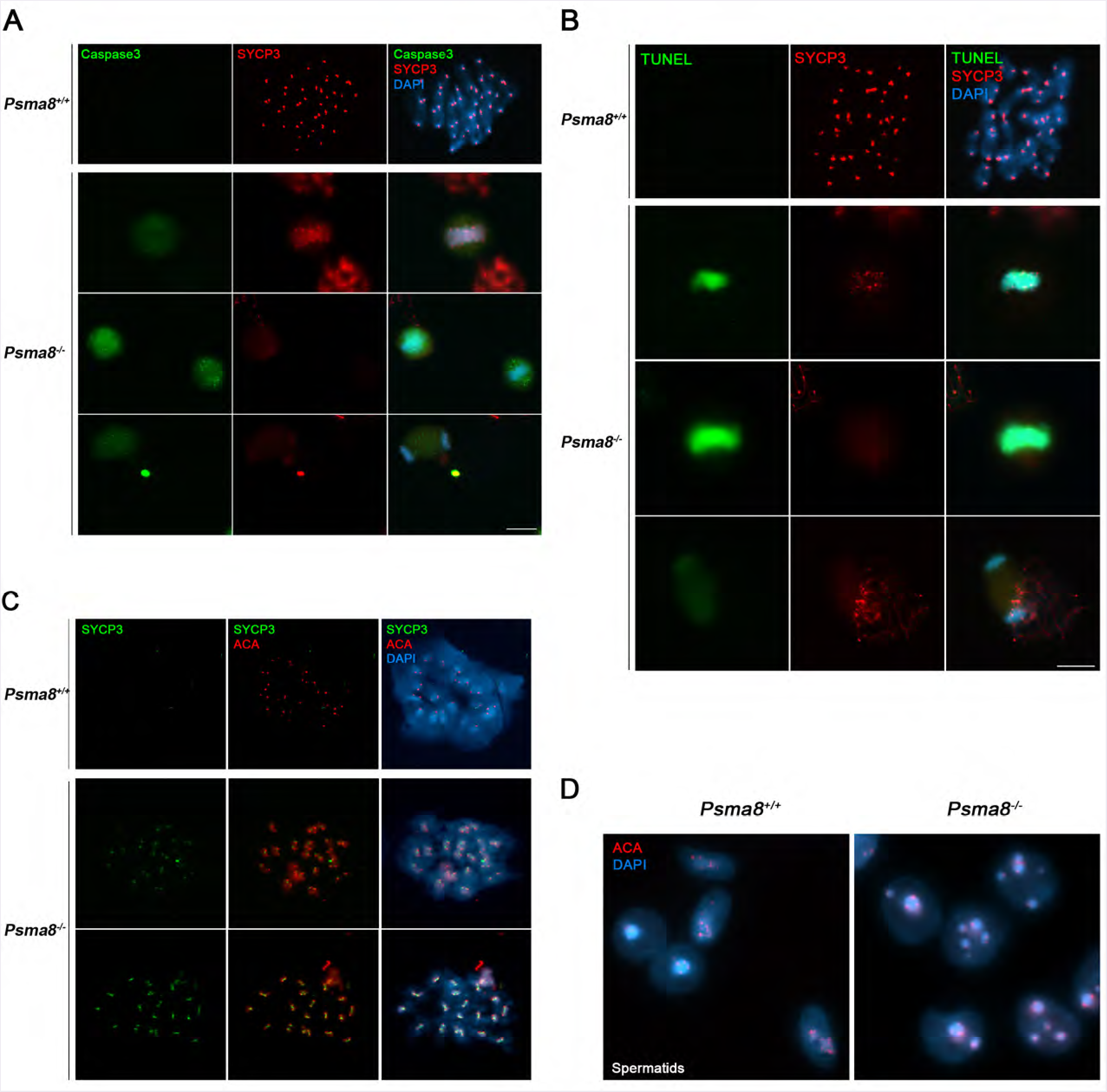
Aberrant metaphase I /II and spermatids in Psma8-deficient mice. (A) Double immunolabelling of Caspase3 (green) and SYCP3 (red) showing normal metaphase I cells and apoptotic metaphase I in chromosomes spreads from seminiferous tubules from *Psma8*^+/+^ and *Psma8*^−/−^ animals, respectively. Chromatin was counterstained with DAPI. Apoptotic cells were pseudocolored in green. (B) Double labelling of TUNEL (green) and SYCP3 (red) was directly performed onto chromosome spreads from WT and KO. *Psma8*^+/+^ metaphase I showing absence of TUNEL labelling and positive SYCP3 labelling at the centromeres. *Psma8r*^−/−^ metaphase I cells showing TUNEL positive staining and SYCP3 positive and negative labelling (upper and mid panel). *Psma8*^−/−^ anaphase I cell showing TUNEL positive staining and SYCP3 positive and negative labelling. (C) Double immunolabelling of SYCP3 (green) with ACA (red) in wild-type and *Psma8^−/−^* spermatocytes at metaphase II. (D) Immunolabeling of ACA (red) and DAPI staining of squashed tubules from *Psma8*^+/+^ and *Psma8*^−/−^ showing unambiguously the presence of aberrant spermatids (ACA and heterochromatin distribution) with atypical chromatin and nuclear shape. Bar in panels, 10 μm.

We next studied the kinetics of DSB repair during meiosis. Meiotic DSBs are generated by the nuclease SPO11 and are then resected to form ssDNA ends that invade into the homologous chromosome. DSBs are marked by the presence of phosphorylated H2AX (γ-H2AX) (Rogakou et al, 1998), which is catalyzed by ATM during a first wave (leptotene) and ATR during the following waves (Jiang et al, 2018). The γ-H2AX distribution in mutant spermatocytes resembles that of wild-type cells in early prophase I (leptonema, WT 82.2±19.6 and KO 81.6±26.7) (Table EV4). As synapsis proceeds to pachytene (similarly in the spermatocytes from *Psma8*^−/−^; see Figs 7A and EV5A), γ-H2AX staining diminished in the autosomes of both wild type and mutant spermatocytes (WT 1.3±0.8 and KO 1.2±0.6, respectively, Figs 7A and EV5A) and appeared mainly at the sex body. From diplotene to diakinesis, β-H2AX labelling is further decreased in the spermatocytes of both genotypes (Fig EV5A). We next analysed if DSBs were processed properly during early stages of recombination. To do that we first studied the distribution of RAD51, a recombinase that promotes homologous strand invasion at the early recombination nodules (Mimitou & Symington, 2009). In wild-type zygonema spermatocytes, RAD51 assembles on numerous foci along the AEs/LEs and finally disappears towards pachytene, with the exception of the unsynapsed sex AEs (Figs 7B and EV5B). We did not observe differences in the number of foci between wild type and *Psma8* deficient spermatocytes from leptotene to late pachytene (Table EV4). Because defective DNA repair finally abrogates crossing over formation (Dai et al, 2017), we then analysed the foci distribution of MLH1, a mismatch repair protein (marker of crossover sites) that functions in the resolution of joint molecules at the end of crossover formation (Moens et al, 2007). We observed a similar value in the KO (24.9±2.5 foci) in comparison with the wild-type (23.3±2.4 foci; Fig 7C). These results indicate that the repair of meiotic DSBs and synapsis/desynapsis proceeds normally during prophase I in the absence of PSMA8 and is not responsible for the observed metaphase I accumulation.

**Figure 7.**
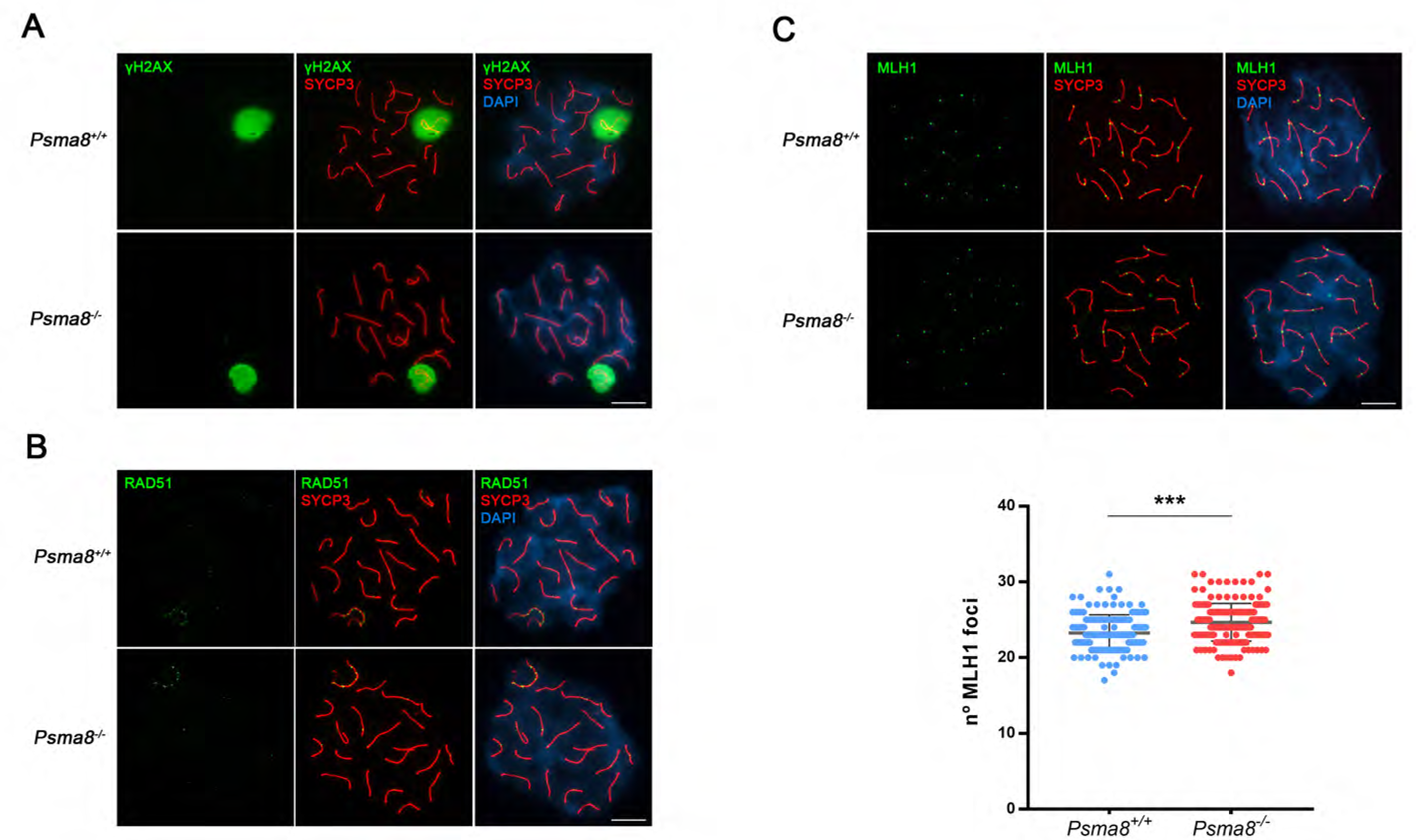
DSB repair is not affected in the absence of PSMA8. (A) Double immunolabelling of γ-H2AX (green) with SYCP3 (red). DSBs are generated at leptotene (see Fig EV5) and at pachytene stage the γ-H2AX labelling disappears from the autosomes and remains only at the sex body of the spermatocytes in both *Psma8*^+/+^ and *PsmaR*^−/−^ (B) Double immunolabelling of SYCP3 (red) and RAD51 (green). RAD51 foci associated to the AEs completely dissociates at pachytene with the exception of the sex body. (C) Double immunolabelling of SYCP3 (red) with MLH1 (green). MLH1 foci are present along each autosomal SC in wild-type and *Psma8*^−/−^ pachynema meiocytes in a similar way. Plot under the image panel represent the quantification of number of foci from *Psma8*^+/+^ and *Psma8*^−/−^ spermatocytes. Welch’s t-test analysis: *p<0.01; ** p<0.001; *** p<0.0001. Bar in panels, 10 μm.

### PSMA8 deficiency abolishes H4ac turnover from late prophase to round spermatid

During spermatogenesis most of the histones are replaced by basic transition proteins, and ultimately by protamines facilitating chromatin compaction. Hyperacetylation of core histones during this process, and specially acetylation of H4, is supposed to play a pivotal role in the initiation of the histone displacement and chromatin ultracondensation (Pivot-Pajot et al, 2003; Shang et al, 2007). In addition, H4ac participates in meiotic DNA repair, chromatin-wide expansion of H2AX phosphorylation and the formation of crossovers during male meiosis (Jiang et al, 2018).

To understand the acetylated-dependent degradation of histones by spermatoproteasome, we measured the acetylation status of three core histones H2acK5, H3ac and H4ac in chromosome spreads by double IF for SYCP3 and the corresponding acetylated histones (Fig 8A-C). This procedure enables a more precise staging of the spermatocytes and a more precise way to quantitate the signals than with the peroxidase immunostaining (IHC) of testis sections (Qian et al, 2013). As expected, the loss of PSMA8 (and pa200) leads to the accumulation of H2acK5, H3ac and H4ac though to a different degree, being the most affected H2acK5, a fact that has been linked to defects in somatic DNA repair.

**Figure 8.**
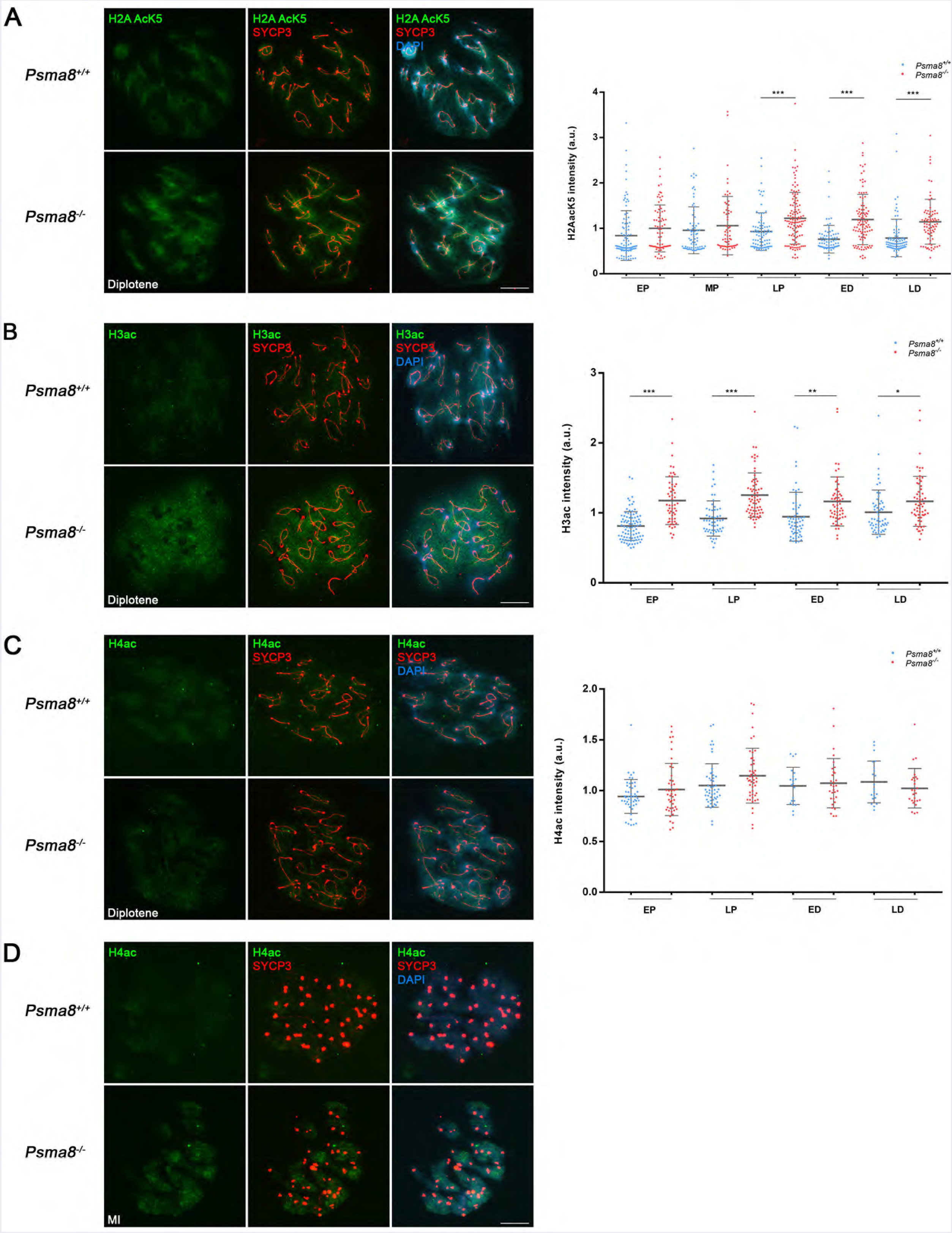
PSMA8 deficiency provokes an slight increase of acetylated histones at prophase I. (A) Double immunolabelling of H2AacK5 (green), (B) H3ac and (C) H4ac with SYCP3 (red) in wild-type and *Psma8*^−/−^ diplonemas. (D) Double immunolabelling of H4ac (green) with SYCP3 (red) in wild-type and *Psma8*^−/−^ metaphase I cells showing a qualitative increase of labelling in the mutant metaphase I. Plots right to each panel represent the quantification of the fluorescence intensity from *Psma8*^+/+^ and *Psma8*^−/−^ spermatocytes at early pachytene (EP), mid pachytene (MP), late Pachytene (LP), early diplotene (ED) and late diplotene (LD) corresponding to the immunolabeling of EV6-8. Welch’s t-test analysis: * p<0.01; ** p<0.001; *** p<0.0001. Bar in panels, 10 μm.

The results show that the levels of H2acK5, H3ac and H4ac are moderated from pachytene to diakinesis with a relative increase of all of them during the later stages of prophase I in the *Psma8*^−/−^ (see Fig. 8A-C and EV6-EV8). We did not detect further staining of H2acK5 and H3ac between late diakinesis spermatocytes and round spermatids in the wild type and arrested spermatids in the mutant cells. In contrast, H4ac also labeled metaphase I chromosomes (whole chromosomes and especially centromeres), interkinesis nuclei and arrested round spermatids, with greater intensity in the mutant cells than in the wild type (Fig 8C).

A role of H4ac in the three waves of H2AX phosphorylation has been recently revealed (Jiang et al, 2018). Despite the slight accumulation of H4ac in mutant prophases, we did not observe defects in this process through the comparison of γ-H2AX staining between WT and *Psma8*-deficient spermatocytes (leptonema and zygonema) including the expansion of γ-H2AX staining to the chromatin of the sex body (in pachynema) by a DMC1-mediated ATR phosphorylation mechanism. This result further support the notion that loss of acetylation-dependent degradation of histones through the spermatoproteasome has no physiological impact onto the meiotic recombination process. In early and round spermatid the only acetylated histone showing labelling is H4ac. Accordingly, H4ac is the only acetylated histone showing an accumulation in the arrested spermatid from spermatoproteasome-deficient mice (Figs 8C and EV8).

Histone displacement, through histone ac-dependent degradation, occurs in round spermatids during the middle of their elongating development (Rathke et al, 2014) and mice lacking pa200 (*Psme4*^−/−^) are fertile (subfertile) and their spermatocytes are not arrested at any stage and generate functional spermatozoa (Khor et al, 2006). The presence of the malformed spermatids in the *Psme4^−/−^* tubules has been explained by the accumulation of acetylated histones (histone displacement) which in turns leads to a delay in the disappearance of the core histones provoking apoptosis (Qian et al, 2013). This is in contrast with the infertility and aberrant transition from metaphase I to early round spermatid stage in the *Psma8* mutant. Thus, this phenotype is more severe and cannot be attributed to accumulation of acetylated histones in spermatids (H4Kac) that could provoke defects in the canonical histone displacement process during mid spermiogenesis.

PSMA8 deficiency is more severe than the pa200 single mutant and that in pa200 and pa28γ double mutant which do not arrest their spermatogenesis despite being infertile *in vivo* but not *in vitro* (spermatozoa are not motile but can fertilize *in vitro* (Huang et al, 2016)). Thus, this genetic analysis shows that the PSMA8-dependent CP has biochemical functions independently of the activators pa200 and pa28γ. Our proteomic analysis together with others (Uechi et al, 2014) further support this notion and indicate that the PSMA8-containing CP can be associated with other regulators such as the 19S subunit, expanding its substrate promiscuity.

### Proteasomal activity in the *Psma8*-deficient mice

We next investigated the biochemical activity of the testis extracts lacking spermatoproteasomes by measuring the chymotrypsin-like activity (corresponding to the catalytic subunit β1), caspase like activity (corresponding to the β5) and the trypsin-like activity (β3) by a standard fluorogenic assay (Gomes et al, 2009) in the presence and absence of SDS (activated proteasome). The results show that the proteasome activity in the *Psma8-* deficient testis extracts is not noticeably down-regulated in comparison with the WT testis extracts. The trypsin-like activity (ARR) was the only proteolytic function with a slight reduction in the KO (see Fig 9A).

**Figure 9.**
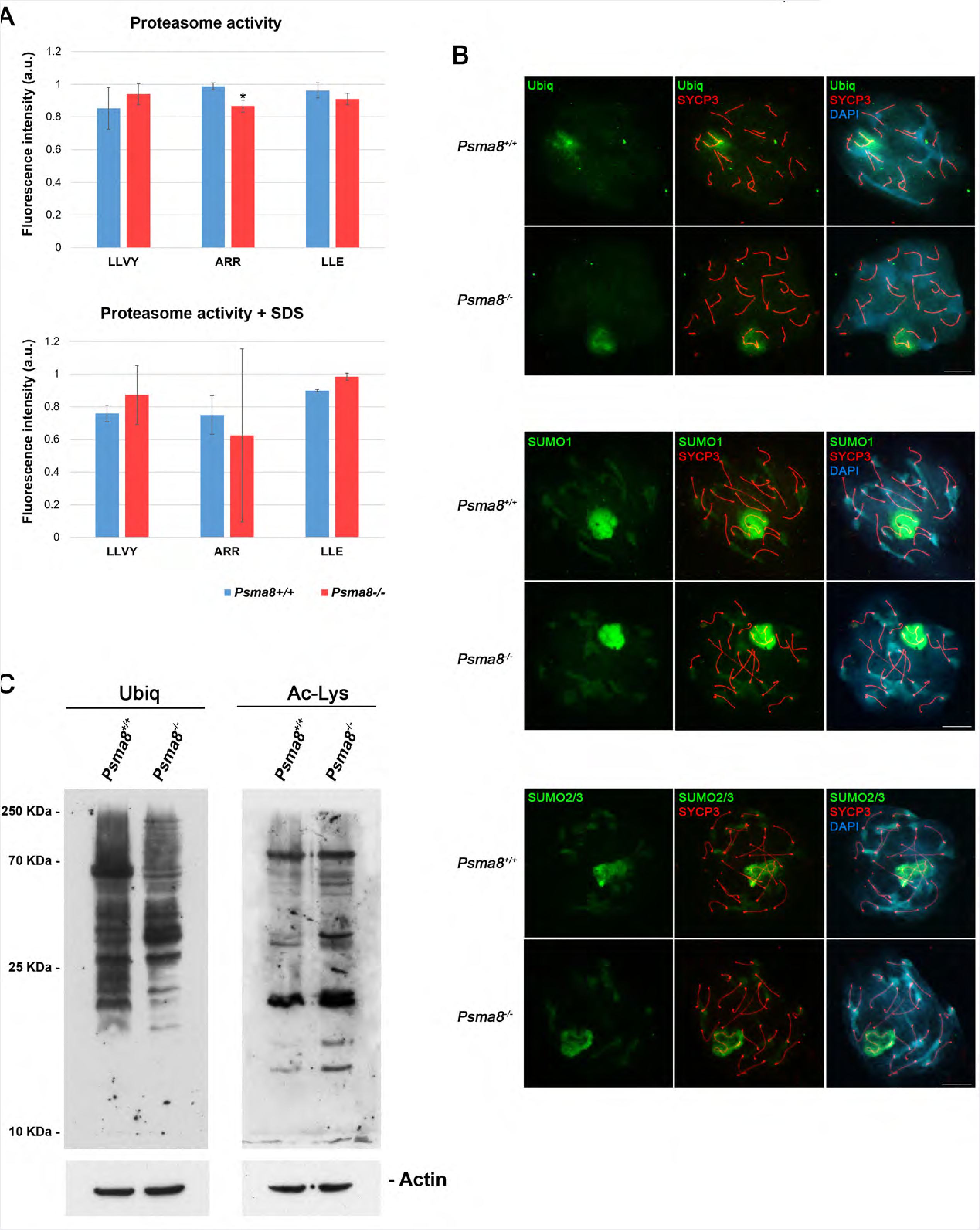
Proteasome activity and global pattern of ubiquitination, sumoylation and acetylation in Psma8-deficient testis. (A) Proteasome activity of *Psma8* deficient testis. 100 μg of protein from whole testis extracts of *Psma8*^+/+^ and *Psma8*^−/−^ mice were inoculated into 96-well plate and the proteasome peptidases activities were measured. The activities relative to WT are shown. Bars, represent Standard Deviation (SD). (B) Double immunolabelling of SYCP3 (red) and Ubiquitin, SUMO1 and SUMO2/3 (green) in mouse spermatocytes in pachytene. Chromatin was stained with DAPI. (C) Western blot analysis of protein extracts from adult wild type and KO testis with a specific antibody against Ubiquitin (Ubiq) and the Ac-Lysine (Ac-Lys). β-actin was used as loading control. Welch’s t-test analysis: * p<0.01; ** p<0.001; *** p<0.0001. Bar in panels, 10 μm.

Overall, these results point show that the spermatoproteasome-deficient testis does not have a drastic reduction of its general proteasome activity, which is likely due to the presence of Pa28α (*Psmel)*, β (*Psme2*) and Pa28γ (*Psme3*) and/or 19S RP complexed to PSMA7-dependent CP given that *Psmel, Psme2* and *Psme3* (and the *Rpt1-14*) genes are actively transcribed in mouse spermatocytes (see dataset1 in (da Cruz et al, 2016)).

To ascertain the degree of activity *in vivo*, we first investigated the steady state levels of protein ubiquitination and acetylation (targeted substrates of spermatoproteasome) in testis during mouse meiosis. By way of IF, we analysed chromosome spreads of spermatocytes using ubiquitin antibodies (Figs 9B and EV9). Contrary to the expected, the results show a slight decrease in the mutant spermatocytes at pachytene and diplotene stages (first stage with relevant labelling, [Fig 9B (upper panel) and EV9]. Similar results were obtained by western blot against ubiquitin from whole extracts of testis (Fig 9C). Because the existing crosstalk between the SUMOylation and the ubiquitylation pathway in the context of protein degradation and its relevance in for crossing over designation and chromosome cohesion (Ding et al, 2018; Rao et al, 2017), we also analysed the pattern of whole sumoylation (SUMO1 and SUMO2/3) during prophase I by IF (Figs 9B and EV10). The results show similar levels of sumoylation in the *Psma8-deficient* spermatocytes (Fig EV10) with the exception of a slight decrease of SUMO2/3 in the late diplotene. These results are in contrast with the observed increase in the sumoylation state of cultured spermatocytes treated with the proteasome inhibitor MG132 (Rao et al, 2017) and suggest a more specific function of the spermatoproteasome in the protein degradation pathway.

Finally, we also roughly evaluated the state of acetylated proteins by western blot, given the acetylated-dependent degradation of histones by PSMA8-containing spermatoproteasomes (Fig 9C). Though the pattern of bands obtained in the western blots are not identical between wild type and mutant extracts (due also to the absence of large spermatids and spermatozoa in the mutant), overall the amount of acetylated signal detected was moderately increased in the *Psma8* mutants (Fig 9C). Altogether, our results indicate that Psma8-deficient mice do not show an accumulation of ubiquitinated (rather a slight decrease is observed) or acetylated proteins in their testis and suggest a specific function of the spermatoproteasome in the controlled degradation of proteins during the spermatogenesis.

### Interactors of PSMA8 in mouse testis

We reasoned that additional substrates might accumulate in the absence of a functional spermatoproteasome. We focused our attention on potential proteins immunoprecipitated with anti-PSMA8 in testis extracts that in principle could explain the meiotic phenotype. Among the list of interactors, we focused on the intermediate filament SYCP1. Because SYCP1 mutant mice are infertile, but otherwise healthy (de Vries et al, 2005), we analysed in more detail the interaction of SYCP1 with PSMA8 and its localization during meiosis (Fig 10). We co-transfected *Sycp1* with *Psma8* in HEK293T cells and we obtained co-IP between SYCP1 and PSMA8 (Fig 10A).

**Figure 10.**
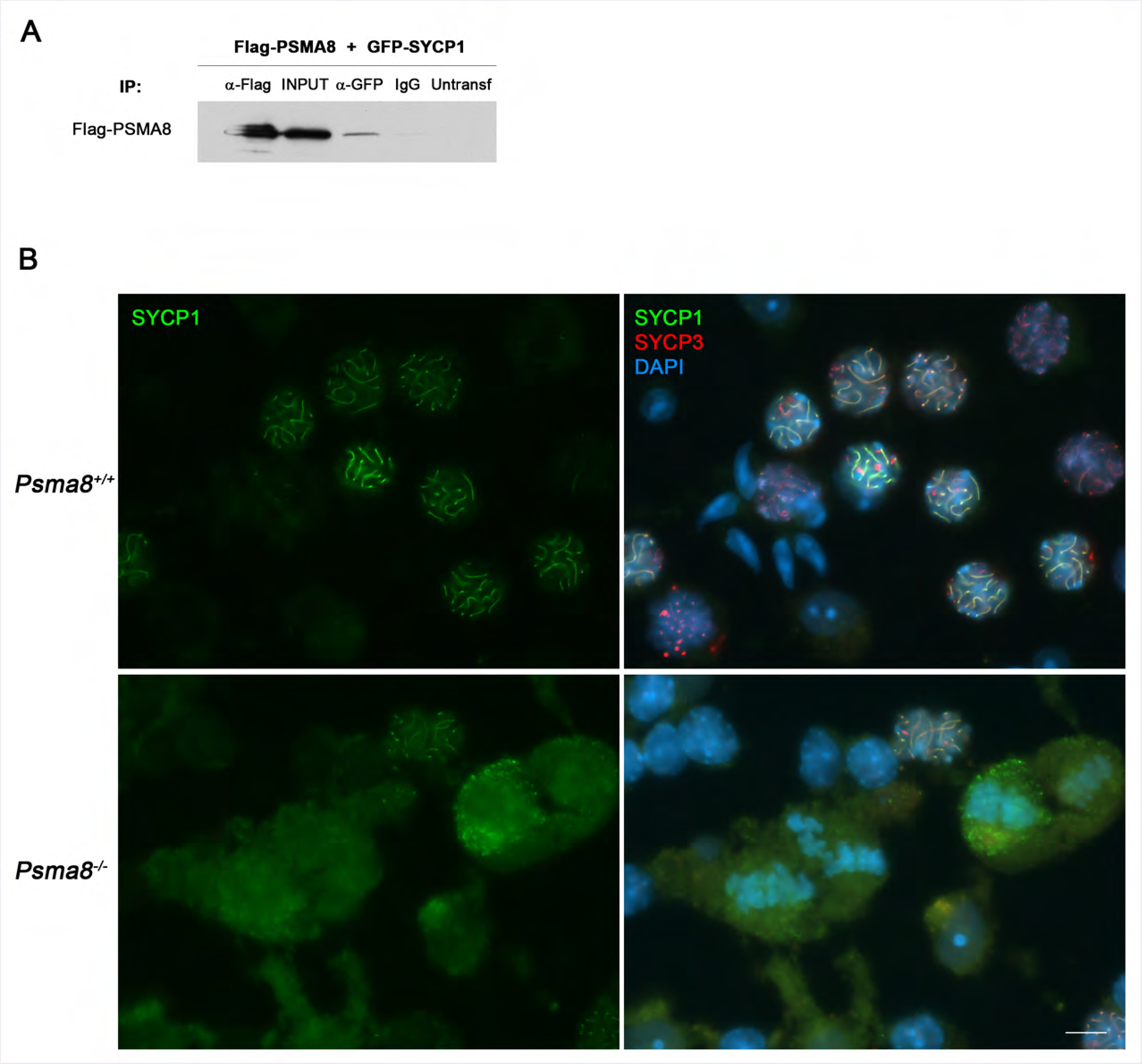
SYCP1 interacts with PSMA8 and is accumulated in metaphase I cells. (A) HEK293T cells were transfected with Flag-PSMA8 and GFP-SYCP1. Protein complexes were immunoprecipitated overnight with either an anti-Flag or anti-EGFP or IgGs (negative control), and were analysed by immunoblotting with the indicated antibody. PSMA8 co-immunoprecipitates (co-IP) with SYCP1. (B) Double immunolabelling of endogenous SYCP1 (green) and SYCP3 (red) in mouse spermatocytes at metaphase I. Chromatin was stained with DAPI (blue). Bar in panels, 10 μm.

Despite the observation that SYCP1 is properly loaded to the synapsed LEs concomitant with chromosome synapsis from zygotene to pachytene and also removed from desynapsed regions (Fig EV3) we only observed an abnormal accumulation of SYCP1 around *Psma8*-deficient metaphase I. This pattern of SYCP1 is never observed in wild type controls (see Fig 10B). However, the widespread labelling surrounding the whole spermatocyte suggests defective degradation with unknown functional consequences in the progression of meiosis.

Because the IP procedure could capture proteins interacting via the UIPD (Sanchez-Lanzas & Castano, 2014), we carried out a validation approach of additional meiotic candidate interactors by co-IP with PSMA8 making use of the same heterologous system of HEK293T cells. These include TEX30, PIWIL1, PIWIL2 and CDK1. All protein-protein interaction assays carried out were negative (Fig EV11) with the exception of the cyclin dependent kinase CDK1 (see Fig 11A). Because of the relevance of CDK1 in metaphase transition and to validate it as a candidate interactor, we first determined by western blot and IF the expression levels of CDK1. The results clearly show that there is more CDK1 (but not the related kinase CDK2 (Fig 11B-C) in the centromeres of metaphase I chromosome from mutant cells (Fig 11B; 19.7±13.4 KO vs 9.6±5.4 WT; increased of 49,2 %). Similar results were obtained by western blot of whole testis extracts (Fig 11D). In order to determine if the increase expression of CDK1 correspond to its active or inactive phosphorylated form, we made use of an antibody against CDK1-tyr15-p (inactivating modification, Fig 11E). The results show no differences in the labelling at the centromeres of the metaphase I chromosomes. Overall these results indicate that there is more active CDK1 in the mutant metaphase I cells. We also analysed the expression levels of the endogenous separase inhibitor securin (PTTG1), a well-known substrate of the proteasome through the E3 ubiquitin ligase Anaphase Promoting Complex/Cyclosome (APC/C). Securin overexpression also delays anaphase I entry in mouse oocytes (Marangos & Carroll, 2008). Our analysis by IF and western blot show similar levels of PTTG1 (Fig 11F), which validates the specific accumulation of CDK1 in mutant spermatocytes. Taken together, we suggest that loss of PSMA8 provokes an increase in CDK1 levels at metaphase I which would lead to the observed delay and accumulation of metaphase I / metaphase II that finally results in apoptotic plates.

**Figure 11.**
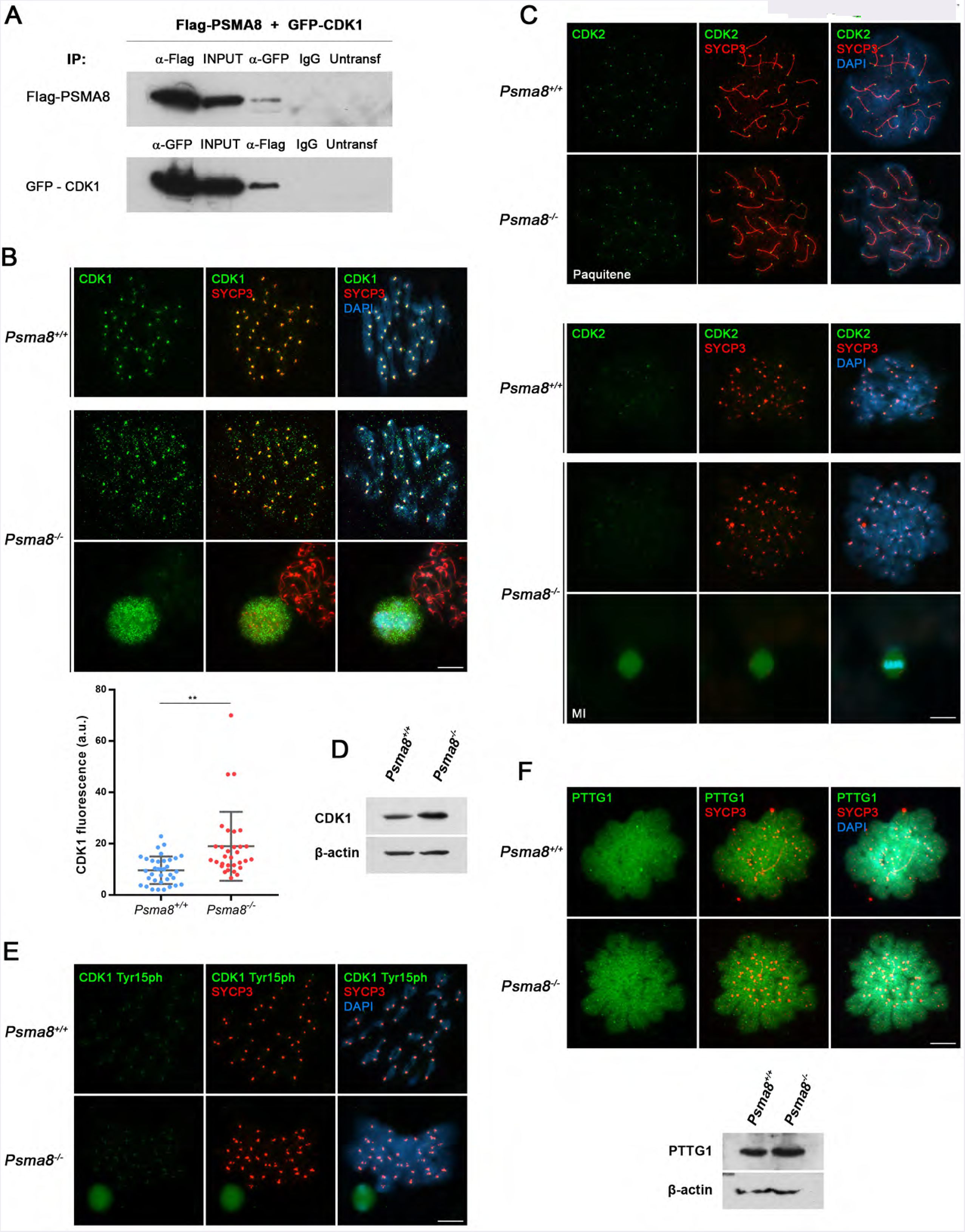
PSMA8 deficiency provokes an accumulation of CDK1 in spermatocytes. **(A)** HEK293T cells were transfected with Flag-PSMA8 and GFP-CDK1. Protein complexes were immunoprecipitated overnight with either an anti-Flag or anti-EGFP or IgGs (negative control), and were analysed by immunoblotting with the indicated antibody. PSMA8 co-immunoprecipitates (co-IP) with CDK1 (as well as reciprocally). (B) Double labelling of endogenous CDK1 (green) and SYCP3 (red) in mouse spermatocytes at metaphase I. Chromatin was stained with DAPI (blue). During metaphase I, CDK1 labels in a slight and disperse way the chromosomes and in a more intensely fashion the centromeres of bivalents. This labelling pattern is enhanced in a normal *Psma8*-deficient metaphase I and greatly enhanced in an aberrant metaphase-like I which shows the accumulation of CDK1 inside the spermatocyte. (C) Double immunolabelling of endogenous CDK2 (green) and SYCP3 (red) in WT and KO mouse spermatocytes in pachytene (upper panel) and metaphase I (lower panel) showing similar labelling at the telomeres and centromeres, respectively (Mikolcevic et al, 2016). (D) Western blot analysis of testis extracts from WT and KO mice with a specific antibody against CDK1. P-actin was used as a loading control. (E) Double labelling of endogenous CDK1-Tyr15phosphorylated (green) and SYCP3 (red) in mouse spermatocytes at metaphase I showing similar expression levels in *Psma8*^+/+^ and *Psma8*^−/−^. Chromatin was stained with DAPI (blue). (F) Double immunofluorescence of PTTG1 (green) and SYCP3 (red) metaphase I and western blot analysis of testis extracts against PTTG1 from *Psma8^−/−^* and *Psma8*^+/+^ showing similar expression levels of PTTG1. Bar in panels, 10 μm.

The role of CDK1 in metaphase-anaphase transition is complex and has a multifaceted function. CDK1 inhibits and activates APC/C by promoting the Spindle assembly checkpoint (SAC) and by a SAC-independent mechanism. The balance between this opposing functions determines cyclin B1 destruction and separase activation giving rise to cohesin cleavage and anaphase onset (Hellmuth et al, 2015). Experimentally, in mouse oocytes *in vivo*, it has been observed that high CDK1 levels can lead to SAC activation (Rattani et al, 2014).

Because CDK1 activates APC/C^Cdc20^ as well as the SAC, it has both positive and negative effects on APC/C^Cdc20^ activity (Rattani et al, 2014). Based on the normal expression levels of PTTG1 in *Psma8^−/−^* metaphase I cells in comparison with the control, it can be argued that there is no precocious APC activation in the Psma8-deficient cells (Fig 11F). Given that in oocytes CDK1 activation of the SAC is dominant over the activation of APC^Cdc20^ (Rattani et al, 2014), we suggest that the prior effect is acting in the Psma8-deficient spermatocytes. Establishing how CDK1 promotes the SAC is still unresolved in oocytes and even more unknown in spermatocytes.

To investigate the structural basis of the PSMA8 localization in the SC and bearing in mind the accumulation of SYCP1 in the spermatocytes lacking PSMA8, we used a candidate gene approach to identify additional putative interactors of PSMA8. We cotransfected *Psma8* with cDNAs encoding each of the known central element proteins (SIX6OS1, SYCE1, SYCE2, SYCE3, and TEX12), transverse filament protein SYCP1, AE protein SYCP3 (Figs 10A, 12A and Fig EV11). As positive controls we used the well-known interaction between SYCE2 and TEX12 (Davies et al, 2012). Surprisingly, we detected co-immunoprecipitation between PSMA8 with SIX6OS1, SYCP1 and SYCE3. Because SYCP3 form filamentous structures in the cytoplasm of transfected cells named polycomplexes (Winkel et al, 2009), co-expression of an interacting protein with SYCP3 may lead to its recruitment to the polycomplexes (Gomez et al, 2016), an indication of protein interaction. We obtained self assembled higher structures when *Psma8* was co-transfected with *Sycp3* (Fig 12B). This SYCP3-dependent cytological interaction was no observed when *Psma7* was co-transfected, further validating the specificity of the interaction given the extensive protein indentity between both PSMA8 and PSMA7. Because of the high complex structures of transfected SYCP3 we were not able to IP SYCP3 (either using a tag or antibodies against the protein) preventing reciprocal co-IP experiments with PSMA8. In order to validate this interaction *in vivo*, we analysed in detail SYCP3 in mouse mutant spermatocytes. We observed SYCP3 aggregates in the nuclei of the *Psma8*-deficient spermatocytes during metaphase I (Fig 12C). This accumulation of SYCP3 is later on manifested in the *Psma8^−/−^* metaphases I/II by an abnormal SYCP3 labelling of the kinetochores of the metaphase II chromosomes (WT metaphase II never show SYCP3 labelling, Fig 12C). Interestingly, it has been previously shown that cultured spermatocytes chemically treated with the proteasome inhibitor MG132 form SYCP3 aggregates (Rao et al, 2017). Altogether, our results suggest that SYCP3 is being targeted for degradation by the spermatoproteasome and that in the absence of PSMA8 its accumulation could be mediating in part the arrest and apoptosis of the spermatocytes.

**Figure 12.**
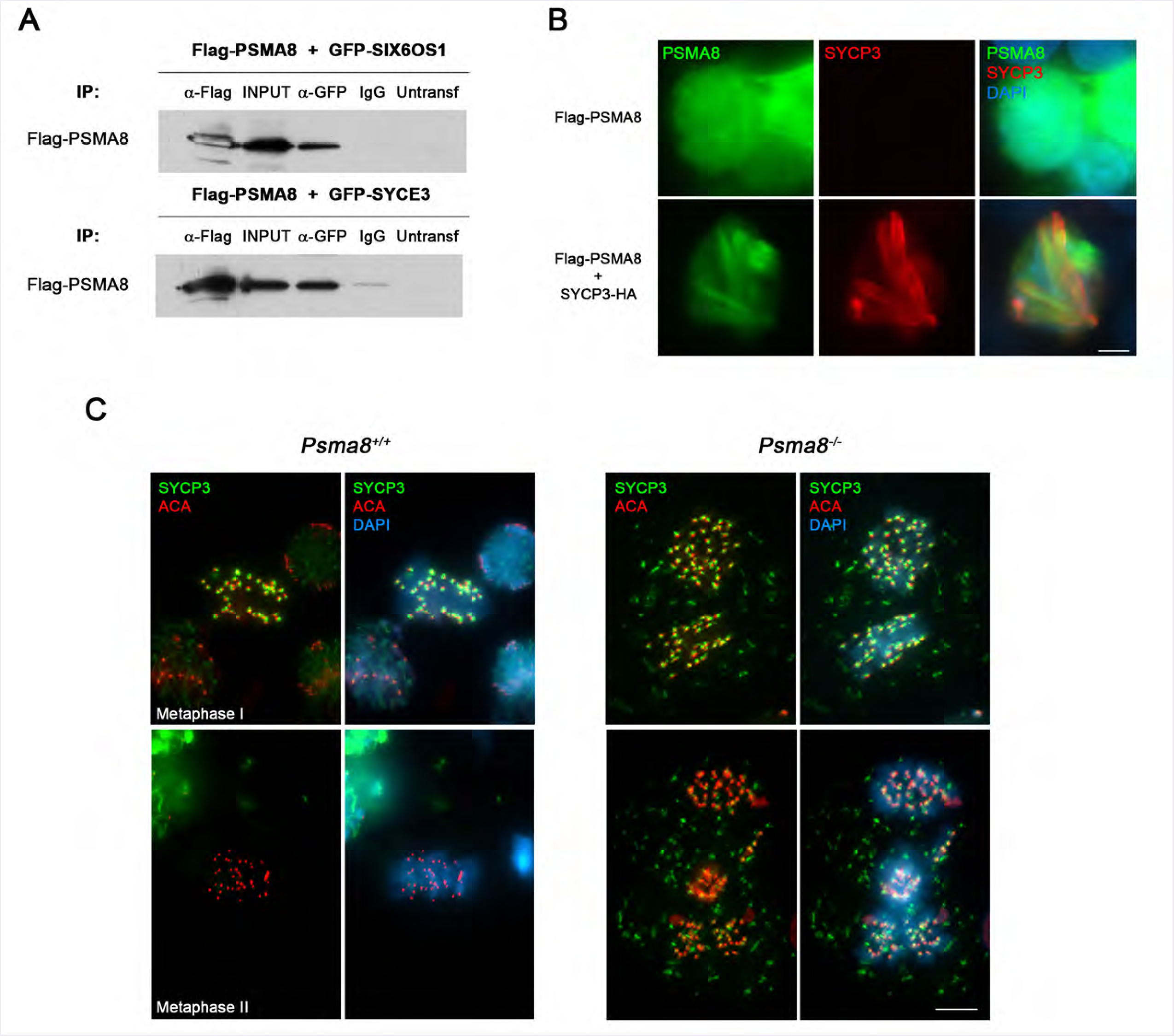
PSMA8 and its interactions with other proteins of the SC. (A) PSMA8 co-IP with SYCE3 and SIX6OS1. HEK293T cells were transfected with plasmids encoding Flag-SIX6OS1, GFP-SYCE3 and Flag-PSMA8. Protein complexes were immunoprecipitated overnight with either an anti-Flag or anti-EGFP or IgGs (negative control), and were analysed by immunoblotting with the indicated antibody. (B) Double immunofluorescence of transfected HEK293T cells with plasmids encoding Flag-PSMA8 alone or together with plasmid encoding SYCP3-HA and immuno-detected with antibodies against PSMA8 (green) or SYCP3 (red). Transfected PSMA8 alone is delocalized and occupies the whole cell whereas when co-transfected with SYCP3-HA is recruited to form polycomplexes. (C) SYCP3 is accumulated *in vivo* in *Psma8*^−/−^ metaphase I and metaphase II. Double immunolabelling of squashed tubules with SYCP3 (green) and ACA (red) in wild-type and *Psma8*^−/−^ spermatocytes at metaphase I and II. *Psma8*^−/−^ metaphases I show labelling for SYCP3 in aggregates (absent in the WT) in addition to its typical labelling at the centromeres. Metaphases II from *Psma8*^−/−^ show labelling for SYCP3 around the kinetochores whereas wild type metaphases II never show any SYCP3 labelling. Bar in panels, 10 μm.

Because overexpressed PSMA8 is able to incorporate into the CP of the somatic proteasome (Uechi et al, 2014), our results should be interpreted in terms of proteasome/spermatoproteasome interactions with the SC proteins and not exclusively to direct interaction with PSMA8. The relationship of the spermatoproteasome with the SC has not been previously analysed in detail in mammals and our results support the idea of a physical anchorage or recruitment to the SC especially through SYCP3, possibly facilitated or mediated by SYCP1, SIX6OS1 and SYCE3 as their most relevant structural partners. Supporting this notion, the Zip1 transverse filament protein of the yeast SC, is the subunit that participates in the recruitment of the proteasome to the SC (Ahuja et al, 2017) which suggests an evolutionary conservation of the mechanism.

### TRIP13 levels are increased in *Psma8*-deficient spermatocytes

Among the “captured” proteins with PSMA8 in the IP, we chose to study TRIP13, a pleiotropic AAA-ATPase (AAA-ATPases associated with diverse cellular activities) (Pch2 in yeast) that participates in meiotic DNA repair and chromosome synapsis through HORMAD interaction and somatic SAC proficiency through MAD2 interaction (Bolcun-Filas et al, 2014; Roig et al, 2010; Wojtasz et al, 2009; Yost et al, 2017). We first conducted IF analysis of TRIP13 in Psma8-deficient and WT spermatocytes (not analysed before). The results (using two independent antibodies) show a sharp labelling of the telomeres starting at zygotene where two dots corresponding to the telomeres are observed faintly. As synapsis progresses, the two dots become more intense and get very closed or even appear as a single dot at pachytene (Figs 13A and EV12). A very faint labelling could also be observed at the axes of the autosomes and XY. From diplotene to dyakinesis the labelling at the telomeres decreases (Figs 13A1 and EV12). However, the intensity of the labelling during prophase I was enhanced in the mutant (5.2±1.5 in pachynema and 4.1±1.4 in late diplonema) in comparison with the wild type (2.9±1.4 in pachynema and 2.3± 1.1 in late diplonema; Fig 13A). Finally, at metaphase I a faint labelling appears at the sister kinetochores of the *Psma8-* deficient spermatocytes which is never observed in the wild type, resembling somatic cells (Wang et al, 2014). These results suggest that TRIP13 is accumulating in the absence of a functional spermatoproteasome.

**Figure 13.**
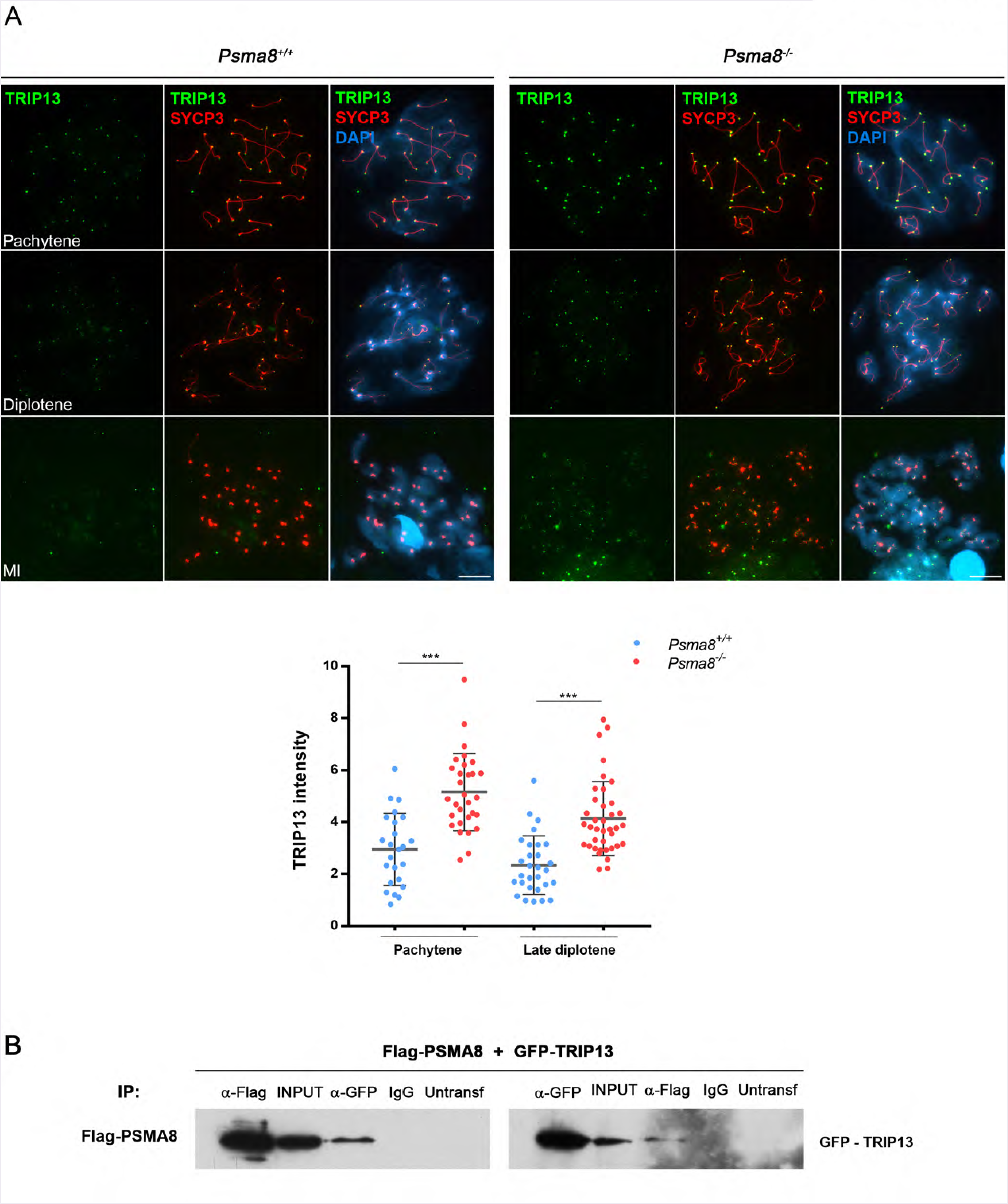
TRIP13 levels are increased in Psma8-deficient spermatocytes. (A) TRIP13 (green) labels the telomeres at pachytene and the intensity of the labelling decreases through desynapsis at diakinesis and diplotene. This labelling is enhanced during prophase I in the *Psma8* mutants but not its main labelling pattern. At metaphase I a faint labelling of sister kinetochores is observed in the *Psma8*^−/−^ spermatocytes that is absent in the wild type. Plot under the panel represents the quantification of the fluorescence intensity from *Psma8*^+/+^ and *Psma8*^−/−^ spermatocytes at pachytene and late diplotene. (B) HEK293T cells were transfected with a plasmid encoding TRIP13-GFP and PSMA8-Flag. Protein complexes were immunoprecipitated overnight with either an anti-Flag or anti-EGFP or IgGs (negative control), and were analysed by immunoblotting with the indicated antibody. Bar in panels, 10 μm.

We next confirmed by co-IP experiments in transfected HEK293T cells that TRIP13-GFP is able to co-immunoprecipitate with Flag-PSMA8 (Fig 13B), further supporting the results obtained in the testis IP experiment and mass spectrometry.

Because HORMADs are one type of the several downstream effectors of TRIP13, we also analysed the distribution and expression of HORMAD1 and HORMAD2, two proteins involved in several processes of meiotic DNA repair and synapsis surveillance mechanisms (Daniel et al, 2011; Wojtasz et al, 2012). We did not observe differences in the expression levels and/or distribution of HORMAD 1/2 in the mutant as compared to the WT spermatocytes (Fig EV13A-B), which is indicative of a normal DNA repair and synapsis.

The functional consequences of an increased expression/stability of TRIP13 has not been determined and cannot be predicted due to the many *in vivo* processes in which TRIP13, through other downstream effectors such as p31 (comet) and MAD2, play a role in meiosis.

Nonetheless, it has been shown in the worm *C. elegans*that in the absence of TRIP13, MAD2 recruitment to the kinetochores was delayed and that in addition to its role in checkpoint silencing, TRIP 13 also contributes to spindle checkpoint activation (Nelson et al, 2015). Thus, it can be postulated that an excess of TRIP13 would increase MAD2 loading to kinetochores delaying mitotic exit as we have observed in the meiosis of the *Psma8*-deficient spermatocytes.

## Discussion

We have shown that the genetic depletion of one structural α-subunit of the spermatoproteasome leads to male mouse infertility. Despite of the proteasome being one of only three tissue-specific proteasomes present in mammalians (together with the immunoproteasome and the thymoproteasome), little is known about its biochemical and physiological function. The groundbreaking work from the Qiu lab showing the acetyl-histone preference of the pa200 subunit of the spermatoproteasome (Qian et al, 2013) yielded novel insight in the field of proteasome-dependent degradation of non-ubiquitylated proteins. However, we have shown that the plasticity of the composition of the spermatoproteasome can explain the mild phenotype of the genetic depletion of pa200 (males are fertile), given the existence of other regulators (activators that can be associated to the CP). Thus, the genetic depletion of the *Psma8* gene generates a more severe male phenotype (it is in fact a double mutant of *Psma8* and pa200) in which it can be ascertained an essential role of the spermatoproteasome during spermatogenesis. Our results show that spermatoproteasome-deficiency provokes severe defects in protein turnover of key meiotic players that affect metaphase I / II exit but not the complex process of meiotic recombination that takes place during the prophase I (crossover). By using a candidate approach of PIPs, we have identified CDK1 and TRIP13 as being the crucial proteins that have an abnormal expression pattern during metaphase I in *Psma8^−/−^* deficient spermatocytes. Given the key roles in all aspects of mitotic/meiotic division that CDK1 and TRIP13 play (including SAC activation), the accumulation of aberrant metaphase I /II spermatocytes in *Psma8^−/−^* deficient mice is to be expected.

Another group of proteins found to be deregulated in the spermatoproteasome-deficient mice are the SC structural proteins SYCP1 and SYCP3. The exact contribution of the accumulation of SYCP1 in the cytoplasm of *Psma8^−/−^* spermatocytes (only detectable in squashed spermatocytes) cannot be experimentally analysed. However, the coiled-coil structure and self-assemblies abilities of SYCP1 strongly suggest a functionally detrimental consequence. Similarly, the presence of SYCP3 in metaphase I spermatocytes and the resulting accumulation at the kinetochores of metaphase II cells, where no SYCP3 is observed in wild type cells, also suggest a functional consequence in the meiotic accumulation and its entrance in apoptosis which is even more pronounced in metaphase II than in metaphase I (Table EV3).

Finally, and as expected from the previous work on pa200 function, Psma8-deficient spermatocytes also accumulated acetylated histones. However, given the arrest of very early round spermatids during Psma8-deficient spermatogenesis where massive acetylation of histones has yet to occur and only a relatively small accumulation of H4ac is observed, it is difficult to argue histone displacement as the molecular mechanism underlying the spermatogenic defect.

The spermatoproteasome through its complex interactome would serve as a hub for the fine tuning coordination of several fundamental key molecules of the spermatogenic process such as analysed during the present work (SYCP1, TRIP 13, CDK1 and acetyl - histones). Our data suggest that deregulation of proteostasis of several meiotic proteins leads to the observed chaotic end of meiosis and the spermatogenic arrest. Thus, we favor an explanation in which the joint contribution of several pathways is responsible for the observed infertility.

From an organismal point of view, *Psma8* transcription is mainly restricted to the testis (Fig 1A). However, as is the case with several other testis genes (named cancer testis genes (Bruggeman et al, 2018)), its expression may be deregulated in some human tumors. Accordingly, *PSMA8* is expressed in Burkitt lymphoma cell lines (i.e. Daudi or Toledo), large B-cell lymphoma and cutaneous melanoma (www.cbioportal.org). The observed *PSMA8* upregulation seems to be mediated by a MYC-dependent transcription binding site present in the proximity of the transcription start site (TSS) of the human *PSMA8* gene (Planello et al, 2014; Rouillard et al, 2016). In addition, this TSS is regulated by DNA methylation by DNMT3B, a methyl transferase involved in lymphomagenesis (Hlady et al, 2012). Thus, we suggest that this MYC binding site is responsible for the overexpression of Psma8 in Myc-dependent tumors such as Burkitt lymphomas and cutaneous melanoma (TCGC). Also in relation with human disease, protein degradation is one of the top cellular functions found in an unbiased differential proteomic profiling of spermatozoa proteins from infertile men with varicocele (Agarwal et al, 2015). In this analysis, PSMA8 is among the top 7 in the list of proteins that are differentially expressed, suggesting a causal role in the severity of the disease.

Taken together, and bearing in mind the PSMA8 dependency of the mouse male germline, we suggest that the spermatoproteasome may be an effective target for male contraception and for the treatment of human malignancies.

## Acknowledgements

We wish to express our sincere thanks to Drs. Liu (Univ of Toledo, USA), A. Toth (Dresden Univ. Germany), J. Schimenti (Cornell Univ, USA), and S. Khochbin (Univ. of Grenoble, France) for providing antibodies (TRIP13, Hormad1, Hormad2, H2AL2) and reagents (plasmid and mice). This work was supported by BFU2017-89408-R. LGH and NFM are supported by European Social Fund/JCyLe grants (EDU/1083/2013 and EDU/310/2015) and YBC by a FPI grant from the MINECO. IR was supported by BFU2016-80370-P. The proteomic analysis was performed in the Proteomics Facility of Centro de Investigation del Cancer, Salamanca, Grant PRB3 (IPT17/0019 - ISCIII-SGEFI / ERDF).

## Material and Methods

### Immunocytology

Testes were detunicated and processed for spreading using a conventional “dry-down” technique or squashing. Antibody against the C-term of PSMA8 was a gift from Dr. Murata (Univ of Tokyo, Japan) and has been previously described (Uechi et al, 2014). Rabbit polyclonal antibodies against PSMA8 were developed by ProteintechTM (R1 and R2) against a fusion protein of poly-His with full length PSMA8 (pET vector) of mouse origin (see Fig EV1 for validation) and was used to validate the immunofluorescence and western results. The primary antibodies used for immunofluorescence were rabbit αSYCP1 IgG ab15090 (1:200) (Abcam), rabbit anti-γH2AX (ser139) IgG #07-164 (1:200) (Millipore), ACA or purified human α-centromere proteins IgG 15-235 (1:5, Antibodies Incorporated), mouse αMLH1 51-1327GR (1:5, BD Biosciences), mouse αSYCP3 IgG sc-74569 (1:100), rabbit αRAD51 PC130 (1:50, Calbiochem), Mouse αCDK1 sc-54 (1:20 IF; 1:1000 wb, Santa Cruz), rabbit αCDK1 Tyr15p #4539 (1:10, Cell Signaling), rabbit αCDK2 sc-6248 (1:20, Santa Cruz), rabbit αPTTG1 serum K783 (1:20 IF, 1:1000 wb), rabbit αTRIP13 19602-1-AP (1:20, Proteintech), rabbit αTNP1 and αTNP2 (1:60, from Dr. Stephen Kistler), rabbit αH2AL2 (1:100, from Dr. Saadi Khochbin), rabbit αPa200 (1:20, Bethyl A303-880A), rabbit α-Caspase3 #9661 (1:30, Cell Signaling), rabbit αH2AacK5 ab45152 (1:20, Abcam), Rabbit αH3ac (K9 and K14) #06-599 (1:20, Millipore), Rabbit αH4ac (K5, K8, K12 and K16) #06-598 (1:20, Millipore), Rabbit αAc-Lys #06-933 (1:1000 wb, Upstate), Mouse αUbiquitin 11023 (1:20 IF, 1:1000 wb, QED Bioscience), Mouse αSUMO1 21C7 (1:40, ThermoFisher), Mouse αSUMO2/3 8A2 (1:30, Medimabs), Rabbit αHORMAD1 and αHORMAD2 (1:50, from Dr. Attila Toth). TUNEL staining of chromosome spreads was performed with the *in situ* cell death detection kit (Roche).

### Cell lines

The HEK293, GC1-spg, leydig TM3, and sertoli TM4 cell lines were directly purchased at the ATCC and cultured in standard cell media.

### Proteasome assay

The 26S proteasome assay was carried out in a total volume of 250 μl in 96 well plates with 2 mM ATP in 26S buffer using 100 μg of protein supernatants from whole extracts of mouse testis. Fluorescently labeled substrates employed were: succinyl-Leu-Leu-Val-Tyr-7-amino-4-methylcoumarin (Suc-LLVY-AMC), Z-Ala-Arg-Arg-AMC (Z-ARR-AMC, Bachem), and Z-Leu-Leu-Glu-AMC (Z-LLE-AMC) for the detection of the chymotrypsin-(β5 catalytic subunit), trypsin-(β2 catalytic subunit) and caspase-(β1 catalytic) like activity measurements respectively. The final concentration of substrate in each assay was 100 pM.

### Histology

For histological analysis of adult testes, mice were perfused and their testes were processed into serial paraffin sections and stained with hematoxylin-eosin or were fixed in Bouin’s fixative and stained with Periodic acid-Schiff (PAS) and hematoxylin.

### In vivo electroporation of testis

Testes were freed from the abdominal cavity and 10 μl of DNA solution (50 μg) mixed with 1pl of 10*FastGreen (Sigma Aldrich F7258) was injected in the rete testis with a DNA embryo microinjection tip. After a period of 1 h following the injection, testes were held between electrodes and four electric pulses were applied (35 V for 50 ms each pulse) using a CUY21 BEX electroporator.

### MS/MS data analysis

Raw MS data were analized using MaxQuant (v. 1.5.7.4) and Perseus (v. 1.5.6.0) programmes (Cox & Mann, 2008). Searches were generated versus the Mus musculus proteome (UP000000589, May 2017 release) and Maxquant contaminants. All FDRs were of 1%. Variable modifications taken into account were oxidation of M, acetylation of the N-term and ubiquitylation remnants di-Gly and LRGG, while fixed modifications included considered only carbamidomethylation of C. The maximum number of modifications allowed per peptide was of 5. For the case of the protein group of CDK1 to 3, experimental results showed that the protein detected was CDK1.

For the PSMA8 antibodies R1 and R2, ratios of their respective iBAQ intensity versus the correspondent iBAQ intensity in the control sample were calculated. Proteins with ratio higher or equal to 5 and two or more unique peptides for at least one RP antibody were selected for ulterior analysis. Aditionally, in order to avoid filtering rare proteins, those with at least one unique peptide and one peptide for both Rabbit antibodies (R1 and R2) and none for anti-IgG were also selected for further analysis.

### Functional and pathway analysis

GO and KEGG over-representation tests were performed using the R package *clusterProfiler* (Yu et al, 2012) using standard parameters except for a FDR cutoff of 0.01. KEGG pathways where some key genes (TRIP 13, CDK1, SYCP1, DDX4, SYCP3, SYCE3, SIX6OS1) operate and the role of the co-immunoprecipitated proteins were studied using the R package*pathview* (Brouwer et al, 2013).

### Immunoprecipitation and western blotting

200 μg of antibody R1 and R2 were bound to 100 ul of sepharose beads slurry (GE Healthcare). Testis extracts were prepared in 50mM Tris Hcl (pH8), 500mM NaCl, 1mM EDTA 1% tritonX100. 20 mg of proteins extracts were incubated o/n with the sepharose beads. Protein-bound beads were packed into columns and washed in extracting buffer for three times. Protein were eluted in 100 mM glycine pH3. The whole immunoprecipitation of PSMA8 was performed in a buffer lacking ATP and glycerol to increase the stringency of the interactors and regulators/activators subunits.

HEK293T cells were transiently transfected and whole cell extracts were prepared and cleared with protein G Sepharose beads (GE Healthcare) for 1 h. The antibody was added for 2 h and immunocomplexes were isolated by adsorption to protein G-Sepharose beads o/n. After washing, the proteins were eluted from the beads with 2xSDS gel-loading buffer 100mM Tris-Hcl (pH 7), 4% SDS, 0.2% bromophenol blue, 200mM β-mercaptoethanol and 20% glycerol, and loaded onto reducing polyacrylamide SDS gels. The proteins were detected by western blotting with the indicated antibodies. Immunoprecipitations were performed using mouse αFlag IgG (5μg; F1804, Sigma-Aldrich), mouse αGFP IgG (4 μg; CSB-MA000051M0m, Cusabio), rabbit αMyc Tag IgG (4μg; #06-549, Millipore), mouse αHA.11 IgG MMS-(5μL, aprox. 10μg/1mg prot; 101R, Covance), ChromPure mouse IgG (5μg/1mg prot; 015-000-003), ChomPure rabbit IgG (5μg/1mg prot.; 011-000-003, Jackson ImmunoResearch), ChomPure goat IgG (5μg/1mg prot.; 005-000-003, Jackson ImmunoResearch). Primary antibodies used for western blotting were rabbit αFlag IgG (1:2000; F7425 Sigma-Aldrich), goat αGFP IgG (sc-5385, Santa Cruz) (1:3000), rabbit αHA IgG (H6908, Sigma-Aldrich) (1:1.000), mouse αMyc obtained from hybridoma cell myc-1-9E10.2 ATCC (1:5). Secondary horseradish peroxidase-conjugated α-mouse (715-035-150, Jackson ImmunoResearch), α-rabbit (711-035-152, Jackson ImmunoResearch), or α-goat (705-035-147, Jackson ImmunoResearch) antibodies were used at 1:5000 dilution. Antibodies were detected by using ImmobilonTM Western Chemiluminescent HRP Substrate from Millipore.

### Generation of plasmids

Full-length cDNAs encoding PSMA8, PSMA7, CDK1, SYCP1 and SYCP3, SYCE2, TEX12, TEX30, PIWIL1 and PIWIL2 were RT-PCR amplified from murine testis RNA. Full-length cDNAs were cloned into the EcoRV pcDNA3-2XFlag or SmaI pEGFP-C1 expression vectors under the CMV promoter. In frame cloning was verified by sanger sequencing.

### Cell lines and transfections

HEK293T cell line was transfected with Lipofectamine (Invitrogen) or Jetpei (PolyPlus). Cell lines were tested for mycoplasma contamination (Mycoplasma PCR ELISA, Sigma).

### Production of CRISPR/Cas9-Edited Mice

Psma8-gRNAs G71 5 - GGGCATACT CCACTTGGAAA −3’ G84 5’-ACCGCGGTAAGCTGCTCCCC-3’ targeting exon 1 and intron 1 were predicted at crispr.mit.edu. Psma8-sgRNAs were produced by cloning annealed complementary oligos at the BbsI site of pX330 (#42230, Addgene), generating PCR products containing a T7 promoter sequence that were purified (NZYtech), and then in vitro transcribed with the MEGAshortscript™ T7 Transcription Kit (Life Technologies). The plasmid pST1374-NLS-flag-linker-Cas9 (#44758; Addgene) was used for generating Cas9 mRNA. After linearization with AgeI, it was transcribed and capped with the mMESSAGE mMACHINE T7 Transcription Kit (AM1345; Life Technologies). RNAs were purified using the RNeasy^®^ Mini Kit (Qiagen). RNAs (100 ng/μl Cas9 and 50ng/μl each guide RNA) were microinjected into B6/CBA F2 zygotes (hybrids between strains C57BL/6J and CBA/J)(Singh et al, 2015) at the Transgenic Facility of the University of Salamanca. Edited founders were identified by PCR amplification (Taq polymerase, NZYtech) with primers flanking exons 1 and intron 1 (Primer F 5-CTTCTCGGTATGACAGGGCAATC-3’ and R 5’-ACTCTACCTCCACTGCCAACCTG-3’) and either direct sequenced or subcloned into pBlueScript (Stratagene) followed by Sanger sequencing. The wild-type and mutant allele were 222 bp and 166 bp long, respectively. The selected founder was crossed with wild-type C57BL/6J to eliminate possible unwanted off-targets. *Psma8*^+/−^ heterozygous mice were resequenced and crossed to give rise to *Psma8*^−/−^ homozygous. Genotyping was performed by analysis of the PCR products of genomic DNA with primers F and R. Mouse mutants for *Rec8* and *Six6os1* have been previously developed (Bannister et al, 2004; Gomez et al, 2016). Mice were housed in a temperature-controlled facility (specific pathogen free, spf) using individually ventilated cages, standard diet and a 12h light/dark cycle, according to EU laws at the “Servicio de Experimentation Animal, SEA”. Mouse protocols were approved by the Ethics Committee for Animal Experimentation of the University of Salamanca (USAL). We made every effort to minimize suffering and to improve animal welfare. Blinded experiments were not possible since the phenotype was obvious between wild-type and Psma8 deficient mouse for all of the experimental procedures used. No randomization methods were applied since the animals were not divided in groups/treatments. The minimum size used for each analysis was two animals/genotype. The mice analysed were between 2 and 4 months of age, except in those experiments where is indicated.

## Author contributions

LGH with the help of NFM, YBC and ELC performed the characterization of the mutant mice including the cytological and biochemical analysis. MSM carried out the Cas9 injections and the testes electroporation. RGV carried out the bioinformatics proteomic analysis. IsR carried out infertility phenotyping of mutant mice. IgR provided unpublished information and antibodies of TRIP13. DdR performed the histological analysis of the phenotype of the mutant mice. AMP and ELC designed the experiments and wrote the paper with the input of the remaining authors (DdR).

---

**Figure EV1.**
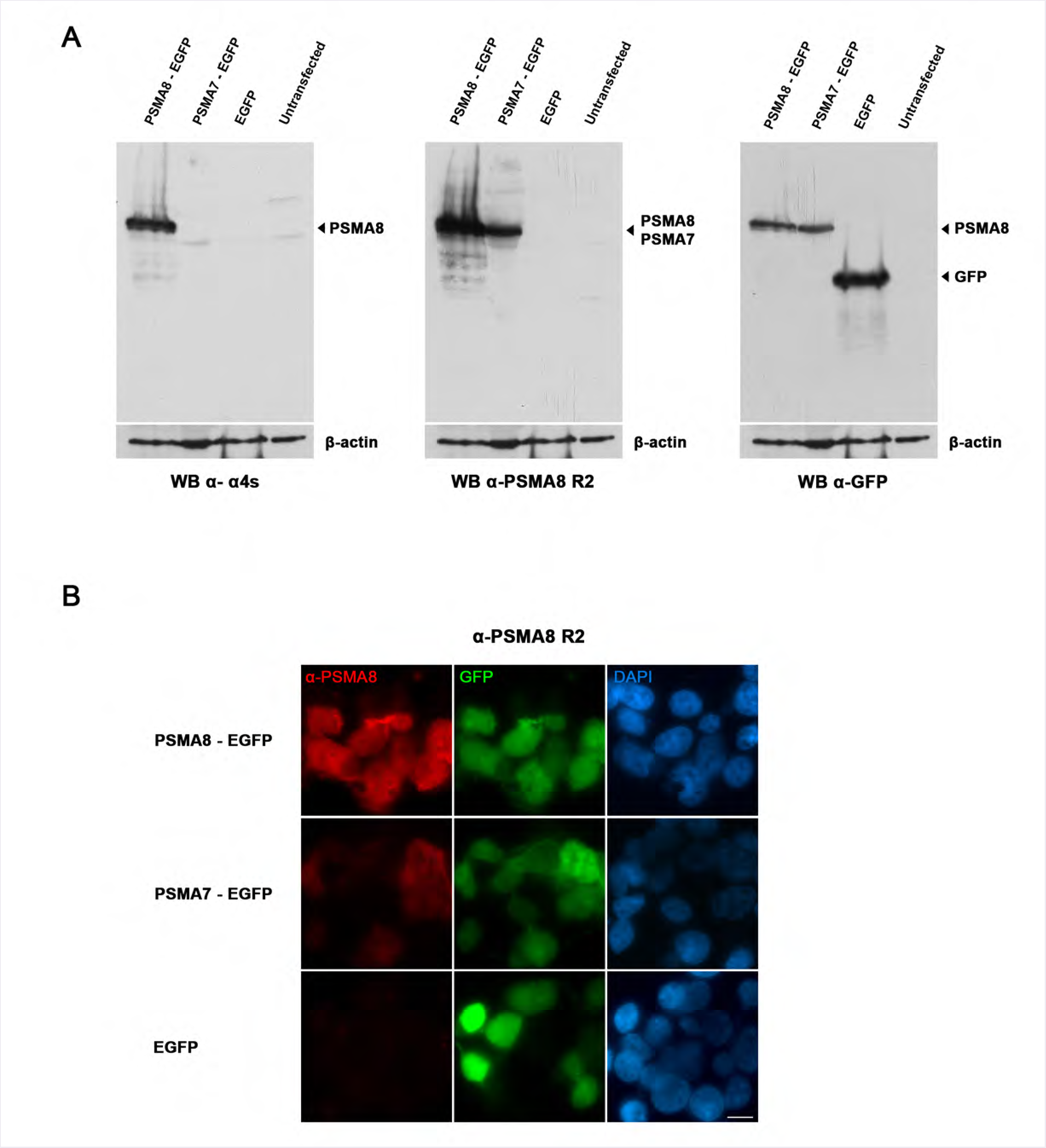
Validation of the antibodies raised against PSMA8. (A) HEK293T cells were transfected with a plasmid encoding PSMA8-GFP, PSMA7-GFP or GFP and the whole extracts were analyzed by western blot using rabbit α-PSMA8-Cterm (left panel, 4S), rabbit α-PSMA8 (central panel, R2) and α-GFP (right panel, GFP). Immunodetection of α-tubulin was used as loading control. The corresponding band of 60 kDa representing PSMA8-GFP was obtained only with the α-α4S. Both bands of 60 kDa representing PSMA8-GFP and PSMA7-GFP were detected with the rabbit a-Psma8 R2. The bands of 60 kDa (PSMA7 and PSMA8) and 30 kDa (GFP) were all detected with the goat α-GFP (arrowheads) validating the experiments. (B) Immunofluorescence of HEK293T cells transfected with plasmids encoding PSMA8-GFP, PSMA7-GFP or GFP. Both PSMA8 and PSMA7 were detected with rabbit α-PSMA8-R2 (red) and GFP by direct fluorescence signal (green). Green and red signals co-localize in the cytoplasm of the transfected HEK293T cells. The experiments were reproduced three times. Bars represent 10 μm.

**Figure EV2.**
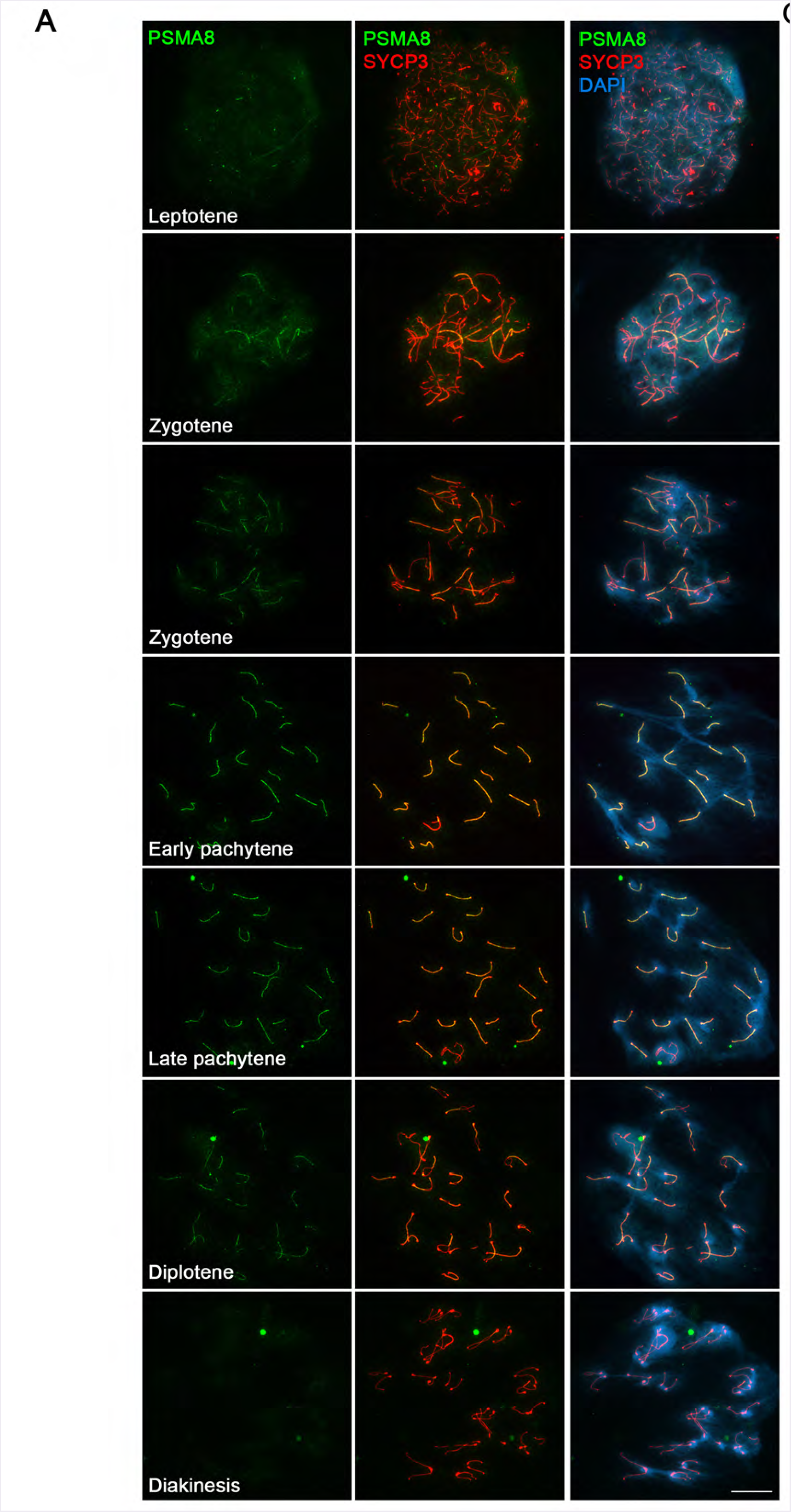
Localization of PSMA8 in mouse spermatocytes. (A) Double labelling of endogenous PSMA8 (green) and SYCP3 (red) in mouse spermatocytes. DNA was stained with DAPI (blue). From the leptotene to zygotene stage, PSMA8 is detected at the synapsed autosomal LEs. At pachytene, PSMA8 is located at the totally synapsed axes and at the PAR of the sex XY bivalent. In diplotene, PSMA8 localizes at the still synapsed LEs and disappears at diakinesis. Bars represent 10μm.

**Figure EV3.**
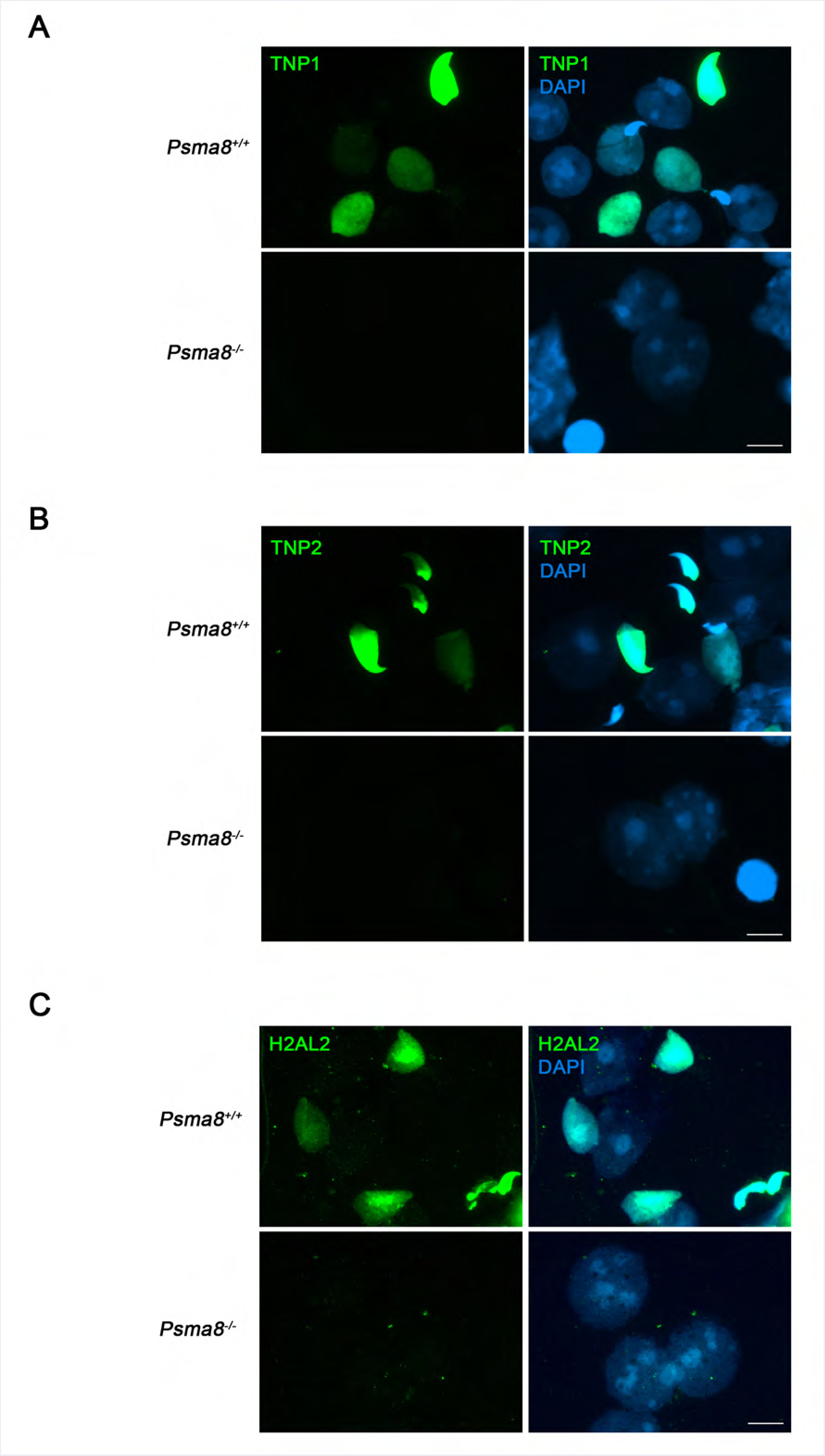
*Psma8*^−/−^ arrested spermatids do not express the transition proteins TNP1, TNP2, H2AL2. Immunolabelling of TNP1 (A), TNP2 (B) and H2AL2 (C) (green) show positive staining in round and elongatid spermatids from wild type mice but lack of staining in *Psma8*^−/−^ mice. Chromatin was stained with DAPI. Bars represent 10 μm.

**Figure EV4.**
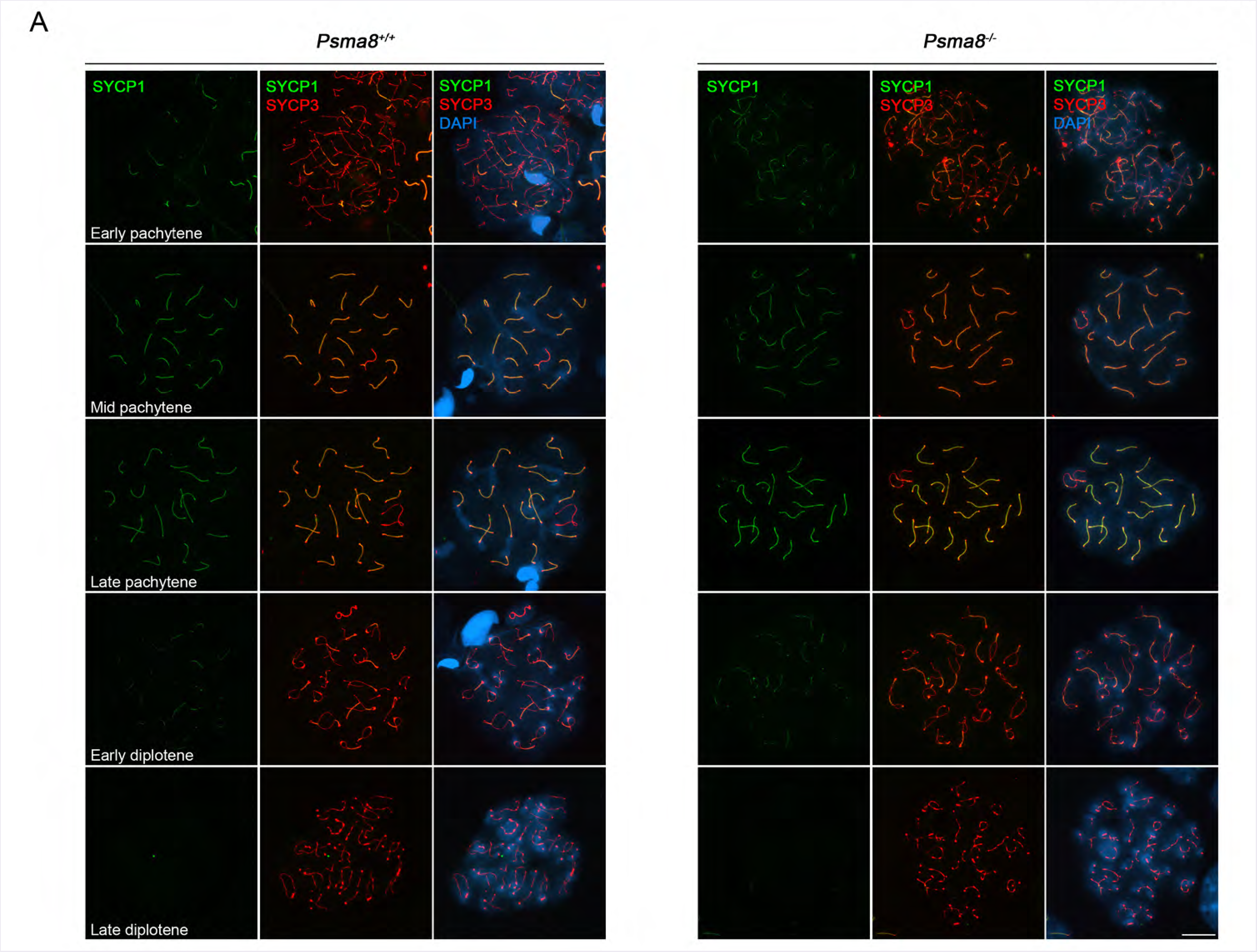
Normal synapsis and desynapsis in spermatocytes lacking PSMA8. Double immunolabeling of SYCP3 (red) and SYCP1 (green) showing normal synapsis and desynapsis from early pachytene to late diplotene. DNA was stained with DAPI. Bars represent 10 μm.

**Figure EV5.**
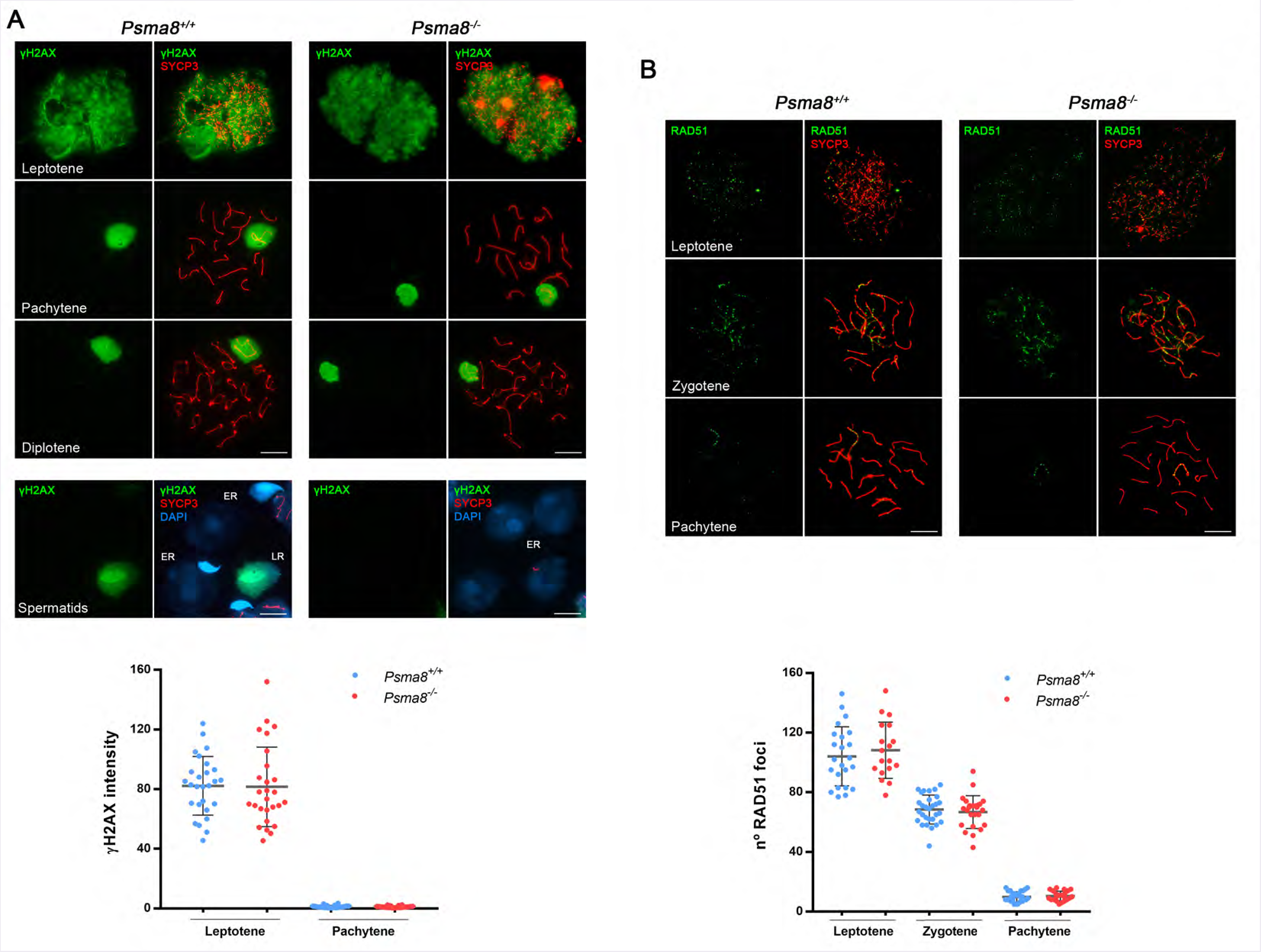
DSBs are generated and repaired in spermatocytes lacking PSMA8. Double immunolabelling of γ-H2AX (green) with SYCP3 (red) in wild-type and *Psma8*^−^ spermatocytes from leptotene to diplotene (upper pannel). In WT and KO leptonemas, γ-H2AX labels intensely the chromatin. After repair at pachytene, γ-H2AX labelling remains only in the chromatin of the unsynapsed sex body. Late round spermatids (LR) but not early round spermatid (ER) from wild type mice show positive staining for γ-H2AX but these highly differentiated cells are lacking in the *Psma8^−/−^* tubules which are arrested at early round spermatids without γ-H2AX staining (bottom panel). (B) Double immunolabelling of SYCP3 (red) and RAD51 (green). RAD51 foci associates to the AEs in leptonema spermatocytes of both genotypes (similar number of foci) and dissociate towards pachytene with a similar kinetics. Bars represent 10 μm. Plots under each image panel represent the quantification of intensity or number of foci from *Psma8*^+/+^ and *Psma8*^−/−^ spermatocytes. Welch’s-test analysis: *p<0.01; ** p<0.001; ***p<0.0001. (Table EV4).

**Figure EV6.**
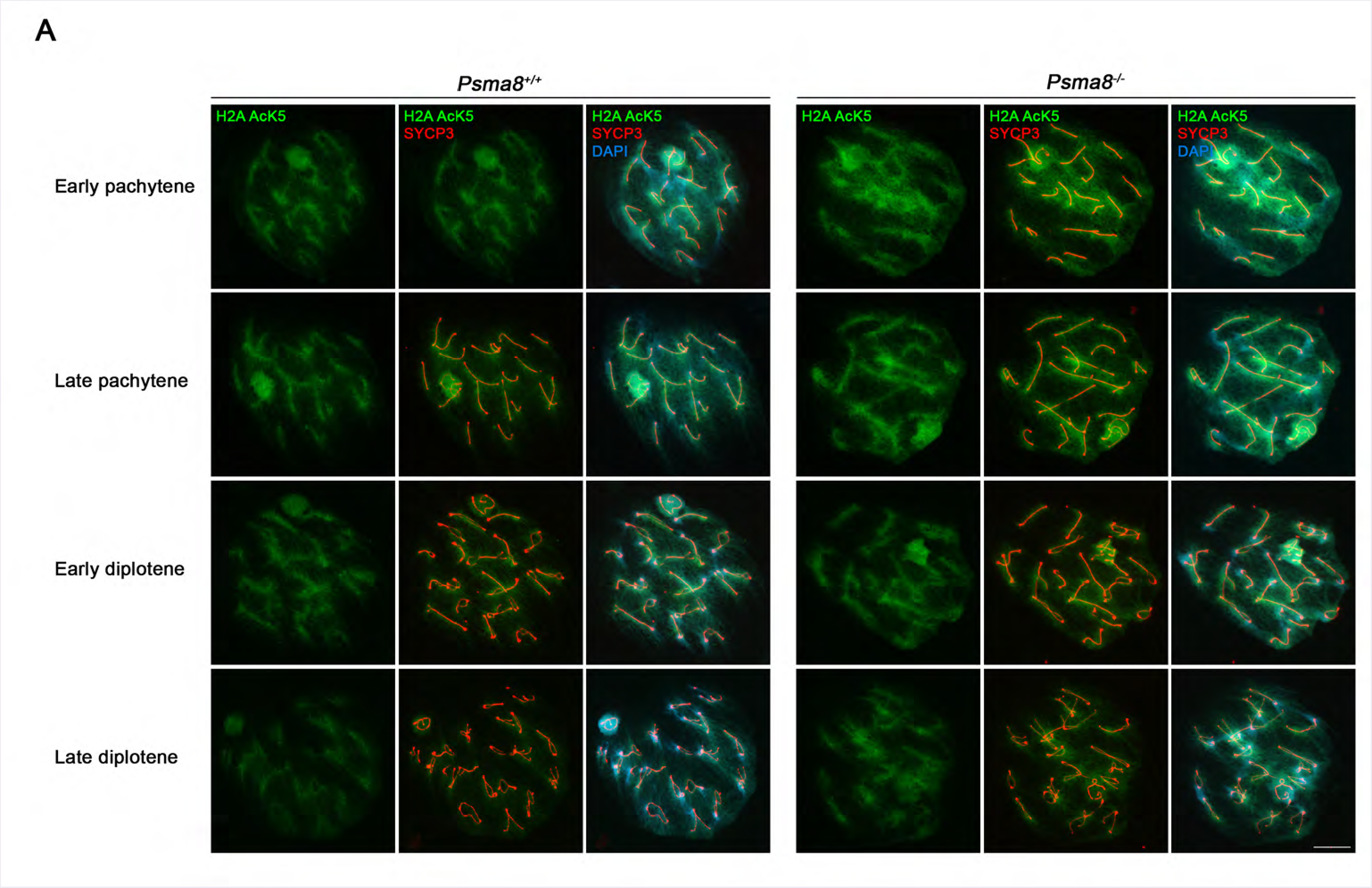
PSMA8 deficiency provokes an slight increase of H2AacK5 at prophase I. (A) Double immunolabelling of H2AacK5 (green) with SYCP3 (red) in wild-type (left panel) and *Psma8*^−/−^ spermatocytes (right panel). In WT and KO spermatocytes chromatin start to be labelled at early pachytene around chromosomes axes. Plots from each panel representing the quantification of fluorescence intensity from *Psma8*^+/+^ and *Psma8*^−/−^ spermatocytes are in Fig 8A. Bars represent 10 μm.

**Figure EV7.**
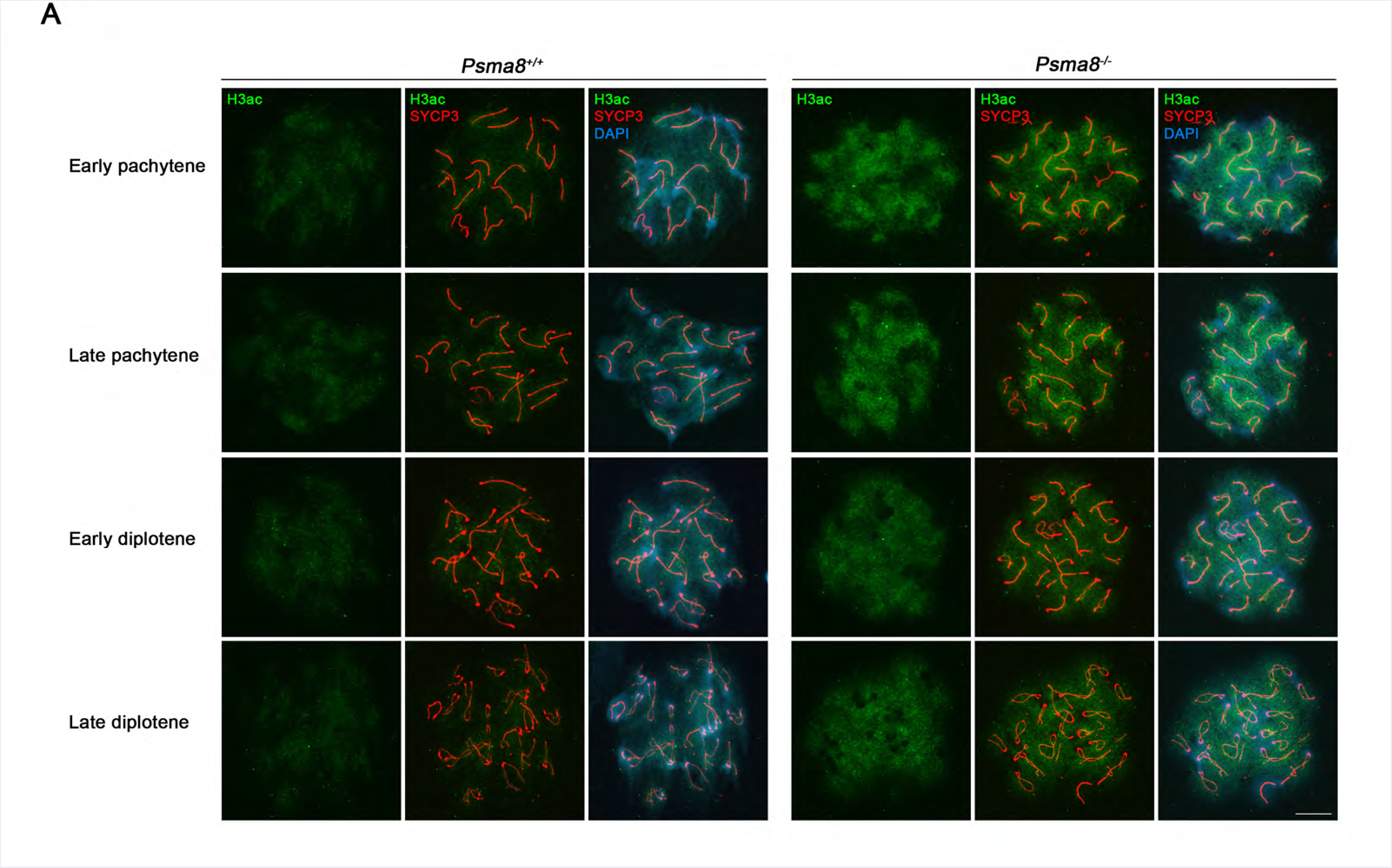
PSMA8 deficiency provokes an slight increase of H3ac at prophase I. (A) Double immunolabelling of H3ac (green) with SYCP3 (red) in wild-type (left panel) and *Psma8*^−/−^ spermatocytes (right panel). Spermatocytes from *Psma8*^+/+^ and *Psma8*^−/−^ show labelling for H3ac at early pachytene in a very diffuse manner surrounding chromosomes axes. Plots from each panel representing the quantification of fluorescence intensity from *Psma8*^+/+^ and *Psma8*^−/−^ spermatocytes are in Fig 8B. Bars represent 10 μm.

**Figure EV8.**
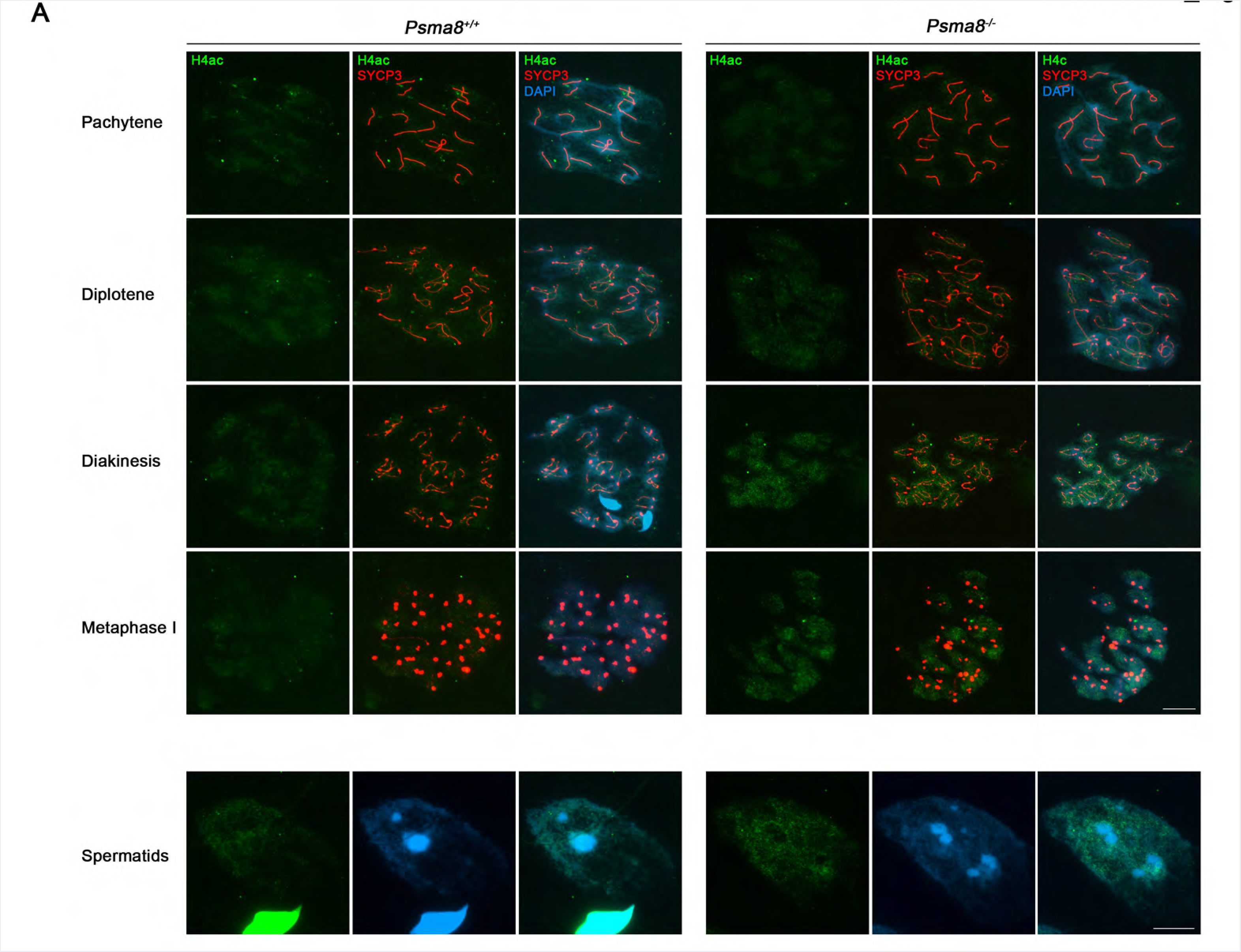
PSMA8 deficiency provokes an slight increase of H4ac at prophase I and in round spermatids. (A) Double immunolabelling of H4ac (green) with SYCP3 (red) in wild-type and *Psma8*^−/−^ spermatocytes. Spermatocytes from *Psma8*^+/+^ and *Psma8*^−/−^ show labelling for H4ac in a very diffuse manner surrounding chromosomes from pachytene to metaphase I (right panel). In wild type metaphase I, H4ac labelling appears weakly painting the chromosomes and on some of the centromeres. However, *Psma8*-deficient cells show a more intense labelling specially at the centromeres. (Lower panel) Round spermatid from *Psma8*^−/−^ accumulates H4ac labelling at the chromatin in comparison with the WT. Plots from each panel representing the quantification of fluorescence intensity from *Psma8*^+/+^ and *Psma8*^−/−^ spermatocytes are in Fig 8C. Bars represent 10 μm.

**Figure EV9.**
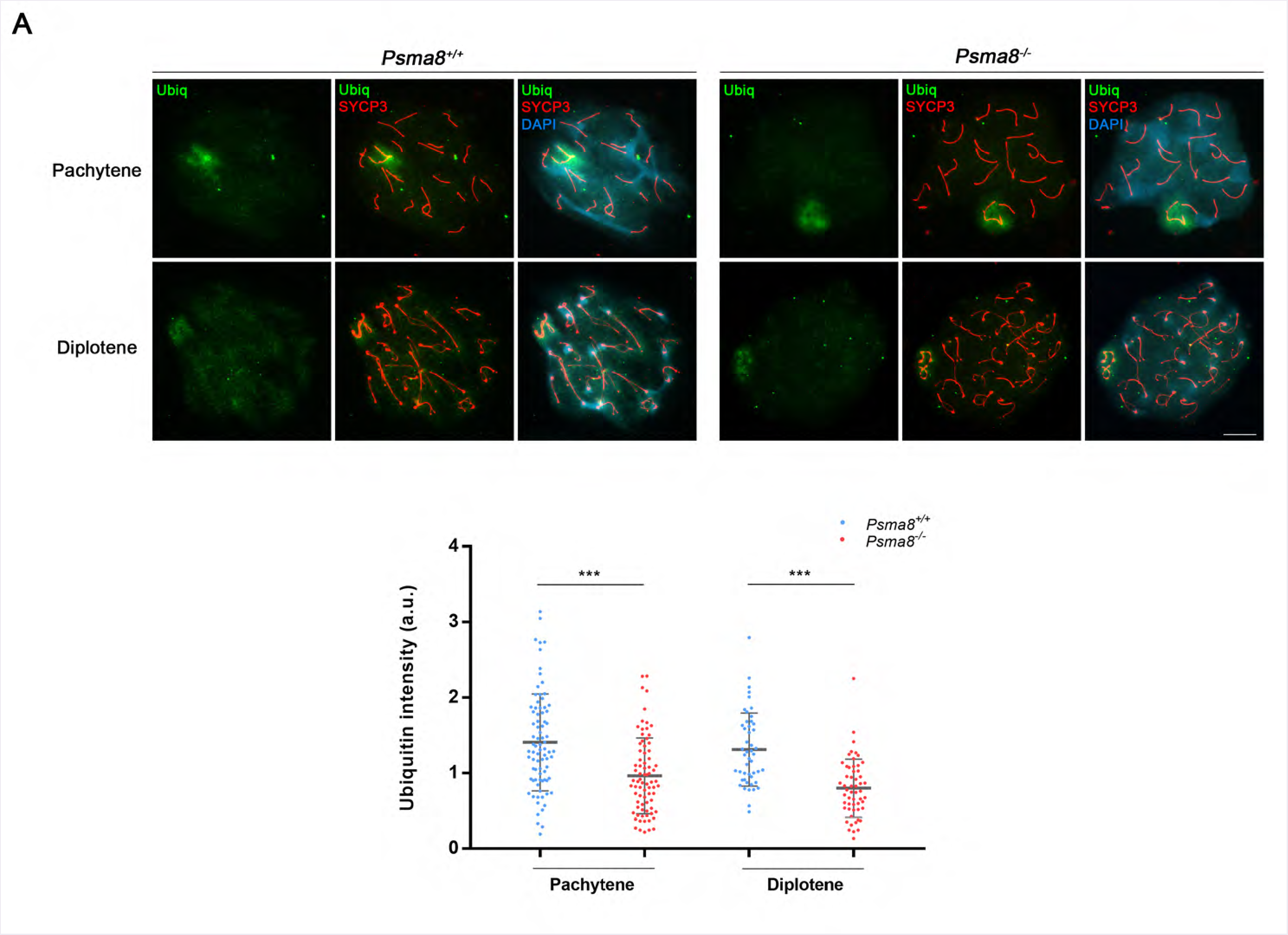
PSMA8 deficiency does not affect global ubiquitination during prophase. (A) Double immunolabelling of Ubiquitin (green) and SYCP3 (red) of spermatocytes at pachytene and diplotene stage from *Psma8*^+/+^ and *Psma8*^−/−^ Ubiqutin labelling decorates faintly the chromatin around chromosome and more intensely the sex body in a similar fashion in both genotypes. Bars represent 10 μm. Plot under the image panel represents the quantification of fluorescence intensity from wild-type and mutant meiocytes. Welch’s *t*-test analysis: *p<0.01; **p<0.001; ***p<0.0001.

**Figure EV10.**
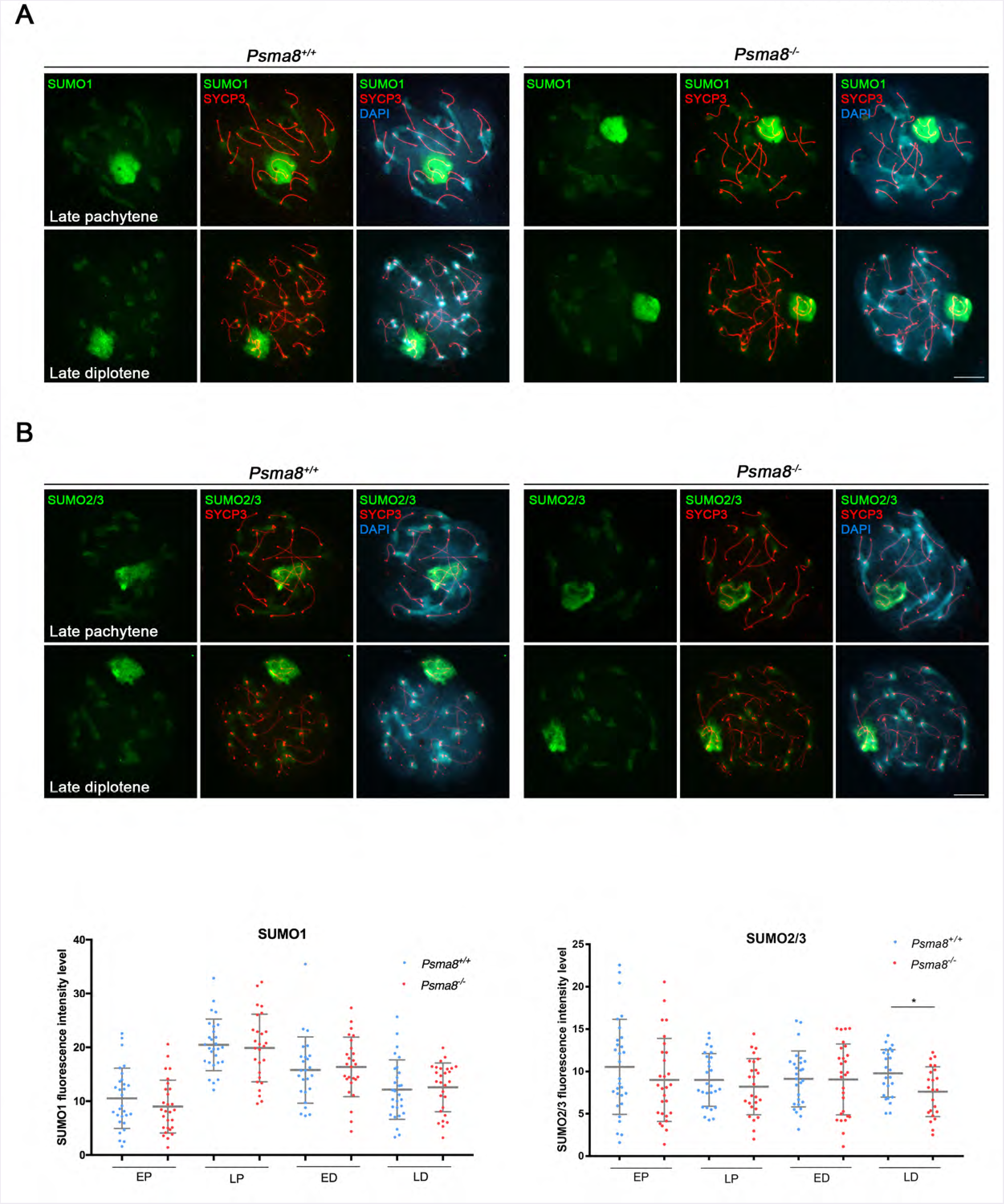
PSMA8 deficiency does not affect global sumoylation during prophase I. (A) Double immunolabelling of SUMO1 (green) and SYCP3 (red) of spermatocytes at late pachytene and late diplotene stage from *Psma8*^+/+^ and *Psma8*^−/−^ SUMO1 labelling decorates the chromosome axes with very low intensity and the sex body intensely. (B) Double immunolabelling of SUMO2/3 (green) and SYCP3 (red) of spermatocytes at late pachytene and late diplotene stage from *Psma8*^+/+^ and *Psma8*^−/−^. SUMO2/3 labelling is very similar to SUMO1. Bars represent 10 μm. Plots under each image panel represent the quantification of intensity from wild-type and mutant meiocytes. Welch’s *t*-test analysis: *p<0.01; **p<0.001; ***p<0.0001.

**Figure EV11.**
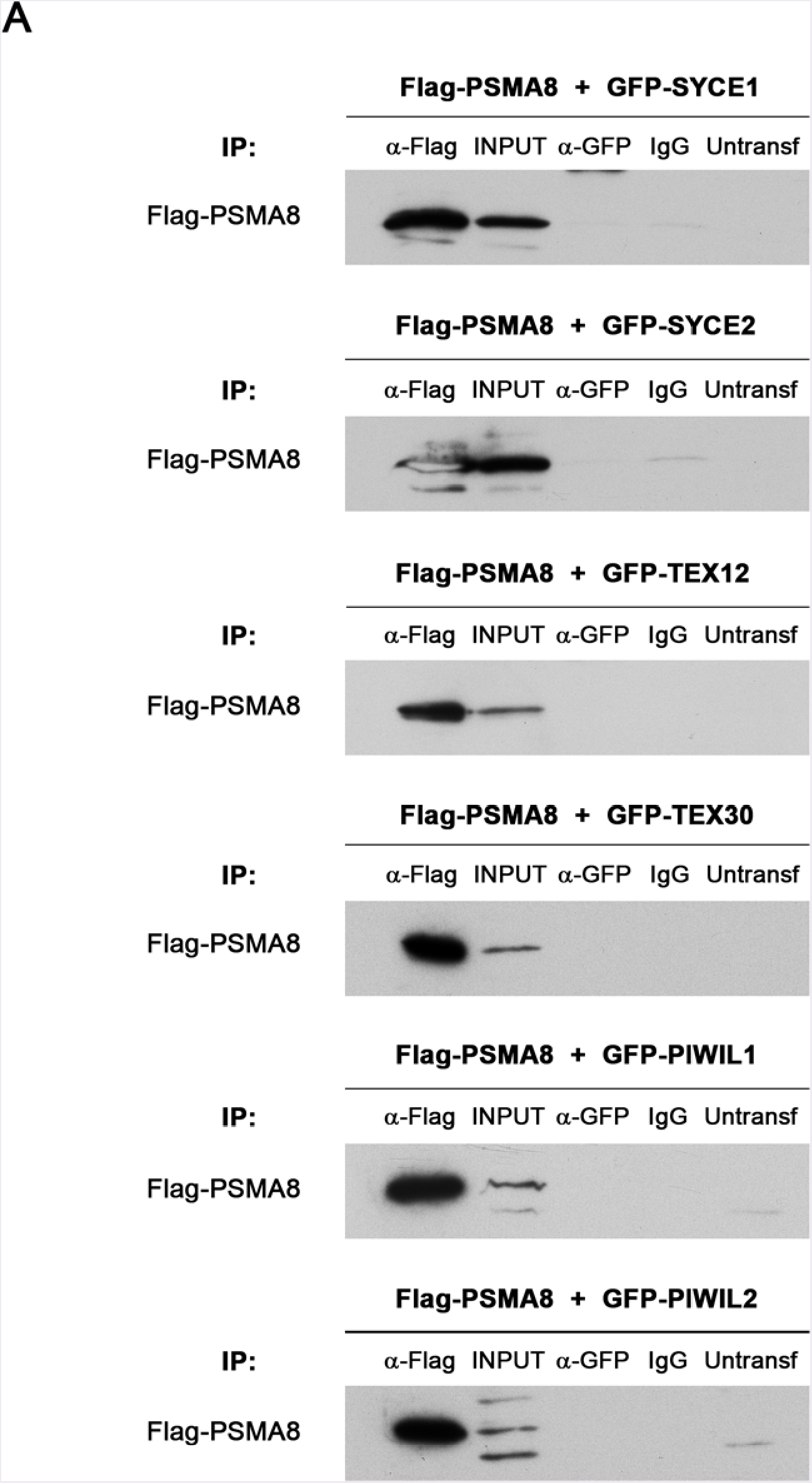
Lack of co-immunoprecipitation of PSMA8 with candidate interactors. (A) HEK293T cells were co-transfected with GFP-SYCE1, GFP-SYCE2, GFP-TEX12, GFP-TEX30, GFP-PIWIL1, and GFP-PIWIL2 and with Flag-PSMA8. Protein complexes were immunoprecipitated overnight with either an anti-Flag or anti-EGFP or IgGs (negative control) and were analysed by immunoblotting with the indicated antibody. PSMA8 does not co-immunoprecipitates (co-IP) with any of them.

**Figure EV12.**
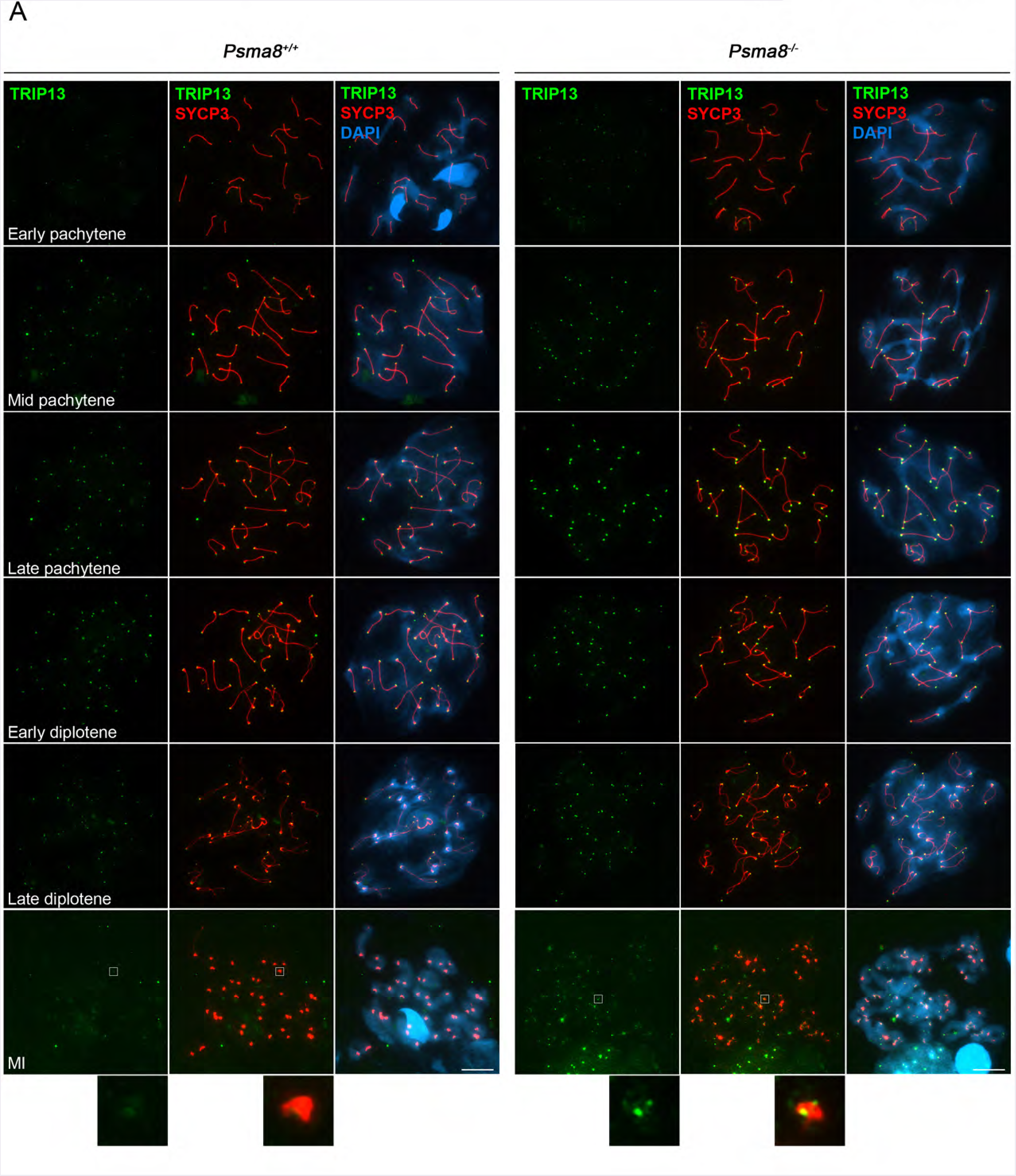
TRIP13 levels are increased in *Psma8*-deficient spermatocytes. (A) The results (using two independent antibodies) show a sharp labeling of the telomeres starting with low intensity at zygotene, increasing at pachytene and decreasing again through desynapsis at diplotene and diakinesis. The intensity of the TRIP13 (green) labelling is enhanced during prophase I in the *Psma8* mutants but not its labelling pattern. At metaphase I a faint labelling of sister kinetochores is observed in the *Psma8*-deficient spermatocytes that is absent in the wild type (faint stain over the centromeres). Bars represent 10 μm.

**Figure EV13.**
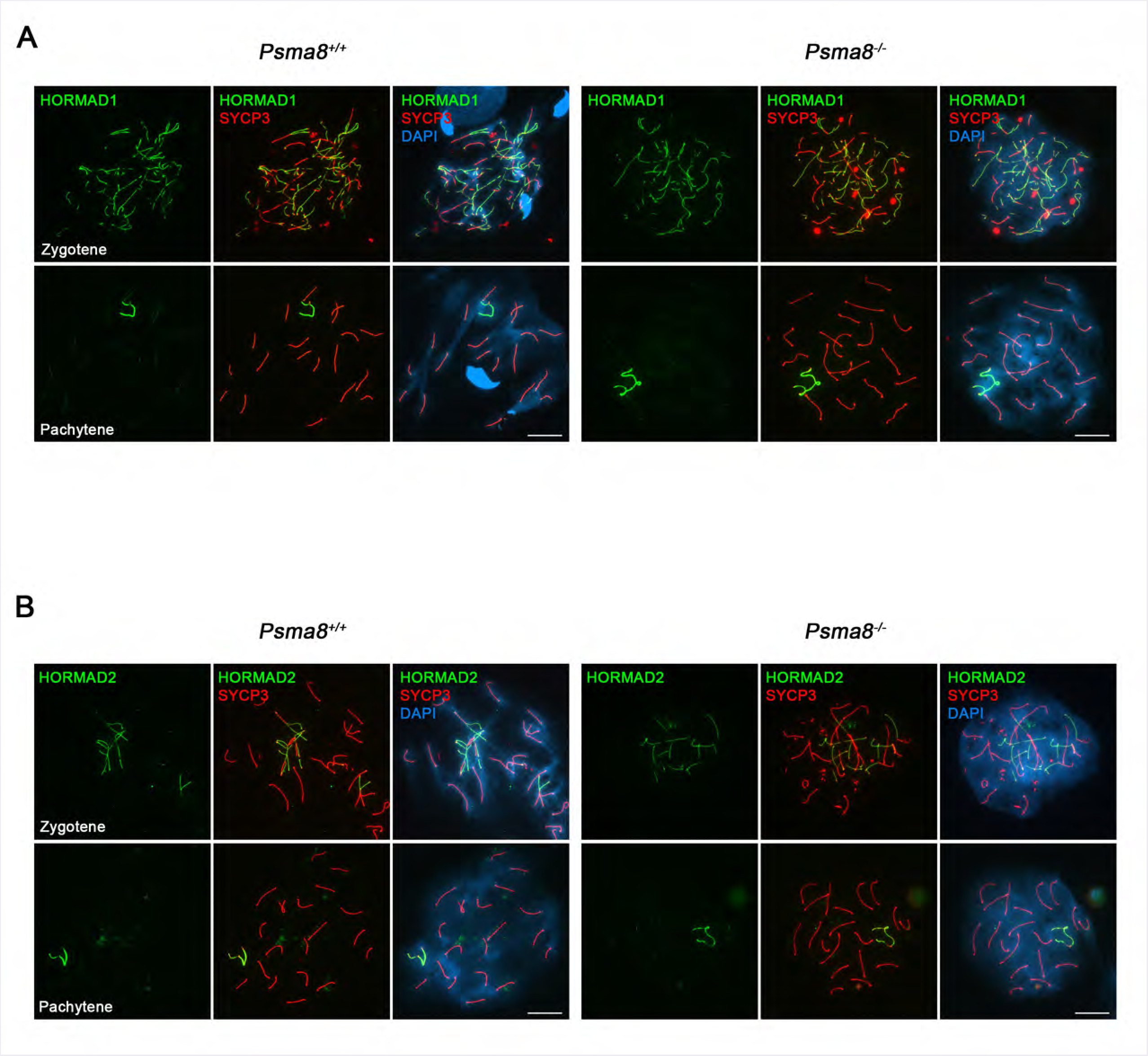
HORMADS are not affected by the increased expresion of *Trip13* in the *Psma8^−/−^* spermatocytes. Double immunolabelling of HORMAD 1 (A) and HORMAD2 (B) (green) with SYCP3 (red) in *Psma8*^+/+^ and *Psma8*^−/−^ spermatocytes at zygotene and pachytene stages. As synapsis progresses HORMAD1 and HORMAD2 are released from the AEs and maintained at the AE of the sex body similarly in the wild type and in the mutant spermatocytes. Bars represent 10 μm.

**Table EV1:**
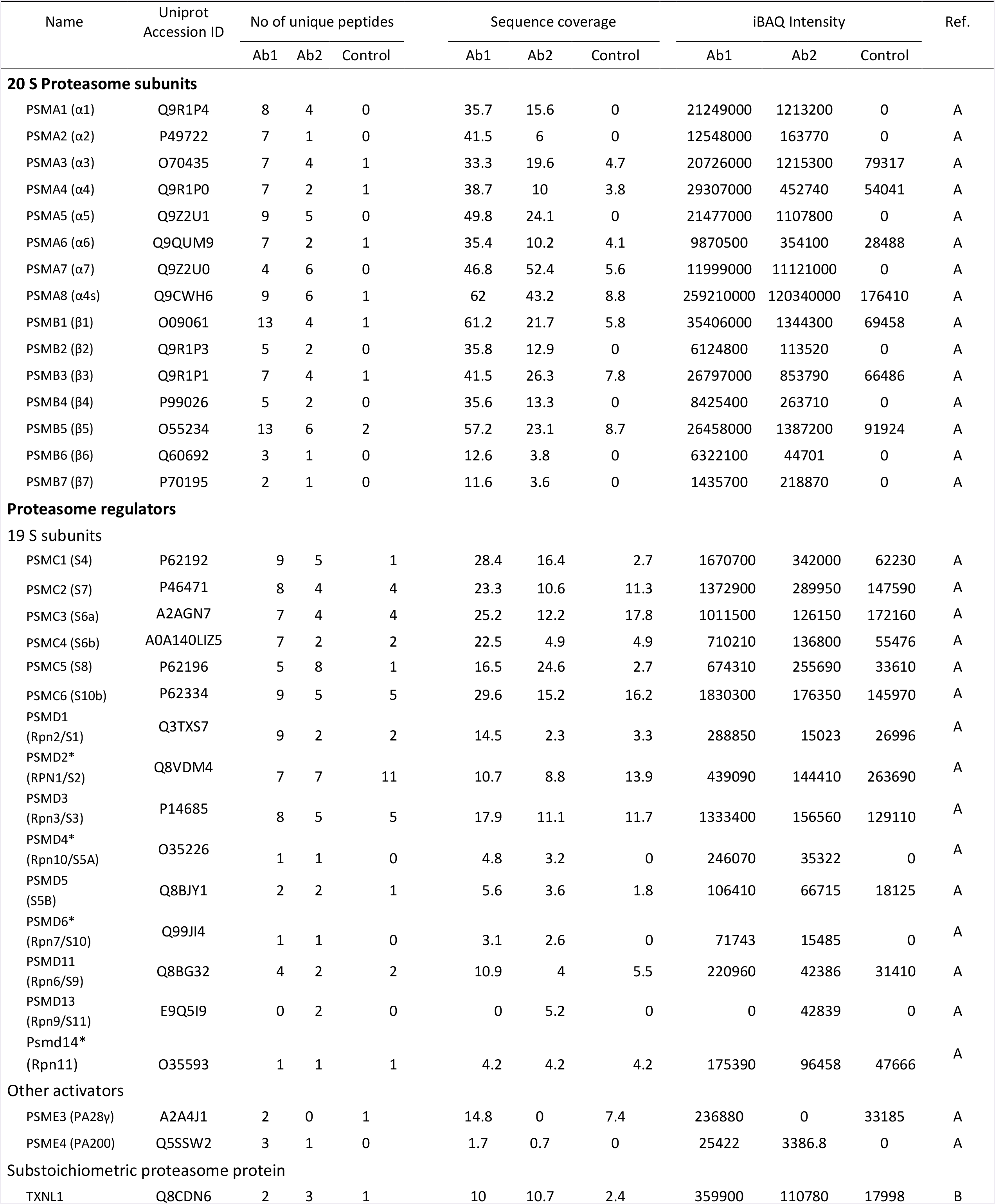
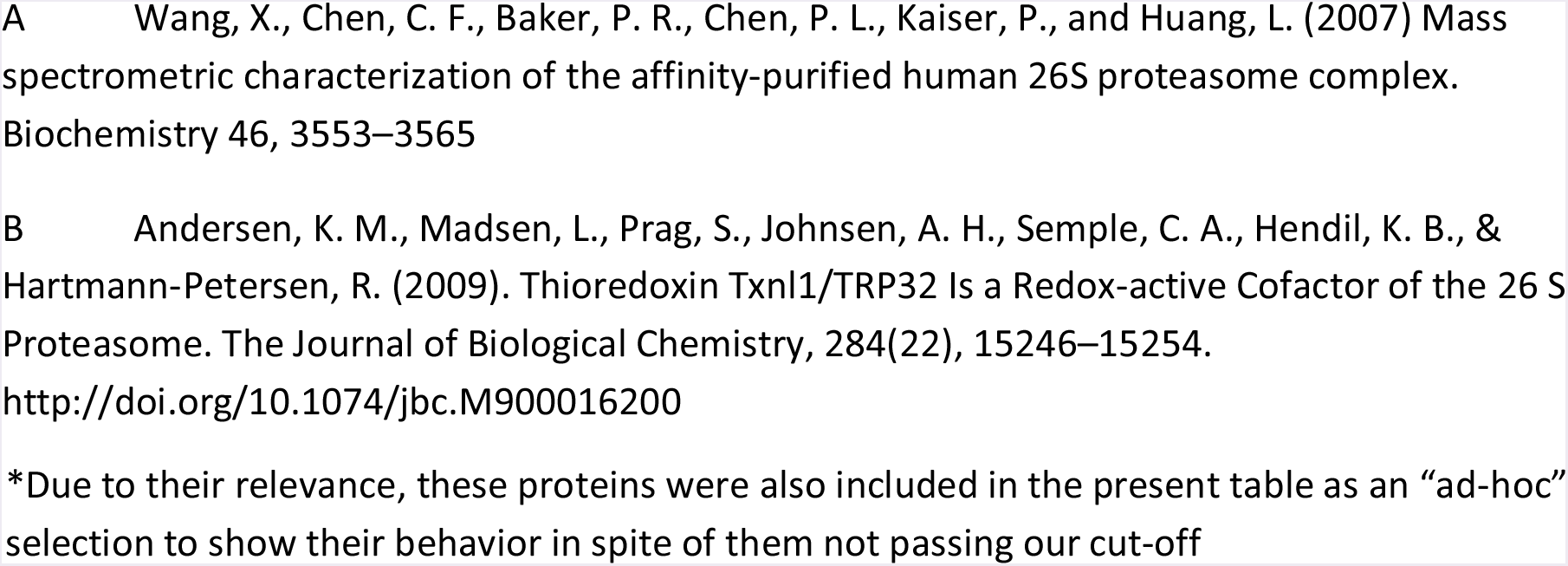
Proteasome subunits and proteasome regulators coimmuniprecipitated with PSMA8 selected after analysis and filtering of the data.

**Table EV2:** Selection of some of the proteasome-related proteins coimmuniprecipitated with PSMA8 selected after analysis and filtering of the data.

| Name | Uniprot<br>Accession ID | No of unique peptides |  |  | Sequence coverage |  |  | iBAQ Intensity |  |  |
| --- | --- | --- | --- | --- | --- | --- | --- | --- | --- | --- |
|  |  | Ab1 | Ab2 | Control | Ab1 | Ab2 | Control | Ab1 | Ab2 | Control |
| E3 ligases |  |  |  |  |  |  |  |  |  |  |
| CAND1 | Q6ZQ38 | 10 | 8 | 5 | 10 | 7.9 | 5.2 | 275300 | 143200 | 27601 |
| CCT2 | P80314 | 14 | 5 | 7 | 37.2 | 15 | 21.3 | 1713000 | 367550 | 184630 |
| CUL3 | Q9JLV5 | 3 | 6 | 1 | 6.8 | 9.6 | 1.3 | 104730 | 187950 | 6432.1 |
| CUL9 | E9QP09 | 3 | 2 | 0 | 1.1 | 0.6 | 0 | 20528 | 4934 | 0 |
| NEDD4 | P46935 | 2 | 0 | 0 | 4.7 | 0 | 0 | 68196 | 0 | 0 |
| RAD18 | E9Q392 | 3 | 0 | 0 | 43.7 | 0 | 0 | 20227000 | 0 | 0 |
| RBX1 | P62878 | 0 | 2 | 0 | 0 | 17.6 | 0 | 0 | 192900 | 0 |
| SKP1 | Q9WTX5 | 2 | 1 | 1 | 10.4 | 7.4 | 7.4 | 503960 | 136060 | 60179 |
| TRIP12 | A0A087WRV6 | 3 | 0 | 0 | 4.7 | 0 | 0 | 11882 | 0 | 0 |
| TRIM36 | E9Q3A0 | 5 | 1 | 0 | 9.3 | 2 | 0 | 237780 | 18278 | 0 |
| UBR5 | E9Q2H1 | 2 | 0 | 0 | 1.1 | 0 | 0 | 17518 | 0 | 0 |
| UFL1 | Q8CCJ3-1 | 2 | 0 | 0 | 3.8 | 0 | 0 | 57646 | 0 | 0 |
| ZC3HC1 | D3Z3D0 | 0 | 2 | 0 | 0 | 7.4 | 0 | 0 | 50351 | 0 |
| Deubiquitinases |  |  |  |  |  |  |  |  |  |  |
| USP5 | Q3U4W8 | 1 | 2 | 0 | 1.9 | 4.1 | 0 | 3715.8 | 28815 | 0 |
| USP7 | F8VPX1 | 5 | 0 | 0 | 5.7 | 0 | 0 | 105410 | 0 | 0 |
| USP9X | Q4FE56 | 10 | 2 | 0 | 4.9 | 1 | 0 | 121220 | 7683.3 | 0 |
| USP14* | E9PYI8 | 0 | 1 | 0 | 0 | 5.2 | 0 | 0 | 32751 | 0 |
| USP34 | F6WJB7 | 6 | 0 | 0 | 2 | 0 | 0 | 41209 | 0 | 0 |
| USP40* | Q8BWR4-3 | 1 | 1 | 0 | 1 | 1 | 0 | 22401 | 0 | 0 |
| USP47 | A0A1L1SV73 | 2 | 1 | 0 | 2.5 | 1.2 | 0 | 18169 | 2177.4 | 0 |
| Chaperones |  |  |  |  |  |  |  |  |  |  |
| AHSA1 | Q8BK64 | 2 | 1 | 1 | 8.6 | 3 | 6.5 | 227160 | 2825.1 | 20386 |
| CCT6B | Q61390 | 3 | 1 | 0 | 12.1 | 4.9 | 3.2 | 132900 | 25309 | 0 |
| DNAJA1 | P63037 | 2 | 1 | 0 | 8.1 | 3 | 0 | 1047200 | 67368 | 0 |
| DNAJB9 | Q9QYI6 | 1 | 2 | 0 | 4.5 | 11.3 | 0 | 66228 | 60676 | 0 |
| DNAJC7 | Q9QYI3 | 13 | 8 | 2 | 31.6 | 18.2 | 4 | 1376900 | 227200 | 25321 |
| HSP90B1 | P08113 | 17 | 8 | 9 | 28.8 | 12.2 | 14.5 | 4511300 | 1355800 | 619050 |
| HSP90AB1 | P11499 | 8 | 5 | 5 | 23.9 | 19.8 | 19.6 | 3171500 | 1127200 | 452050 |
| HSPBP1 | A0A0U1RPF2 | 2 | 0 | 0 | 6.9 | 0 | 0 | 46940 | 0 | 0 |
| PSMG1<br>(PAC1) | Q9JK23 | 2 | 1 | 1 | 5.9 | 3.5 | 3.5 | 258480 | 35351 | 19395 |
| TRAP1 | Q9CQN1 | 4 | 5 | 0 | 6.9 | 8.1 | 0 | 143070 | 57267 | 0 |
| Putative pips/ a priori unrelated / spermatogenesis related |  |  |  |  |  |  |  |  |  |  |
| ADAD1 | F8WI80 | 3 | 0 | 0 | 10 | 0 | 0 | 202110 | 0 | 0 |
| BOLL | G3UYE8 | 3 | 1 | 1 | 11.7 | 3.2 | 3.2 | 400070 | 23474 | 37352 |
| CAP1 | P40124 | 0 | 2 | 2 | 0 | 4 | 6.8 | 0 | 46407 | 28930 |
| CDK1 | P11440 | 1 | 1 | 0 | 8.1 | 6.4 | 0 | 7093.3 | 41663 | 0 |
| CDK5 | P49615 | 3 | 1 | 2 | 12 | 6.5 | 6.5 | 542280 | 197000 | 29690 |
| CDK16 | Q04735-2 | 0 | 2 | 0 | 1.7 | 6.3 | 0 | 0 | 25675 | 0 |
| DAZL | Q64368 | 3 | 3 | 0 | 13.4 | 15.4 | 0 | 473450 | 206150 | 0 |
| MAEL | A0A0A6YWQ9 | 2 | 0 | 0 | 4.8 | 0 | 0 | 247850 | 0 | 0 |
| RNF17 | Q99MV7 | 8 | 4 | 1 | 5.7 | 2.9 | 0.7 | 161450 | 29164 | 4053.3 |
| SHCBP1L | Q3TTP0 | 13 | 5 | 4 | 24.4 | 11 | 9.5 | 6787500 | 299050 | 173530 |
| SMC4 | E9Q2X6 | 4 | 1 | 0 | 3.6 | 0.7 | 0 | 130810 | 9056.2 | 0 |
| SMC6 | Q924W5 | 2 | 0 | 0 | 2.3 | 0 | 0 | 17639 | 0 | 0 |
| SPAG1 | Q80ZX8-3 | 4 | 3 | 0 | 8.1 | 6.1 | 0 | 290980 | 30285 | 0 |
| SPATA5 | A0A0G2JFY0 | 9 | 2 | 0 | 17.5 | 3.1 | 0 | 239890 | 13235 | 0 |
| SPATA20 | Q80YT5 | 7 | 7 | 3 | 11.6 | 12.4 | 4.1 | 1291900 | 234350 | 63556 |
| SYCP1 | Q62209 | 11 | 0 | 0 | 13.7 | 0 | 0 | 727550 | 0 | 0 |
| TDRD1 | Q99MV1 | 4 | 1 | 0 | 4.2 | 0.9 | 0 | 60474 | 3896.1 | 0 |
| TDRD6 | F2Z429 | 18 | 8 | 3 | 11.3 | 5 | 1.6 | 586920 | 46319 | 10974 |
| TDRD9 | Q14BI7 | 8 | 1 | 0 | 7.4 | 0.8 | 0 | 133480 | 5407.1 | 0 |
| TDRKH | A0A0G2JFB2 | 4 | 2 | 0 | 9.4 | 4 | 0 | 384790 | 71610 | 0 |
| TRIP13 | Q3UA06 | 2 | 2 | 0 | 5.3 | 5.3 | 0 | 98523 | 41275 | 0 |

\*Due to their relevance, these proteins were also included in the present table as an “ad-hoc” selection to show their behavior in spite of them not passing our cut-off.

**Table EV3:** **Quantification of the percentage of metaphases I and metaphases II in the seminiferous tubules** (number of metaphases relative to the number of prophases).

|  | **MI - AI** |  | **MII - AII** |  |
| --- | --- | --- | --- | --- |
|  | **WT** | **KO** | **WT** | **KO** |
| **Total** | 1.66 ± 0.50 | 6.75 ± 0.52 | 0.92 ± 1.03 | 8.32 ± 3.35 |
| **Apoptotic** | 0.09 ± 0.13 | 3.90 ± 0.53 | 0.00 ± 0.00 | 6.17 ± 4.05 |
| **n** | 855 | 1028 | 855 | 1028 |

**Table EV4.**
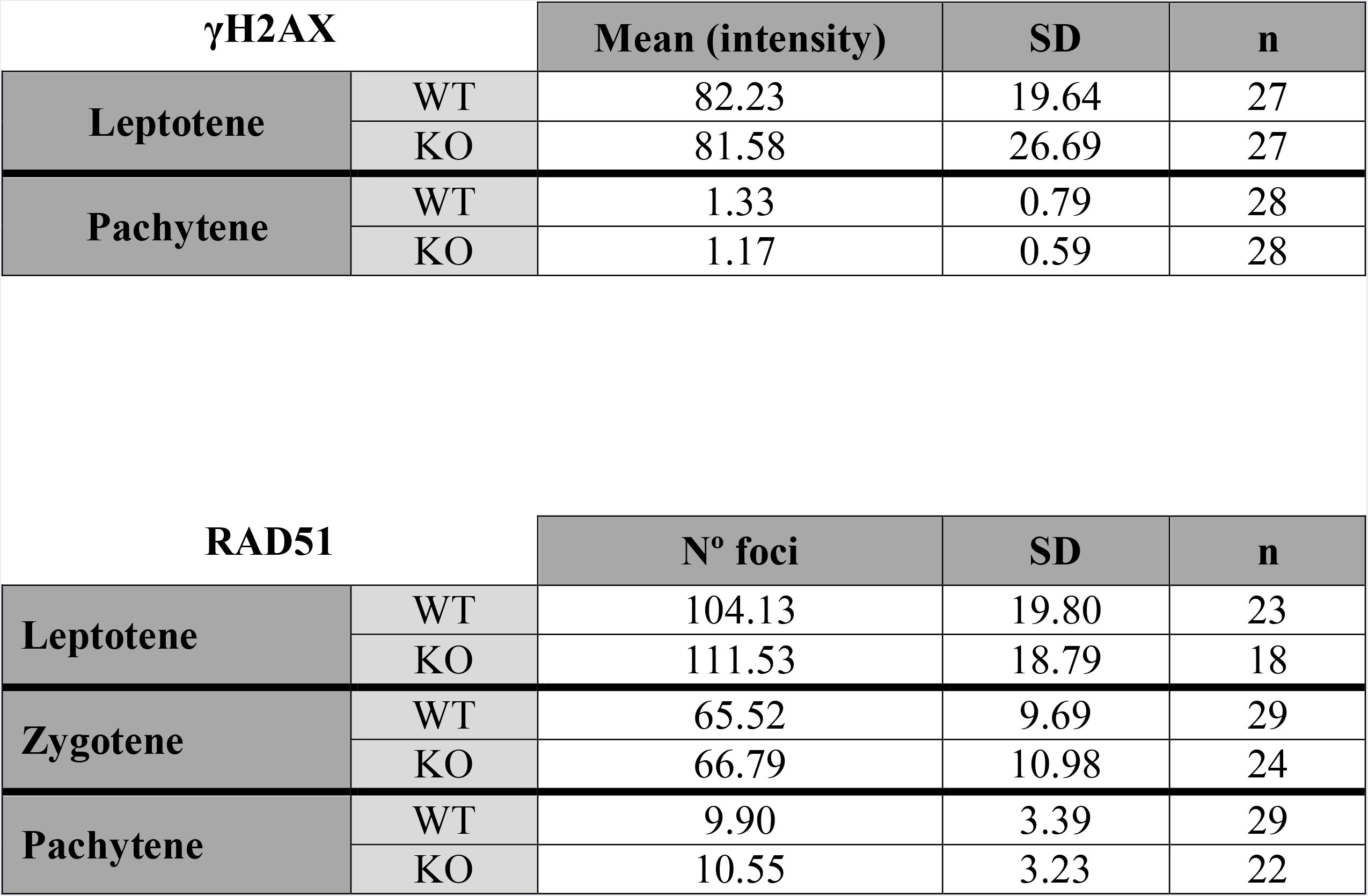
Quantification of γH2AX levels and RAD51 foci

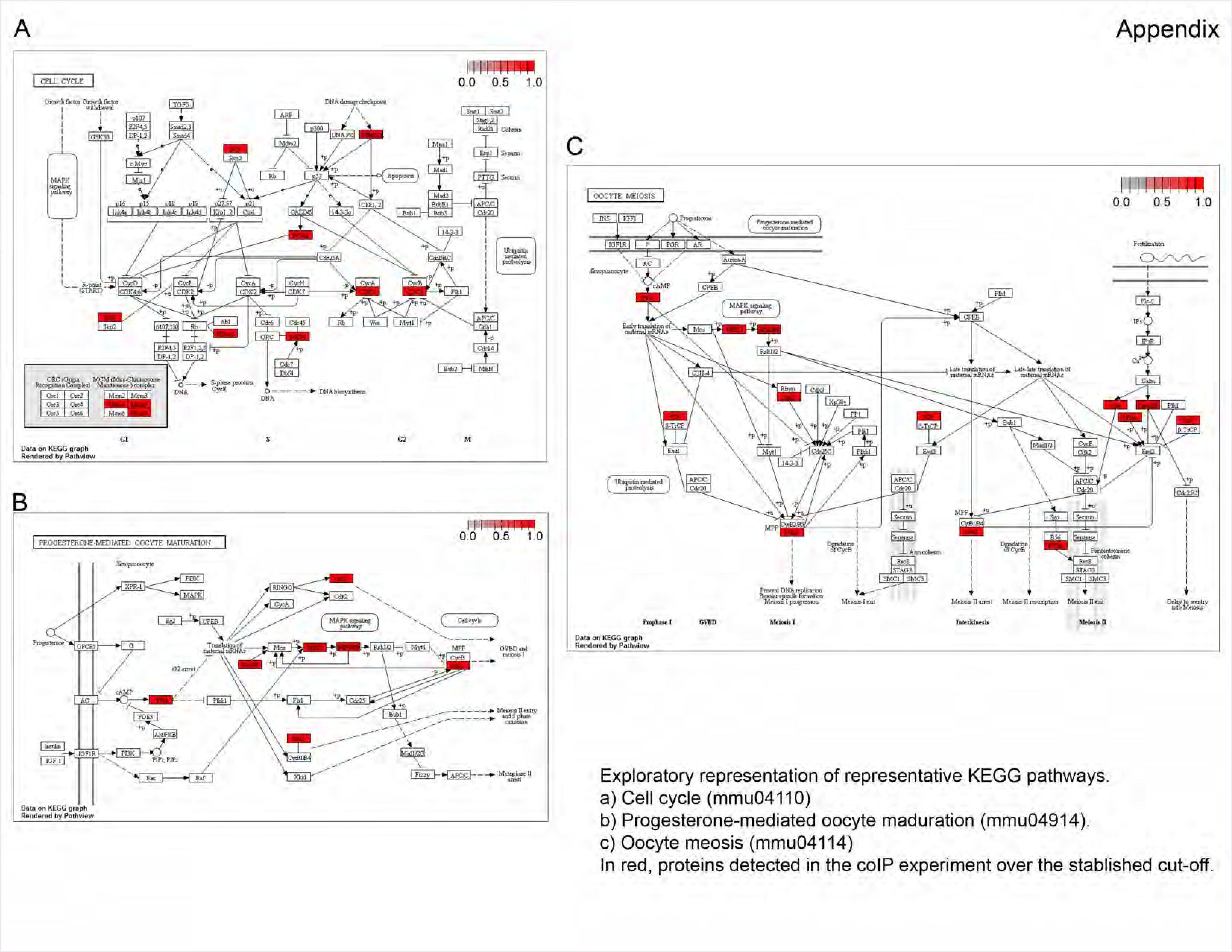
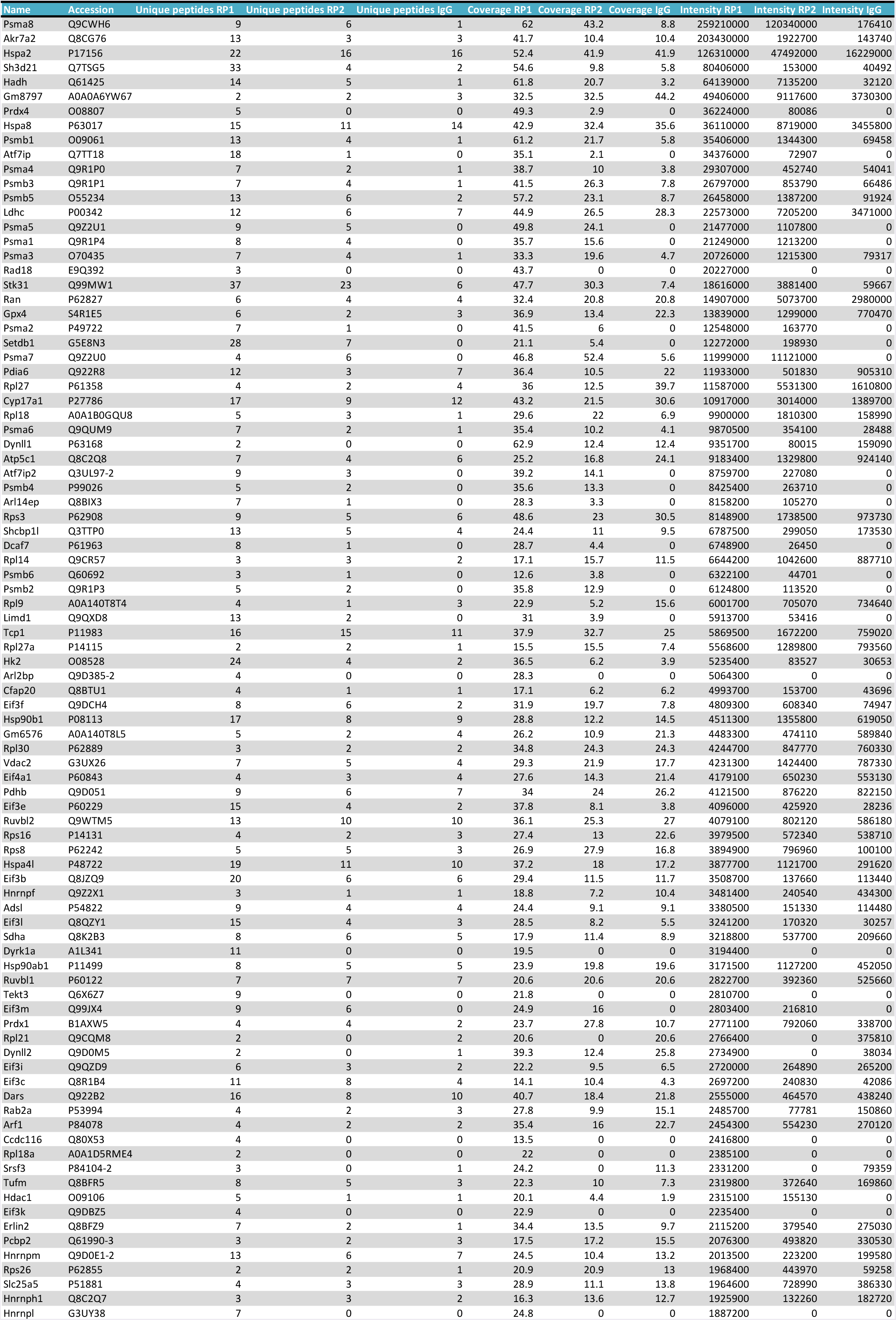
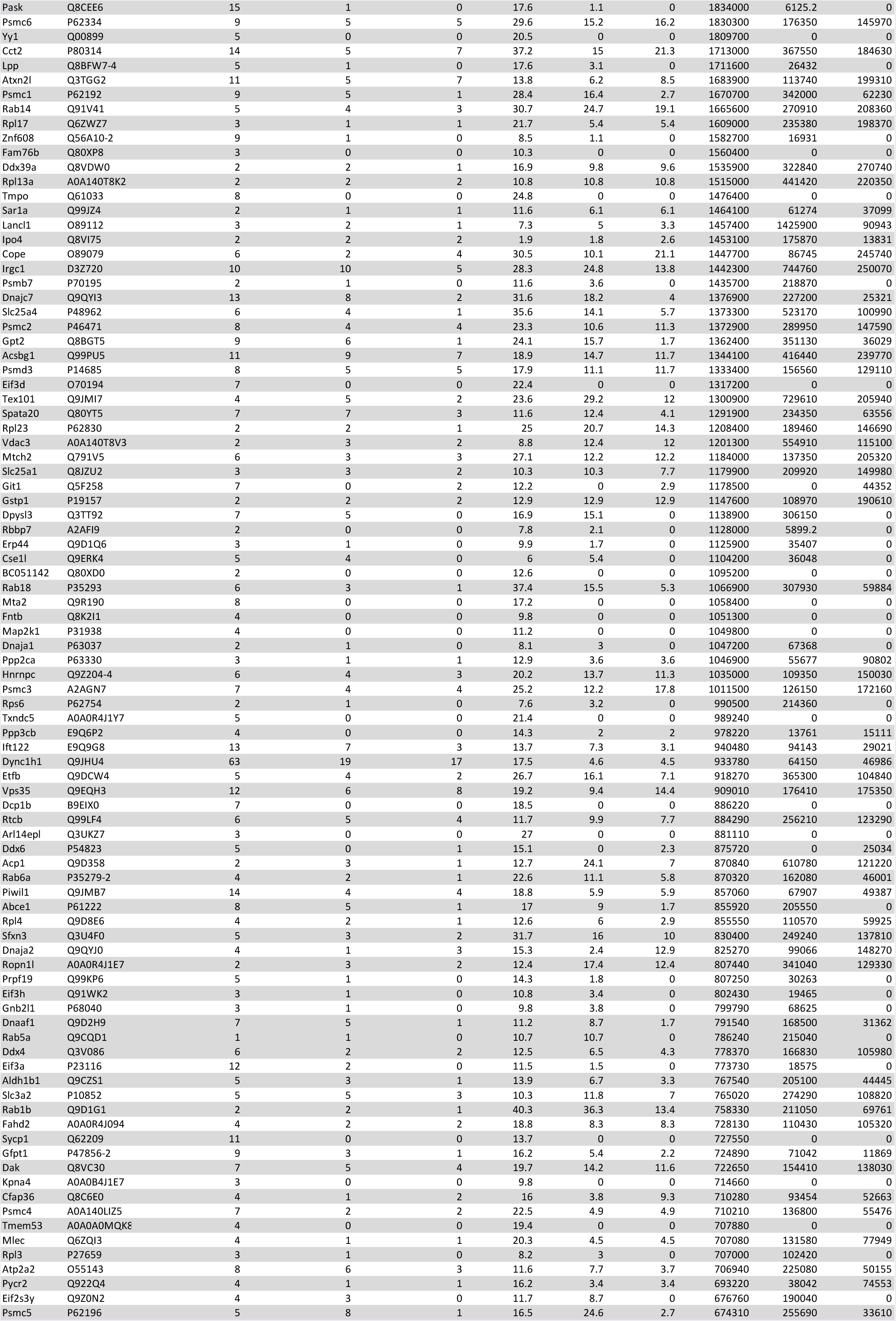
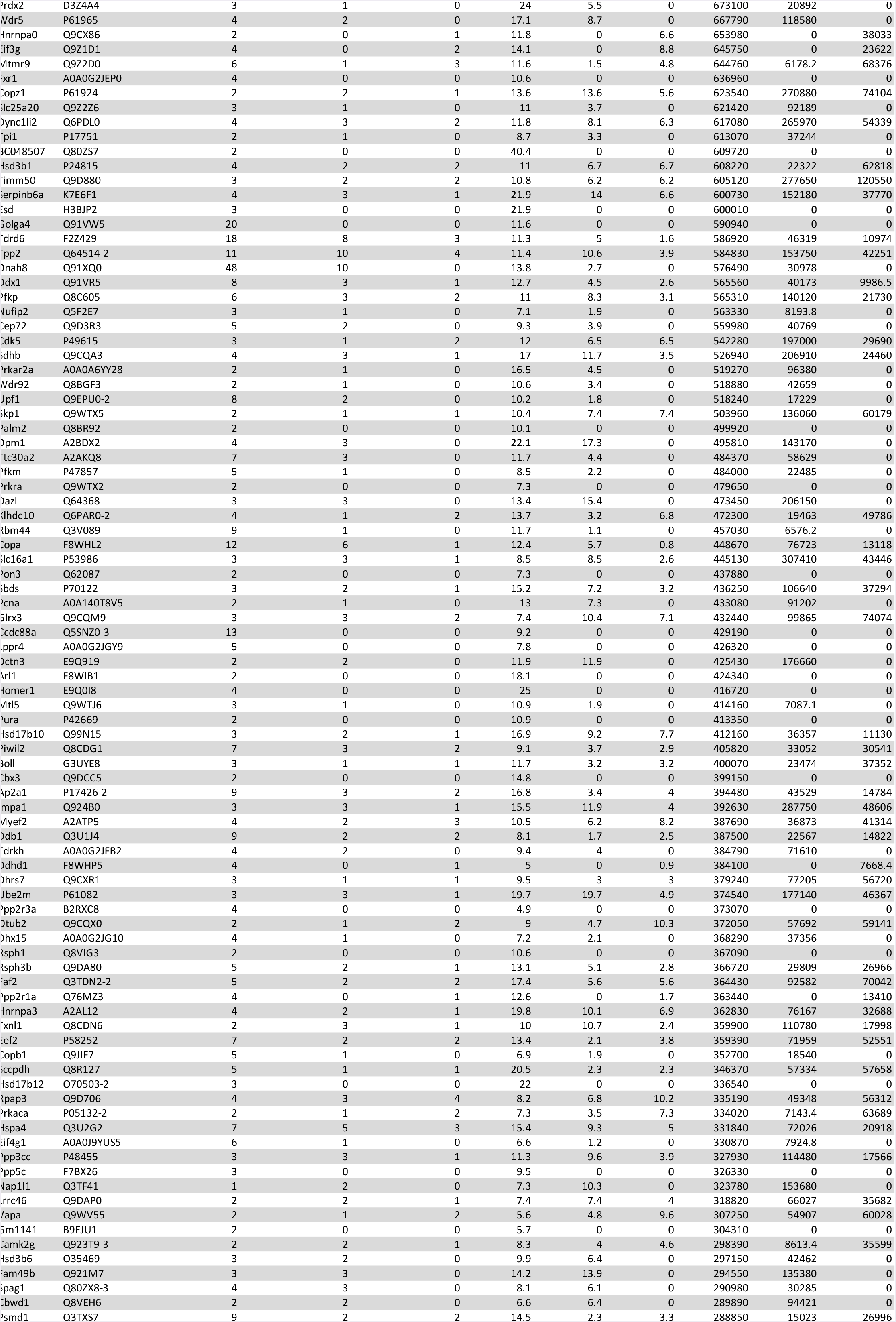
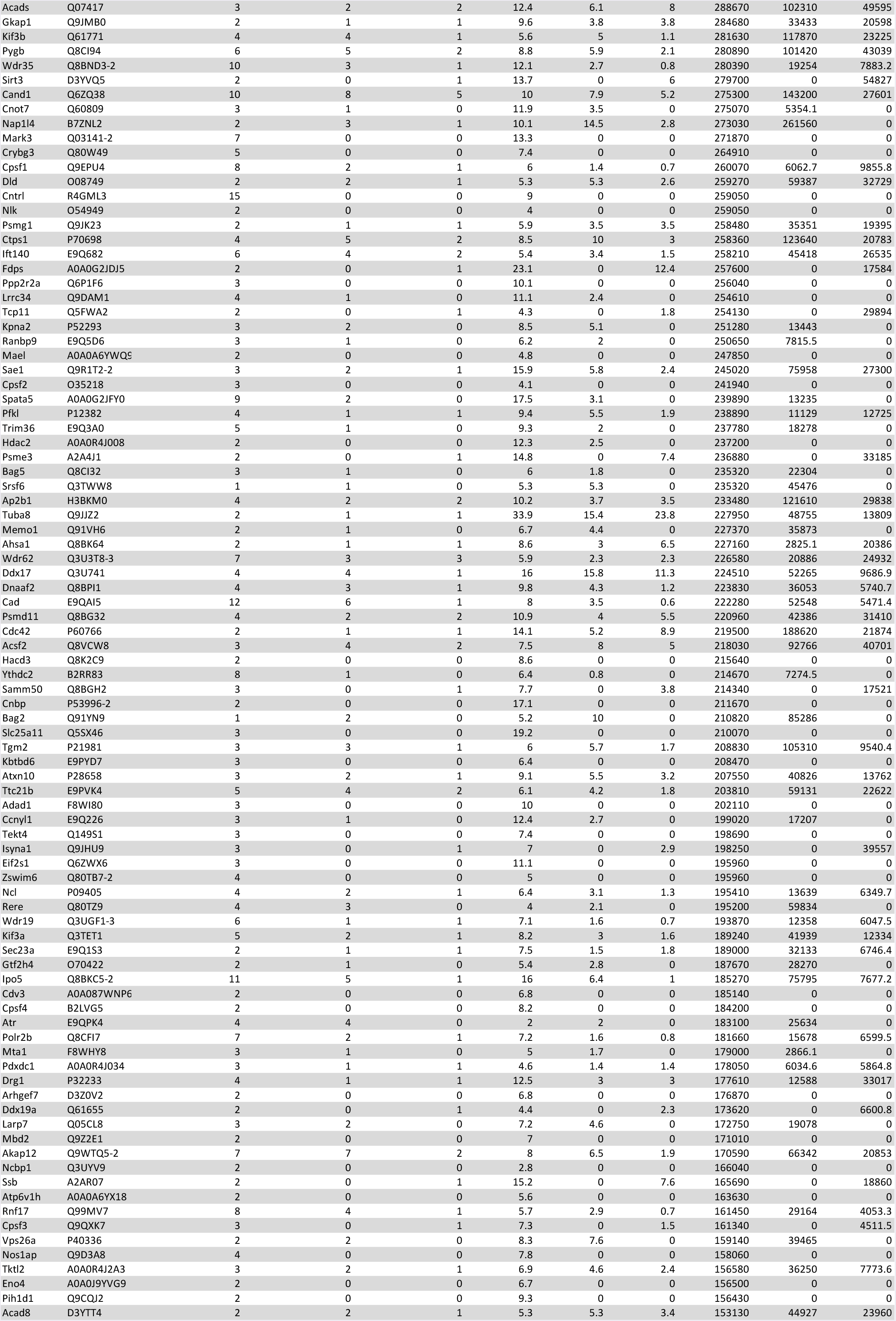
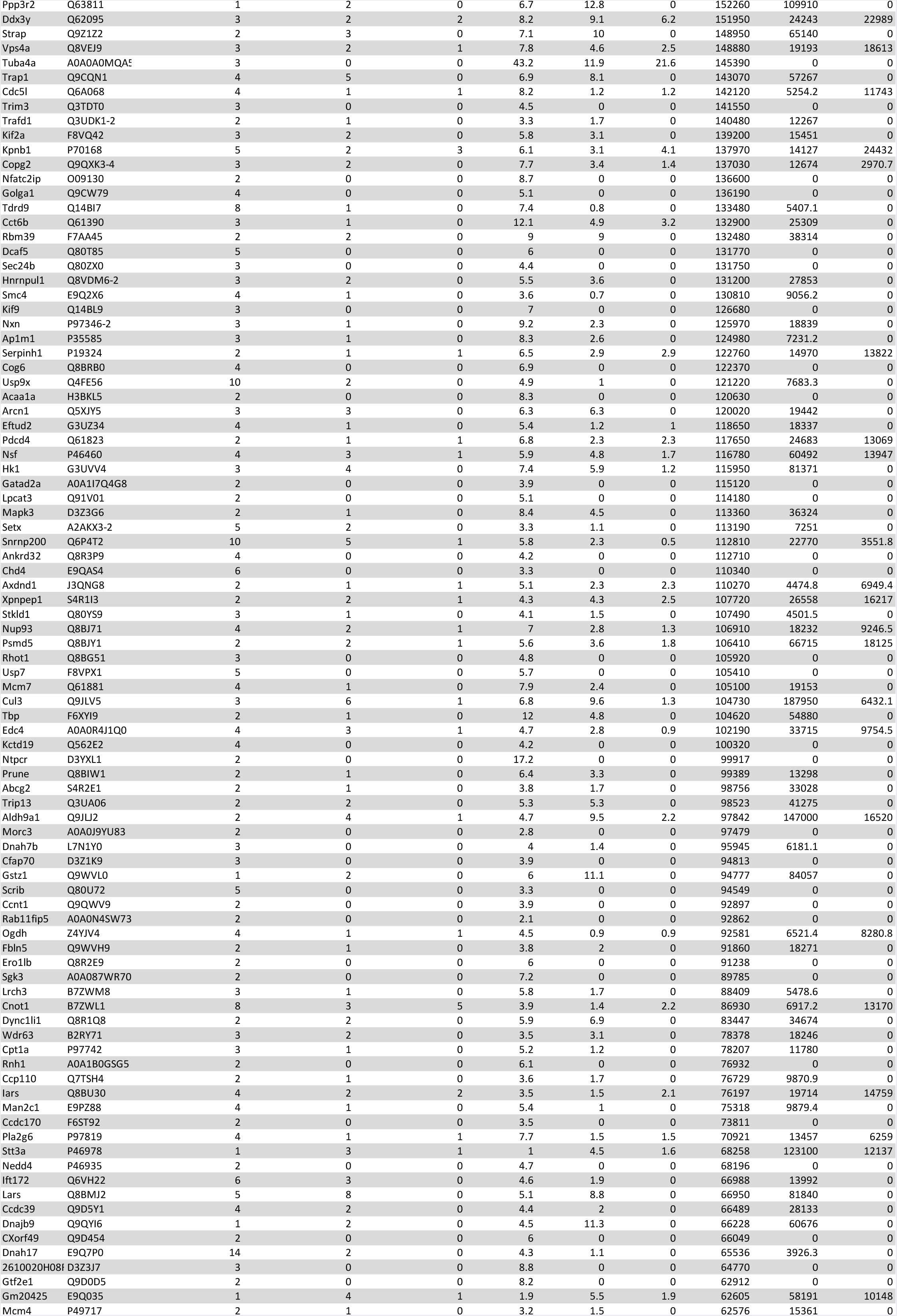
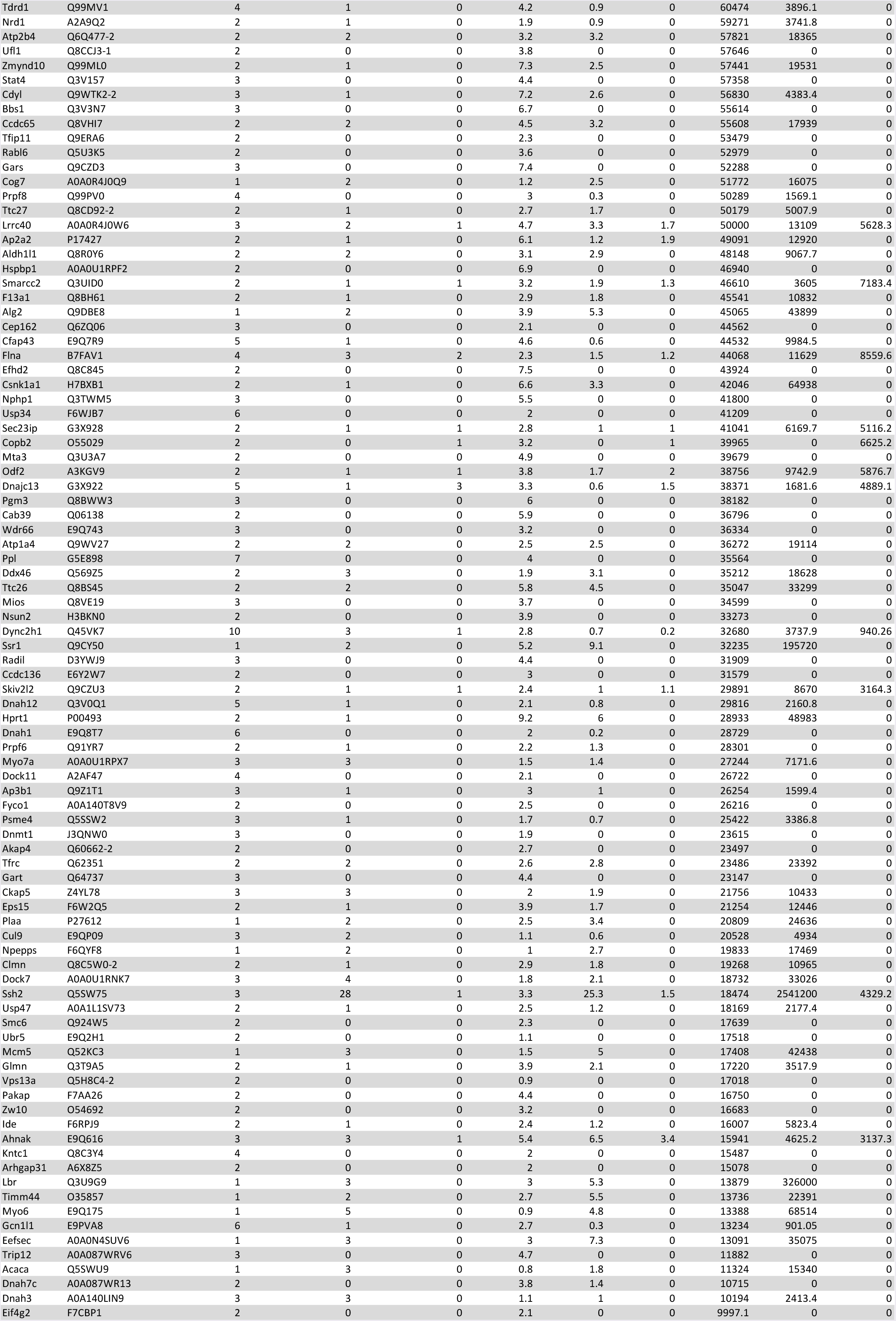
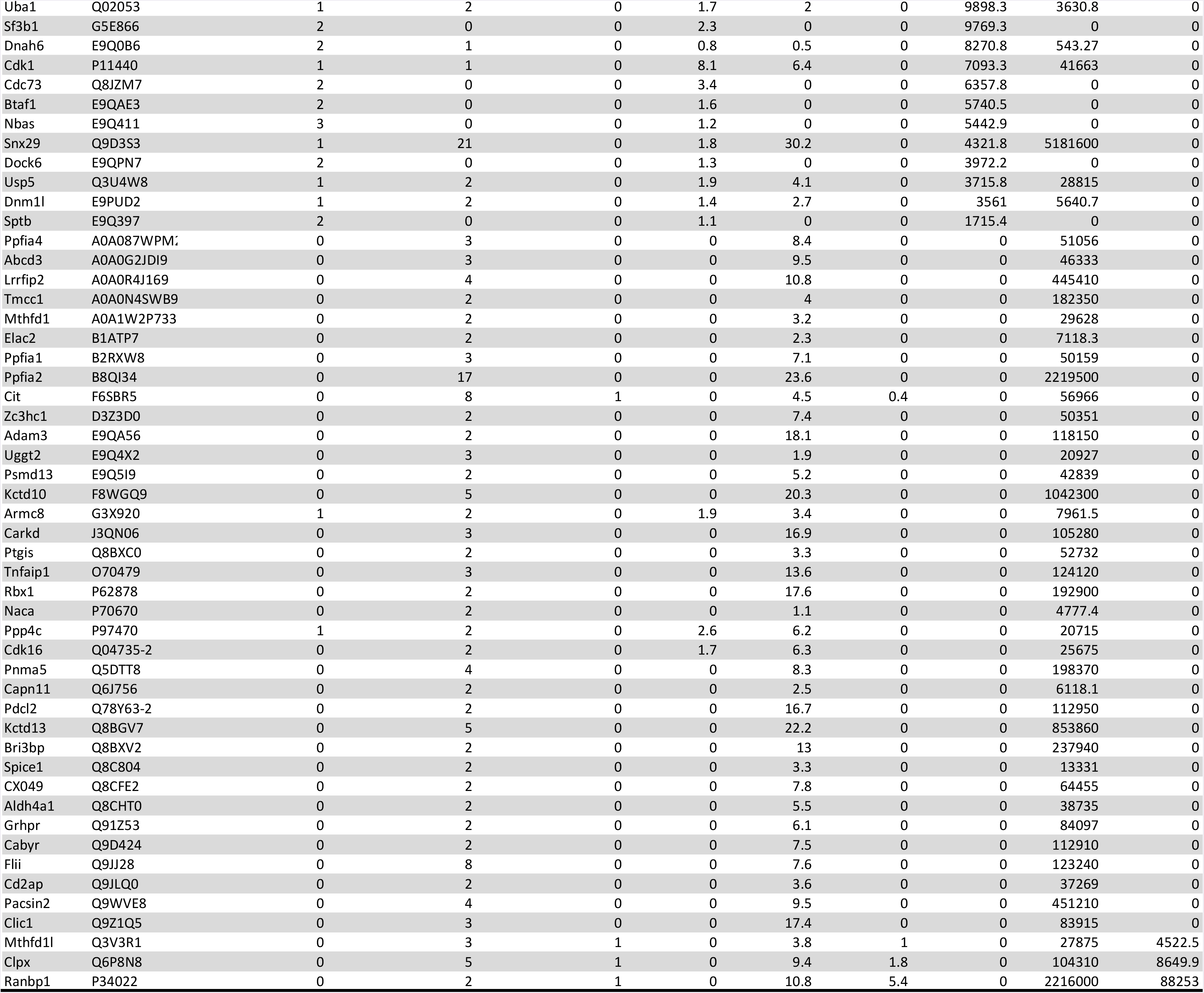

